# Genomic alterations drive brain metastases formation in colorectal cancer: The role of IRS2

**DOI:** 10.1101/2022.06.10.495658

**Authors:** Inbal Greenberg, Ethan Sokol, Anat Klein Goldberg, Dor Simkin, Moshe Benhamou, Shai Bar-Shira, Michal Raz, Rachel Grossman, Eilam Yeini, Paula Ofek, Bertrand Liang, Ronit Satchi-Fainaro, Hadas Reuveni, Tami Rubinek, Ido Wolf

## Abstract

Colorectal cancer (CRC) is currently the fourth leading etiology of brain metastasis (BM). Yet, mechanisms supporting the formation of CRC BM are mostly unknown. In order to identify drivers that lead to tropism and adaptation of CRC cells to the brain environment, we analyzed an extensive genomic database, consisting of over 36,000 human CRC primary and metastasis samples. Several genomic alterations specific for BM were noted, among them increased prevalence of insulin receptor substrate 2 (IRS2) amplification, observed in 7.6% of BM, compared to only 2.9% of primary tumors or other metastatic sites (p<7E-05). This observation was validated by Immunohistochemistry studies of human clinical samples, showing increased expression of IRS2 protein in BM. IRS2 is a cytoplasmic adaptor mediating effects of insulin and IGF-1 receptors and is involved in more aggressive behavior of different cancer types. In order to study the ability of IRS2 to promote growth of CRC cells under brain microenvironment conditions, we employed an in vitro system consisting of cultured human astrocytes or their conditioned media. Indeed, IRS2-overexpressed CRC cells survived better and formed larger 3D spheres when grown in brain-mimicking conditions, while IRS2-silenced CRC cells showed a mirror image. Similarly, In vivo studies, using intracranial CRC BM mouse model, demonstrated that IRS2-overexpressed cells generated larger brain lesions, while silencing IRS2 dramatically decreased tumor outgrowth and extended survival. Transcriptomic analysis revealed enrichment of oxidative phosphorylation (OXPHOS) and Wnt/β-catenin pathways by IRS2. Indeed, IRS2-expressing cells showed increased mitochondrial activity and glycolysis-independent viability. Furthermore, IRS2-expressing cells had increased β-catenin transcriptional activity, and either β-catenin inhibition or IRS2 inhibition in IRS2-expressing cells decreased their viability, β-catenin transcriptional activity, and mitochondrial activity. These data suggest involvement of IRS2 in modulating OXPHOS through β-catenin. Exploiting this mechanism as a potential vulnerability allowed us to develop novel treatment strategies against CRC BM. NT219 is a novel IRS1/2 inhibitor already being tested in clinical trials. Treatment of mice harboring CRC BM with NT219 and 5-flourouracil reduced tumor growth and prolonged mice survival. These data reveal, for the first time, the unique genomic profile of CRC BM and imply the IRS2 role in promoting CRC BM. These effects may be mediated, at least in part, by modulation of the β-catenin and OXPHOS pathway. These findings may pave the way for clinical trials evaluation this novel IRS2-based strategy for the treatment of CRC BM.

Accession no GSE203017 and here is the link: https://www.ncbi.nlm.nih.gov/geo/query/acc.cgi?acc=GSE203017

## INTRODUCTION

Colorectal cancer (CRC) is the third most common cancer worldwide [1]. Approximately 25% of CRC patients present with distant metastases at the time of diagnosis, and another 25% will suffer from metastases further on, with the liver as the main metastatic site [2]. With the advancement of treatments and prolongation of survival of CRC patients, the incidence of CRC brain metastases (BM) is rising and is currently the fourth leading cause of BM [3]. When BM appear, the disease is incurable, and the median survival of the patients is only three to six months [4]. Thus, the development of effective therapy for CRC BM is of utmost importance. The formation of metastases at a specific metastatic site is not a random process but rather requires adaptation of the cancer cells to the unique micro-environmental conditions of the hosting organ [5]. The brain environment is fundamentally different from that of the colon or other metastatic sites. Thus, the formation of BM requires not just the ability to penetrate through the blood-brain barrier (BBB) but also the ability to cope with low oxygen tension and limited availability of nutrients [6]. Identification and characterization of mechanisms mediating the adaptation of cancer cells to the brain microenvironment may expose novel vulnerabilities, which in turn can serve as potential targets for novel prevention and treatment strategies.

While the genomic landscape of primary CRC or CRC that metastasizes to the liver or lung is well established and includes mutations in the Wnt/β-catenin pathway, KRAS, and p53, genomic alterations (GA) leading to the development of CRC BM remain to be elucidated [7]. Other cancer types like lung, melanoma, and breast BM were characterized by GA, and alterations in the PI3K/AKT/mTOR pathway were common [8].

Insulin receptor substrate 2 (IRS2) is a cytoplasmic adaptor protein mediating the activity of insulin and IGF-1 receptors [9], thus enabling the activation of the downstream PI3K/AKT/mTOR and MAPK pathways [10]. IRS2 is expressed in various malignancies and has been reported to contribute to the development of unique tumor cell metabolism, as well as to tumor motility and invasion [11] [12]. Suppression of IRS2 has been shown to confer an inhibitory effect on the progression of breast, liver, and esophageal cancers, as well as of neuroblastoma [13][14][15][16]. In CRC, IRS2 mRNA and protein levels are positively correlated with progression from normal through adenoma to carcinoma, and deregulated IRS2 expression activates the PI3K/AKT/mTOR pathway and increases cell adhesion [17]. In addition, the frequency of IRS2 copy number gain is higher in CRC compared to other tumor types [18].

We hypothesized that specific GA promote the development of CRC BM, and identification of enriched GA may point to relevant mechanisms and pathways mediating tropism to the brain. To this aim, we analyzed a genomic database consisting of more than 35,000 CRC biopsies from local and metastatic sites. We discovered an increased prevalence of the IRS2 gene amplification in BM relative to local tumors and other metastatic sites. Using *in vitro* and *in vivo* studies, we revealed that IRS2 amplification facilitates CRC cells to grow under brain conditions. This phenotype is mediated by IRS2-dependent transcriptional and metabolic rewiring, supporting growth in the brain micro-environment. Our studies also indicate treatment of CRC cells with the IRS2 inhibitor NT219, in combination with the chemotherapy 5-fluorouracil (5-FU) as a potent suppressor of CRC BM development. Taken together, our data suggest a unique role for IRS2 amplification in facilitating brain tropism of CRC and imply a novel strategy for the treatment of patients harboring CRC BM.

## MATERIALS AND METHODS

### Clinical sample analysis

Comprehensive genomic profiling (CGP) was carried out in a Clinical Laboratory Improvement Amendments (CLIA)-certified, CAP (College of American Pathologists)-accredited laboratory (Foundation Medicine Inc., Cambridge, MA, USA) on CRC all-comers during the course of routine clinical care. Approval for the study was obtained from the Western Institutional Review Board (Protocol No. 20152817). Hybrid capture was carried out on exons from up to 395 cancer-related genes (FoundationOne v3: 323; FoundationOne v5: 395) and select introns from up to 31 genes frequently rearranged in cancer (FoundationOne v3: 24; FoundationOne v5: 31) (Tables S1 and S2). We assessed all known and likely pathogenic alterations across all classes of genomic alterations (GA) including short variant, copy number, and rearrangement alterations, as described previously [19]. Briefly, base substitutions were detected using a Bayesian method allowing the detection of somatic mutations at low MAF with increased sensitivity for mutations at hotspot sites. Indels were identified using a de Bruijn approach. Copy number events were detected by fitting a statistical copy-number model to normalized coverage and allele frequencies at all exons and ~3,500 genome-wide, single-nucleotide polymorphisms. Rearrangements were detected through an analysis of chimeric reads. Tumor mutational burden (TMB) was determined on 0.8–1.1 Mb as described previously [20]. At the time of the analysis, the dataset consisted of 20,858 local-biopsied and 15,139 distantly metastatic CRC samples.

### Constructs

Detailed in Table S3.

### Cells and transfections

Cell lines were originally obtained from the American Type Culture Collection and authenticated with the DNA markers used by ATCC (Manassas, VA, USA). The human colorectal cancer cell lines HCT116, HT29, SW480, SW403, and LS513, and the human embryonic kidney cell line HEK293T were grown in Dulbecco’s Modified Eagle’s Medium (DMEM) containing 10% fetal bovine serum (FBS). Human Astrocytes (HA) were grown and maintained according to provider instructions (ScienCell Research Laboratories, CA, USA). All cells were grown at 37°C in a humidified 5% CO_2_ atmosphere. All transfections used Jet Pei (Polyplus Transfection, Illkirch, France).

### Chemicals

NT219 was kindly provided by TyrNovo (Rehovot, Israel). 5-FU was purchased from Adooq Bioscience (Irvine, CA). ICG-001 and Alpelisib (BYL719) were purchased from Selleck Chemicals (Houston, TX). AKT1/2 kinase inhibitor was purchased from Sigma-Aldrich (St. Louis, MO, USA). IGF-1 was purchased from PeproTech Inc (Rocky Hill, NJ, USA).

### HA conditioned media (CM)

HA were grown to confluency in T75 Flasks, washed three times in PBS to remove any remaining growth medium and incubated in fresh serum-free astrocytes medium (SFM) for an additional 24 h. The CM of HA was collected, filtered in 0.2 µm (Millipore, USA), and used fresh for each experiment. SFM was used as control.

### Generation of stable cells

#### Generation of IRS2 overexpressed cells

pReceiver-Lv247-IRS2 or pReceiver-Lv247-Empty control vectors were transfected along with pCMV-VSV-G and psPax2 (2 µg: 2 µg: 2 µg) into HEK-293T cells using PEI reagent. CM was harvested twice, 48 and 72 h after transfection, filtered through 0.45 μm filters, and media supplemented with polybrene (8 μg/ml) was used to infect HCT116, HT29 and SW480 cells for 8 h. Stably infected cells were selected by puromycin (0.75 µg/ml). Two weeks later, single colonies were isolated and assessed for IRS2 mRNA and protein expression (Fig. S1). Two clones where IRS2 overexpression was most efficient were cultured routinely separately and mixed prior to each experiment.

#### Generation of IRS2 silenced cells

HEK-293T cells were transfected with shRNAs targeting IRS2 or nonspecific (NS) vectors along with pCMV-VSV-G and psPax2, as described above. For IRS2 silencing, SW403 and LS513 cells were infected with the generated viral particles, as described above, and stably infected cells selected by puromycin (0.75 µg/ml) were assessed for IRS2 mRNA and protein expression (Fig. 1E, F).

**Figure 1:**
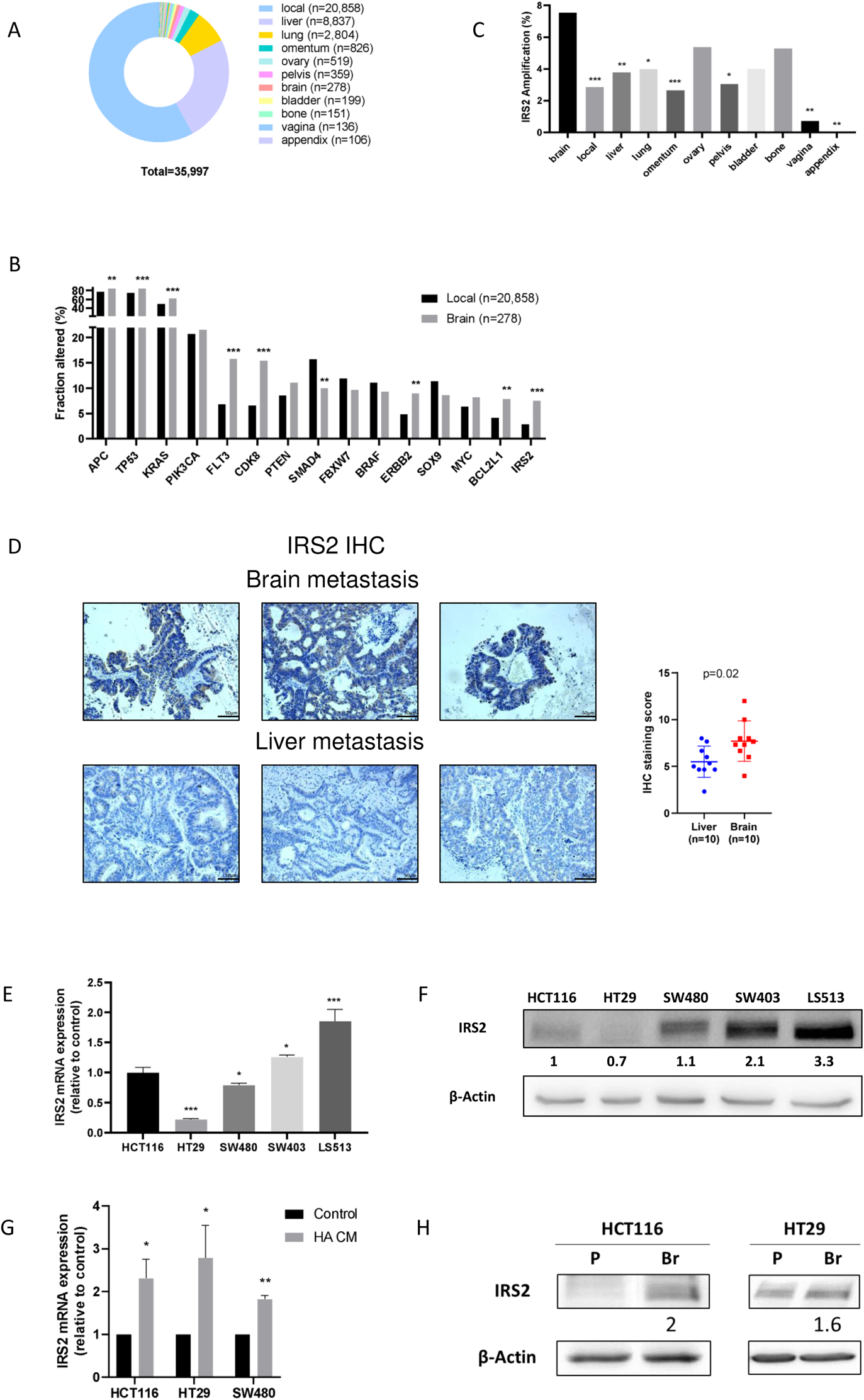
Increased prevalence of IRS2 amplification and expression in CRC brain metastasis. (A) Distribution of 35,997 CRC clinical samples, analyzed for genomic alterations, according to the biopsy site. (B) The most prevalent gene alterations in CRC local and BM are plotted. (C) IRS2 amplification rates in local CRC compared to different metastatic sites. (D) Representative IHC staining of IRS2 protein levels in CRC brain metastasis (n=10) and unmatched liver metastasis (n=10) and dot-plot of IHC data quantification by immuno-reactive score (IRS) (mean ± s.d., unpaired *t*-test). Scale bars represent 50µm. (E, F) Expression of IRS2 (E) mRNA and (F) protein in HCT116, HT29, SW480, SW403 and LS513 CRC cells determined by qRT-PCR and western blot, respectively. (G) HCT116, HT29, and SW480 cells were treated with HA CM or control media (SFM). IRS2 mRNA levels were determined by qRT-PCR and normalized to β-actin level. Statistical analysis was performed using one-way ANOVA (*, p<0.05, **, p<0.005, ***, p<0.0005). (H) Intracranial CRC BM mouse model was employed using HCT116 or HT29 cells. After tumor development cells were cultured and IRS2 protein levels were determined by western blot and normalized to β-Actin level. P: parental. Br: brain tumor-derived.

### Ethical statement

Mice were maintained at the SPF facilities of the Tel Aviv University. All experiments involving animals were approved by the TAU Institutional Animal Care and Use Committee (approval 01-19-007). Six-week-old athymic nude male mice were used. Patient-derived CRC brain and liver metastases tissues were collected after written informed consent was obtained from the research subjects by the Tel Aviv Sourasky Medical Center, under an approved institutional review board (IRB) (0172-17-TLV). Approval for the Foundation Medicine dataset portion of the study was obtained from the Western Institutional Review Board (Protocol No. 20152817).

### Intracranial mouse model

Six-week-old male athymic nude mice (Envigo CRS, Israel) were anesthetized by ketamine (150 mg/kg) and xylazine (12 mg/kg) intraperitoneally (IP) and secured to a stereotaxic device while a 1-cm incision was made on the skull midline between the ears. A small hole was drilled 2 mm left, 0.5 mm anterior and 6 mm ventral to the bregma. HCT116^CON^ or HCT116^IRS2^ cells or LS513^sh-NS^ or LS513^sh-IRS2^ cells (2 × 10^5^ cells) were slowly inoculated, and the incision was closed by wound clips. Animals were monitored twice a week for general health and body weight. Tumor development was followed by MRI (T1 weighted with contrast agent; Bruker, Germany). Mice were euthanized when they lost 10% of body weight in a week, 20% of their initial weight, or when neurological symptoms appeared.

To evaluate the effect of NT219 and 5-FU treatment in the LS513 model, LS513^sh-NS^ cells (2 × 10^5^ cells) were stereotactically implanted into 6-week-old male athymic nude mice (Envigo CRS, Israel). Mice were treated IP twice a week with 5-FU (30 mg/kg) (n=9), NT219 (70 mg/kg) (n=9), combined treatment (n=8) or control vehicle (n=9; D5W). Tumor development was monitored by MRI (T1 weighted with contrast agent; Bruker, Germany). Mice were euthanized when they lost 10% of body weight in a week, 20% of their initial weight, or when neurological symptoms appeared.

### MR imaging

MR imaging was performed with a 7T/30 MR Biospec (Bruker, Germany) equipped with a gradient unit of 660mT/m. MRI scans were acquired using a cross coil setup including 86mm resonator and quadrature mouse head coil. Animals’ body temperature was maintained by circulating warm water and their respiration was monitored. Mice were placed in the magnet with the ears positioned at the isocenter. T1-weighted 2D images with rapid acquisition with relaxation enhancement (RARE) sequence were acquired as following: TR/TEeff =800/14.8 ms, RARE factor 4, matrix size 256 × 256, slice thickness 0.5 mm, field of view 1.5 cm, resolution 0.08 × 0.08 mm2, number of averages 12, duration 6min. To enhance the brain with MR contrast, Gd-DTPA (Magnevist, Bayer HealthCare Pharmaceuticals, Wayne, NJ, USA) was administered by IP injection (6.5 mmol/kg) 20 minutes prior to MRI. In cases of opening the BBB a contrast-enhancement on T1-weighted MRI is expected. Tumor volume was quantified using MRIcro software.

### Immunofluorescence

Mice were sacrificed, transcardially perfused with PBS, and then with 4% paraformaldehyde (PFA). Brains were removed and fixed with 4% PFA overnight. Tissues were then cryoprotected with 30% sucrose in PBS overnight at 4°C. Fixed tissues were embedded in OCT-Tissue Freezing Medium (Scigen Scientific, Gardena, CA, USA) and frozen on dry ice. The samples were cut into 10 μm thick sections, placed on X-tra adhesive slides (Leica Biosystems, Peterborough, UK), and stored at − 20 °C. Sections were pre-incubated with a blocking solution containing 0.5% Tween 20 (Sigma-Aldrich, USA), 1% bovine serum albumin (BSA; Sigma-Aldrich, USA), and 3% horse serum (Gibco by Life Technologies, New York, NY, USA) for 1 h and then incubated overnight at 4 °C with primary antibodies: rabbit anti-human IRS2 (ab134101; Abcam, Cambridge, UK) and mouse anti-human active β-catenin (05-665; Upstate Biotechnology, Lake Placid, NY, USA). The sections were incubated with Alexa Fluor 488 donkey anti-rabbit IgG and Alexa Fluor 633 goat anti-mouse IgG secondary antibodies for 1 h (1:200; Molecular Probes by Life Technologies, Waltham, MA, USA). Control slides were incubated with the secondary antibody alone. Stained sections were examined and photographed using an LSM 700 confocal microscope (Zeiss, Germany). At least three fields of each individual sample were imaged and quantified using ZEN software (Zeiss, Germany).

### RNA sequencing

SW403^sh-NS^ and SW403^sh-IRS2^ cells (5 × 10^5^ cells/well) were seeded in triplicate for 24 h. Total RNA was extracted using the High Pure RNA Isolation Kit (Roche, Mannheim, Germany). RNA sequencing (RNA-seq) was conducted at the Tel-Aviv University Genomics Research Unit and Bioinformatics Unit (Tel-Aviv, Israel). The libraries were prepared using NEBNext® Ultra™ II RNA Library Prep Kit for Illumina® (New England BioLabs® Inc., USA). For sequencing: briefly, 1000ng of total RNA was fragmented followed by reverse transcription and second strand cDNA synthesis. The double strand cDNA was subjected to end repair, A-base addition, adapter ligation and PCR amplification to create barcoded libraries. Libraries were evaluated by Qubit and TapeStation. Sequencing was conducted with NextSeq 500/550 v2.5 (Illumina, USA) at 75-cycles, Single Read kit. The output was ~21 million reads per sample.

Adaptors were identified and removed from the raw sequence reads using TagCleaner [21]. Trimmed reads were aligned to the human reference genome UCSC hg19 (Gencode gene annotations) using STAR 2.6.1a [22]. Mapping was followed by transcriptome-wide abundance quantification using Salmon [23]. Pre-alignment and post-alignment quality control (QC) reports were generated using MultiQC [24]. Genes that were differentially expressed between the comparison and control groups were characterized using R package DESeq2 [25] by identifying genes with |log2FC|>1 and a P-adjusted value <0.05. Pathway analysis of the differentially expressed genes was performed using the R package clusterProfiler [26].

### Immunohistochemistry

Immunohistochemistry (IHC) staining was performed on formalin-fixed paraffin-embedded CRC brain or liver metastasis tissue using a monoclonal anti-IRS2 antibody (ab84906; Abcam, Cambridge, UK). 3.5mm sections were mounted on X-tra Adhesive Precleaned Micro Slides (Leica, Richmond, USA) and processed by an automated immunostainer (VENTANA BenchMARK ULTRA, Ventana Medical System, Tucson, AZ, USA). Automated immunostaining was performed using the I-View DAB detection kit (Ventana Medical System), according to Ventana program. The I-View DAB detection kit utilizes biotinylated secondary antibodies to locate the bound primary antibody, followed by the binding of Streptavidin-HRP (horseradish peroxidase) conjugate. The complex was then visualized with hydrogen peroxidase substrate and 3, 3’-diaminobenzidine (DAB) tetrahydrochloride chromogen. The incubations were performed at a controlled temperature of 37°C. The sections were then counterstained with Gill’s hematoxylin, dehydrated, and mounted for microscopic examination. The samples were quantitatively analyzed using the immunoreactive score (IRS). IRS gives a range of 0–12 as a product of multiplication between positive cells proportion score (0– 4) and staining intensity score (0–3) [27].

### Seahorse analysis

Oxygen consumption rate (OCR) and extracellular acidification rate (ECAR) measurements were performed with a Seahorse XF 96 Analyzer from Seahorse Bioscience (Agilent, USA) according to the manufacturer’s instructions. Mitochondrial respiration was studied by monitoring OCR using Seahorse Cell Mito Stress Test kit from Seahorse Bioscience and glycolytic activity was studied by monitoring ECAR using Seahorse Glycolysis Stress Test Kit from Seahorse Bioscience. Cells were plated at a density of 1 × 10^4^ cells per well, 10 replicates for each treatment. On the day of analysis, media were changed to Seahorse XF Base Medium supplemented with glutamine (2 mM) for the glycolysis stress test, or pyruvate (1 mM), glutamine (2 mM), and glucose (10 mM) for the mito stress test followed by incubation at 37°C in a non-CO2 incubator for 1 h. Mitochondrial respiration was measured under basal conditions followed by the sequential addition of oligomycin (2 μM), FCCP (0.5 μM), rotenone (0.5 μM), and antimycin A (0.5 μM). Glycolysis activity was measured following sequential addition of glucose (10 mM), oligomycin (1 μM), and 2-deoxy-glucose (2-DG; 50 mM). After each injection, four time points were recorded with approximately 35 minutes between each injection. The OCR and ECAR were automatically recorded and calculated by the Seahorse XF-96 Software and normalized to cell number.

### Quantitative RT-PCR

Total RNA and cDNA synthesis was performed as previously described [28]. Primers were synthesized by IDT (Coralville, IA, USA) and are listed in Table S4. Quantitative RT-PCR (qRT-PCR) was used to determine mRNA level as previously described [28].

### Western blot

Cells were harvested, lysed, and the total protein was extracted as previously described [28]. Lysates were resolved on 10% SDS-PAGE and immunoblotted with the indicated antibodies (Table S5).

### Colony assay

Cells were cultured at low density for two weeks and proceed as previously described [28].

### Methylene blue assay

Cells were plated (5 × 10^3^ cells/well) and after 24h medium was changed to the appropriate media (*e.g.*, regular media or human astrocytes conditioned media (HA CM)) or control media (depending on the experiment) and incubated at 37°C, 5% CO_2_ for 24h. Methylene blue assay was performed as previously described [28] and normalized to viability before treatment.

### Invasion assay

Cells were plated (5 × 10^4^ cells/insert) into the upper side of Matrigel coated 24 transwell inserts, with pore sizes of 8 µm (Corning, New York, USA), in media without FBS, whereas the lower chamber contained media with 10% FBS. After 48 h, the upper side of the apical chamber was scraped gently with cotton swabs to remove non-invading cells, and invading cells were fixed and stained with crystal violet.

### Migration (Transwell) assay

Cells were plated as for invasion assay, only the inserts were not covered with Matrigel.

### 3D sphere formation assays

CRC cells were seeded (2 × 10^3^ cells/drop) in GravityPLUS plates (Insphero, Switzerland) and after four days, spheres were transferred to GravityTRAP plates. Viability of spheres was determined after seven days using Realtime-Glow MT assay (Promega, Wisconsin, USA) and spheres were photographed at the end of the experiment.

For CM experiments, CRC cells were seeded as above. After two days, spheres were transferred to GravityTRAP and medium was changed to HA CM or control media. Viability of spheres was determined after five days using Realtime-Glow MT assay and spheres were photographed at the end of the experiment.

For HA experiments, HA were seeded in GravityPLUS plates. After three days, spheres were transferred to GravityTRAP plates and IRS2-overexpressed or control empty vector cells expressing m-cherry were added. Cells were photographed and viability assessed by m-cherry quantification using IVIS Lumina III instrument (PerkinElmer, USA) after 24, 48 and 72 h.

### Luciferase assay

Cells were plated in 12-well plates (1 × 10^5^ cells/well) and transfected with either pTOPFLASH or pFOPFLASH. Forty-eight hours after transfection, luciferase assay was conducted using the Luciferase Assay System kit (Promega, CA, USA) according to the manufacturer’s instructions. In all assays, FOPFLASH activity was measured by replacing the pTOPFLASH with pFOPFLASH under equivalent conditions. Luciferase units were normalized to total protein concentration

### Drug interaction analysis

*The Bliss Independence method* was employed to determine drug interaction [29]. The model predicts that if the individual drugs have the inhibitory effects *f_1_* and *f_2_*then the expected combined effect of the two drugs is:

*E(f_12_) = 1 – (1 – f_1_) (1 - f_2_) = f_1_ + f_2_ – f_1_ f_2_*

Excess over Bliss (*eob*) is calculated by *eob = f_12_ - E(f_12_)*, where *f_12_* is the observed combined effect. A positive, negative, or null value, is used to determine a synergistic, antagonistic or no interaction, respectively.

### Statistical analysis

Statistical analysis was performed using GraphPad Prism version 8.0.0 for Windows. All information regarding statistical test performed, ‘n’ values and p-values per experiment can be found in the figure legends, figures and results. p values < 0.05 were considered statistically significant, with ∗ p < 0.05, ∗∗ p < 0.005, ∗∗∗ p < 0.0005, unless otherwise indicated in the figure.

## RESULTS

### Increased prevalence of IRS2 amplification in CRC brain metastasis

In order to decipher the molecular landscape of CRC BM, we analyzed the prevalence of GA in CRC clinical samples derived from local (n=20,858) and metastatic (n=15,139) samples in a large real-world database (Foundation Medicine Inc., FMI). Most of the metastatic samples were obtained from the liver (n=8,837; 58%), followed by lung (n=2,804; 19%), omentum (n=826; 5%), and brain (n=278; 2%) (Fig. 1A). Patients with BM biopsies were similar in age and sex to patients with biopsies obtained from other sites (median 60 vs. 60 respectively, p=0.16; fraction male 57.97% vs. 53.43% respectively, p=0.1). Median tumor mutation burden (TMB) was modestly higher in brain metastasis samples compared to other metastatic sites (5.0 vs. 3.8, respectively, p<3E-11) (Table 1). Comparison of BM to local tumor samples indicated a higher frequency of alterations in *IRS2, CDK8, FLT3, FGF14*, and *KRAS* (Fig. 1B) in the BM. Amplification of *IRS2* was noted in 2.9% of local tumors compared to 7.6% of brain metastases (p<7E-05) (Fig. 1B). Importantly, increased IRS2 amplification was unique to BM, with other metastatic sites showing prevalence similar or lower compared to the local site (Fig. 1C), suggesting a role for IRS2 in CRC brain tropism. Enrichment of IRS2 was noted only in CRC BM and not in BM originating from other tumors, including lung cancer (0.8%), breast cancer (2.6%), and melanoma (0.2%) (Table 2).

**Table 1:** Demographic properties of Foundation Medicine CRC population.

| Met location | Brain met | Other met | p-val |
| --- | --- | --- | --- |
| Median Age | 60 | 60 | 0.16 |
| Median TMB | 5.00 | 3.75 | 2.87E-11 |
| Fraction Male | 57.97% | 53.43% | 0.14 |
| Fraction EUR | 77.34% | 75.08% | 0.44 |
| Fraction AFR | 10.79% | 11.10% | 1.00 |
| Fraction EAS | 1.80% | 3.70% | 0.11 |
| Fraction SAS | 0.72% | 0.79% | 1.00 |
| Fraction AMR | 9.35% | 9.34% | 1.00 |

Met, metastasis; TMB, tumor mutational burden; AFR, African; EAS, East Asian; AMR, Admixed American; EUR European, SAS South Asian

**Table 2:** IRS2 amplification in brain metastasis compared to local tumors in different cancers types

| Disease | Brain met | Brain met with IRS2 amplification | % Brain met with IRS2 amplification | Local | Local with IRS2 amplification | % Local with IRS2 amplification | OR | corrected p-val |
| --- | --- | --- | --- | --- | --- | --- | --- | --- |
| NSCLC | 3224 | 26 | 0.8% | 33625 | 215 | 0.6% | 1.3 | 0.40 |
| Breast | 781 | 20 | 2.6% | 11277 | 171 | 1.5% | 1.7 | 0.14 |
| Melanoma | 538 | 1 | 0.2% | 3002 | 8 | 0.3% | 0.7 | 1.00 |
| CRC | 278 | 21 | 7.6% | 20858 | 598 | 2.9% | 2.8 | 0.00 |
| Esophagus | 157 | 3 | 1.9% | 6210 | 46 | 0.7% | 2.6 | 0.32 |
| Kidney | 132 | 0 | 0.0% | 2733 | 11 | 0.4% | 0.0 | 1.00 |
| Ovary | 116 | 0 | 0.0% | 5815 | 34 | 0.6% | 0.0 | 1.00 |
| Small cell | 100 | 0 | 0.0% | 1325 | 36 | 2.7% | 0.0 | 0.35 |

*Met, metastasis; CRC, colorectal cancer; NSCLC, Non-small cell lung cancer*

In order to validate these data and corroborate the association between the genomic data and protein expression, we examined the expression of the IRS2 protein (which correlates with *IRS2* gene copy number and mRNA level [18]) using IHC staining in human CRC BM and liver metastasis (LM) samples (n=10 for each tumor type) and noted increased IRS2 expression in the BM compared to the LM samples (p=0.02) (Fig. 1D).

### Brain environment induces IRS2 expression in CRC cells

We hypothesized that if indeed IRS2 confers a growth advantage to CRC cells grown in the brain environment, exposure of CRC cells that do not normally express IRS2 to this environment will induce IRS2 expression. In order to test this hypothesis, we first characterized IRS2 mRNA and protein levels in three IRS2 non-amplified CRC cell lines (HCT116, HT29, and SW480) and two IRS2-amplified CRC cell lines (SW403 and LS513) [18]. As expected, a correlation between IRS2 mRNA (Fig. 1E) and protein (Fig. 1F) expression and IRS2 copy number was observed. In order to test the effect of brain environment on IRS2 expression, HCT116, HT29, and SW480 cells that do not normally express IRS2 or have low expression, were grown in HA CM, and IRS2 mRNA levels were examined. Indeed, IRS2 expression was significantly upregulated under HA CM in each cell line (Fig. 1G). In order to test this effect *in vivo*, HCT116 and HT29 cells were injected into brains of nude mice. Tumors were harvested and cells were cultured to determine IRS2 protein levels. Western blot revealed elevated expression of IRS2 protein in the brain tumor-derived (Br) cells compared to the parental (P) cells (Fig. 1H). Taken together, these data indicate upregulation of IRS2 under brain environment. Importantly, upregulation of IRS2 under these conditions is independent of IRS2 gene amplification.

### IRS2 amplification enhances tumorigenicity of CRC cells within the brain environment

IRS2 protein is an adaptor in the insulin and IGF signaling cascades [30], both play pivotal role in the regulation of tumor progression and metabolism. In order to test the effects of IRS2 on CRC cells, we either overexpressed IRS2 in the IRS2 non-amplified CRC cell lines HCT116, HT29 and SW480 cells (denoted HCT116^IRS2^, HT29^IRS2^, and SW480^IRS2^) (Fig. 2A, S3), or silenced it in the IRS2-amplified cell lines SW403 and LS513 cells (denoted SW403^sh-IRS2^ and LS513^sh-IRS2^) (Fig. 2A) and assessed the effect of these manipulations on oncogenic characteristics of the cells. HCT116^IRS2^, HT29^IRS2^, and SW480^IRS2^ cells showed significantly increased proliferation, colony formation, migration, invasion, and 3D sphere formation compared to cells expressing control plasmid (denoted HCT116^CON^, HT29^CON^, and SW480^CON^) (Fig. 2B-F). On the other hand, SW403^sh-IRS2^ and LS513^sh-IRS2^ cells showed decreased proliferation, colony formation, migration, invasion, and 3D sphere formation compared to cells expressing control sh-RNA (denoted SW403^sh-NS^ and LS513^sh-NS^) (Fig. 2B-F).

**Figure 2:**
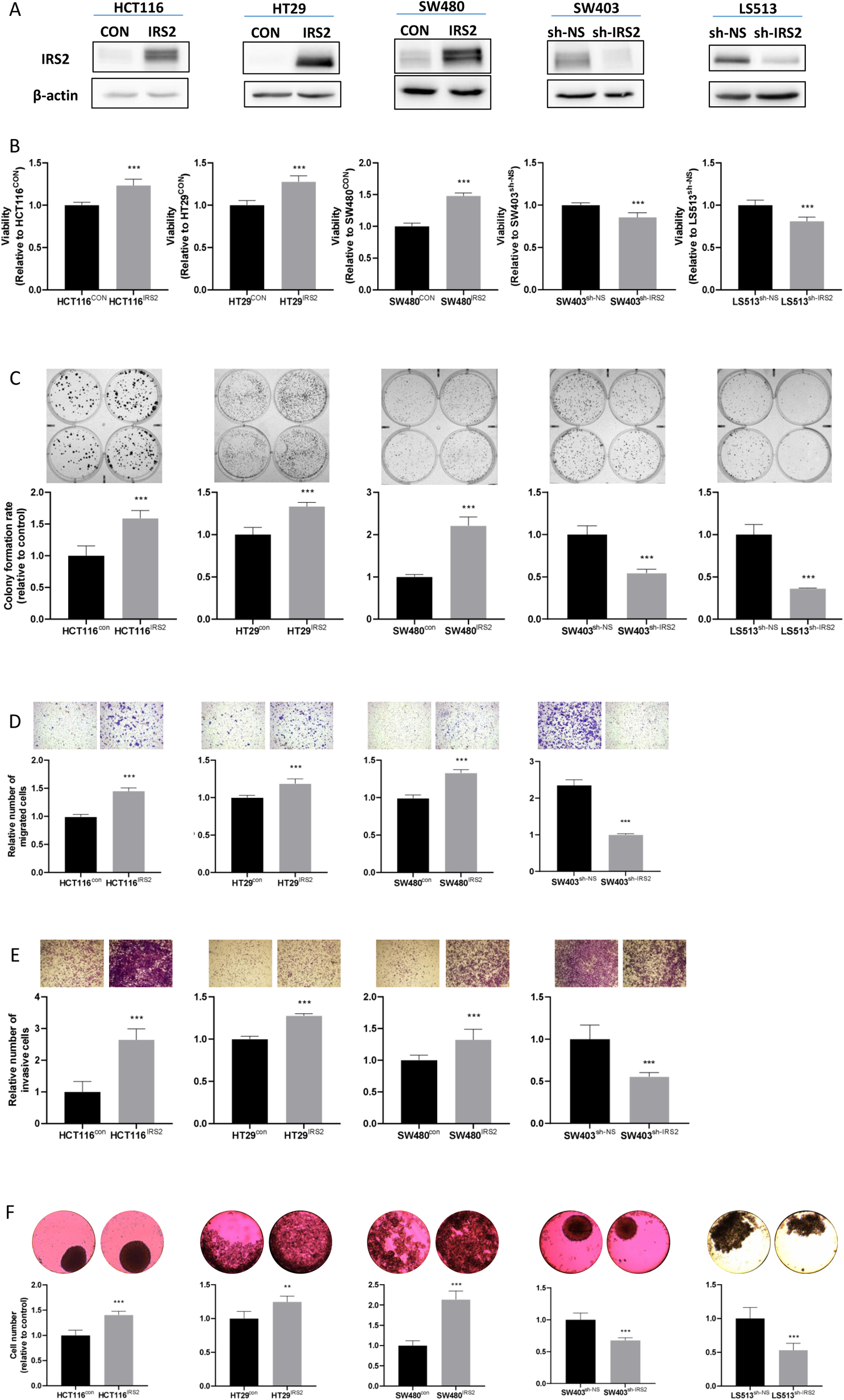
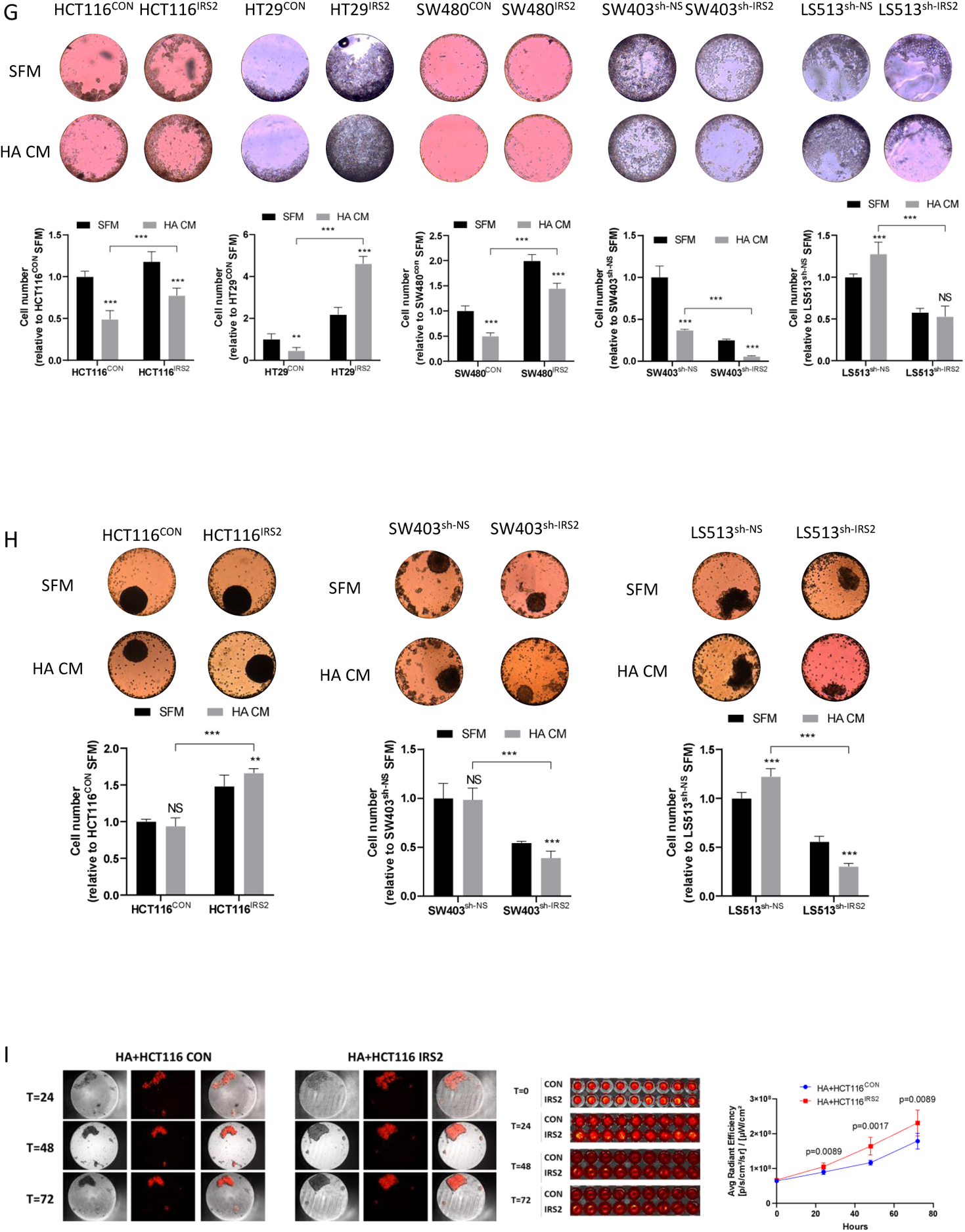
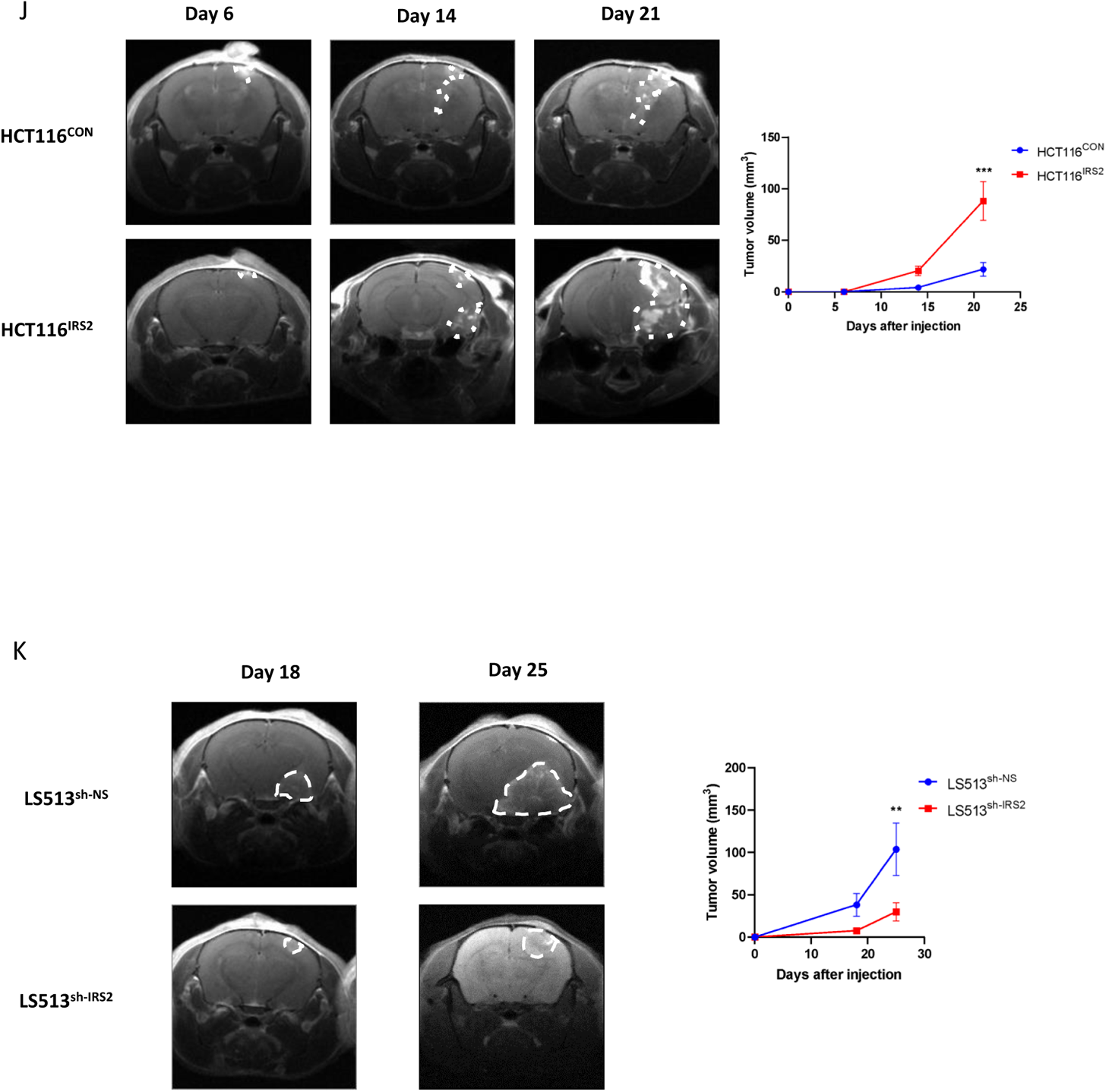
IRS2 enhances tumorigenicity of CRC cells within the brain environment. HCT116, HT29 and SW480 cells were infected with pReceiver-Lv247-IRS2 (IRS2) or pReceiver-Lv247-Empty control (CON) and SW403 and LS513 cells were infected with shRNA against IRS2 (sh-IRS2) or nonspecific sequence control (sh-NS). (A) Western blot of HCT116CON-HCT116IRS2, HT29CON-HT29IRS2, SW480CON-SW480IRS2, SW403sh-NS-SW403sh-IRS2, and LS513sh-NS-LS513sh-IRS2 cells. β-actin was used as a loading control. (B) Manipulated cells were seeded in 96-well plates and viability was assessed 72 h later using methylene blue assay. (C) Manipulated cells were seeded at low density, and 15 days later cells were fixed, and colonies were stained and quantified. (D, E) Cells, as indicated, were seeded in transwell migration assay (D) or transwell invasion assay (E) and the number of cells was quantified after 48h. (F) Spheres of manipulated cells were generated using inSphero assay, and their viability was evaluated after seven days. The figures show representative results of three independent experiments. Each bar represents ± SD, statistical analysis was performed using unpaired t test. (G) Manipulated cells were seeded in HA CM or SFM using inSphero assay. As those conditions did not support 3D spheres generation, we utilized the assay for 2D culture viability. Viability was evaluated by Realtime-Glow MT, and cells were photographed after five days. (H) Sphere of HCT116CON or HCT116IRS2, SW403sh-NS or SW403sh-IRS2, LS513sh-NS or LS513sh-IRS2 were generated, and medium was changed to HA CM or SFM. Viability was evaluated by Realtime-Glow MT and spheres were photographed after 72h. (I) Sphere of HA were generated, and after three days HCT116CON or HCT116IRS2 cells expressing m-cherry were added. Cells were photographed after 24, 48, and 72 h. Viability was evaluated at the indicated time by m-cherry quantification using IVIS. Statistical analysis was performed using Repeated measures two-way ANOVA. (J) Intracranial CRC BM mouse model was employed using HCT116CON or HCT116IRS2 (n=12, per group). MRI T1-weighted gadolinium contrast was conducted on days 6, 14, and 21 post cells implantation. Images of a representative mouse brain from each group is shown. Tumor volume was quantified using MRIcro software. Statistical analysis was performed using repeated measures ANOVA. (K) The same BM model as in (J) was employed using LS513sh-NS or LS513sh-IRS2 (n=7, per group). Tumors were images and quantified similarly, on days 18 and 25 post cells implantation. Statistical analysis was performed using repeated measures ANOVA.

In order to assess the effect of IRS2 on CRC cell tumorigenic phenotypes within the brain microenvironment, we studied the effects of IRS2 manipulations on cells co-cultured with human astrocytes (HA) or grown in HA conditioned media (HA CM). IRS2 overexpression increased, while IRS2 silencing decreased viability of the cells. Thus, HCT116^IRS2^, HT29^IRS2^ and SW480^IRS2^ survived better (by 57%, 901%, and 188% respectively, p<0.0005), while SW403^sh-IRS2^ and LS513^sh-IRS2^ showed reduced viability (by 85% and 59% respectively, p<0.0005) under HA CM condition compared to control cells (Fig. 2G). Next, the ability of manipulated cells to form 3D spheres in the presence of HA CM was assessed. HCT116^IRS2^ formed larger spheres in HA CM by 77% compared to HCT116^CON^ (Fig. 2H, p<0.0005). SW403^sh-IRS2^ and LS513^sh-IRS2^ showed a mirror image; SW403^sh-IRS2^ and LS513^sh-IRS2^ spheres in the presence of HA CM were smaller by 60% and 75% respectively, compared to control cells (Fig. 2H, p<0.0005). The ability of HCT116^IRS2^ to form 3D spheres in direct contact with HA cells was also assessed. HCT116^IRS2^ formed larger spheres when co-cultured with HA by 29% after 72h compared to HCT116^CON^ (Fig. 2I, p=0.009).

To directly address the effects of IRS2 on brain adaptation *in vivo*, manipulated CRC cells were injected into the brains of athymic nude mice and tumor growth was measured by MRI. Consistent with the *in vitro* experiments, HCT116^IRS2^ cells formed larger tumors and showed accelerated growth compared to the control cells (by 304% 21 days after injection, p<0.0005) (Fig. 2J). Moreover, LS513^sh-IRS2^ cells had decreased brain metastasis outgrowth (by 247% 25 days after injection, p<0.005) (Fig. 2K). Collectively, these data indicate growth advantage for IRS2-expressing CRC cells in the brain microenvironment.

### IRS2 amplification in CRC cells enhances expression of metastasis-associated genes

In order to gain insight into the mechanism underlying the aggressive phenotype of IRS2-amplified cells, we performed global transcriptomic analysis, using RNA-seq, on SW403^sh-NS^ and SW403^sh-IRS2^ cells grown under regular conditions. We noted differential expression of 812 genes (pFDR<0.05 and fold change>1.5) (Table S6), with more than 75% being downregulated in SW403^sh-IRS2^ cells (Fig. 3A). Pathway enrichment analysis revealed alterations in pathways associated with adhesion, migration, cell morphogenesis, and differentiation, mainly in relation to a decrease in aggressiveness-associated genes in SW403^sh-IRS2^ cells (Fig. 3B). The genes that were down/up-regulated upon IRS2 silencing included Wnt/β-catenin signaling pathway molecules (JUN, APC, SMAD3, HDAC1, CTNNB1, and RAC1), EMT markers (LOX, FGF2, and SNAI2), and MAPK signaling pathway molecules (KIT, ERBB4, and FGFR3) (Fig. 3C). Verification using qPCR analysis revealed a significant decrease in the expression of genes related to aggressiveness and metastasis formation in SW403^sh-IRS2^ cells compared to SW403^sh-NS^ cells (Fig. 3D). Protein-protein interaction analysis of the 812 DEGs with a focus on the IRS2 interacting cluster showed subnetworks of interrelated proteins (STRING [27]; Fig. 3E), with prominent nodes involving EMT markers (CDH2 and COL4A1), and MAPK signaling pathway molecules (FGF2 and KIT) (Fig. 3F). These findings identify potential dependencies in aggressiveness-related IRS2 activity in CRC cells.

**Figure 3:**
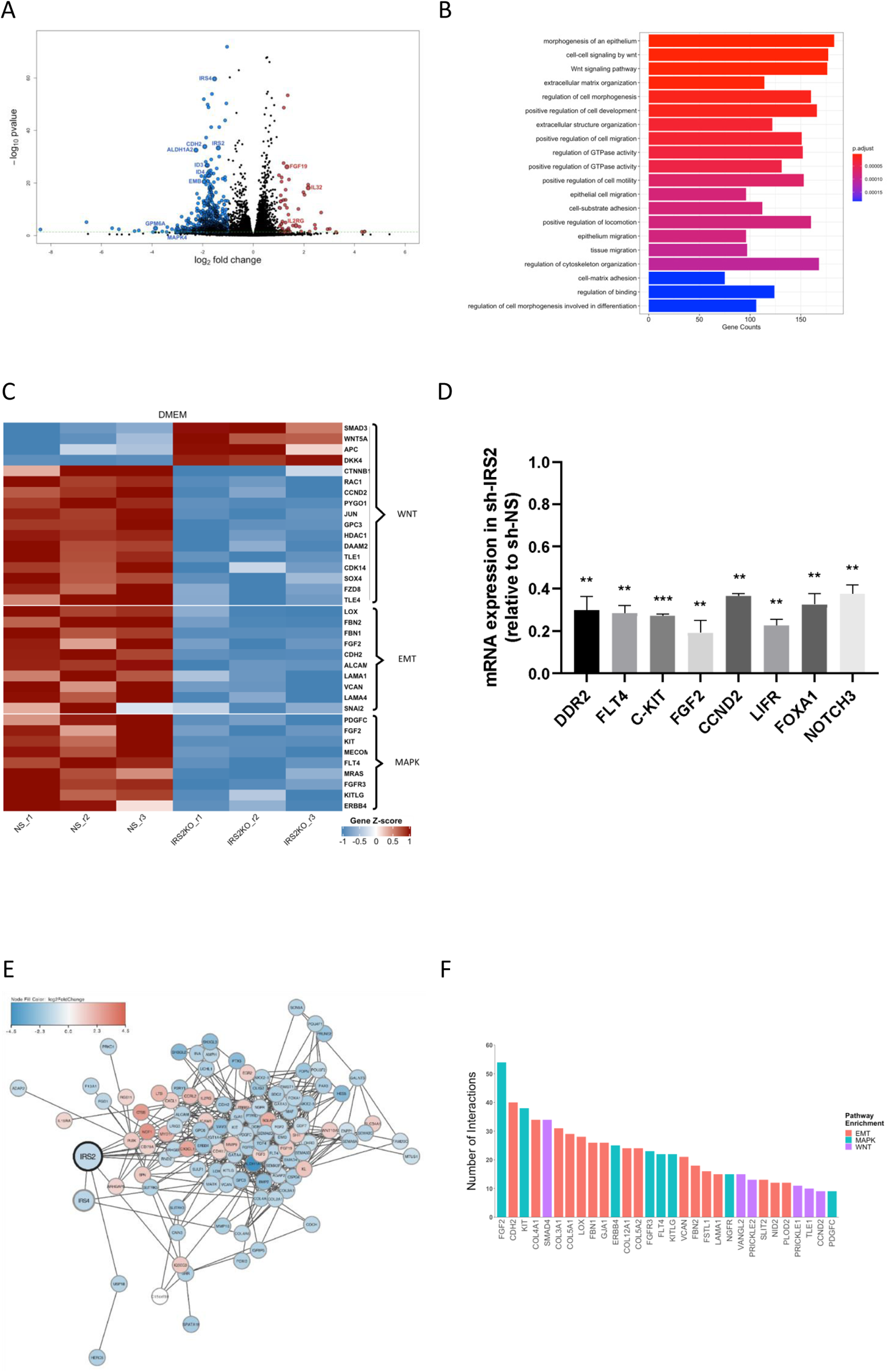
IRS2 expression enhances aggressive-associated gene signature in CRC cells. A transcriptomic analysis of SW403sh-NS or SW403sh-IRS2 cells grown under regular conditions was performed using RNAseq. (A) A volcano plot of differential gene expression between SW403sh-NS and SW403sh-IRS2 cells. Each circle represents a gene. Red color refers to significantly upregulated genes (p value < 0.05 and fold change > 2), while blue color refers to significantly downregulated genes (p value < 0.05 and fold change > 2). (B) Barplot visualization of the top 20 over-represented Gene Ontology (GO) Biological Process terms in SW403sh-NS versus SW403sh-IRS2. The color of the bars represents the P value adjusted by Benjamini–Hochberg correction for each enriched GO term identified by Fisher’s exact. The length of the bar represents the number of genes enriched in the total gene set. (C) Heatmap of significantly differentially expressed genes (p-value<0.05) grouped by their respective function in SW403sh-IRS2 compared with SW403sh-NS cells. (D) Validation of chosen downregulated genes related to metastasis formation in SW403sh-IRS2 compared to SW403sh-NS cells. (E) Protein–protein interactions among products of 897 DEGs (log (FC) > 1.5, Benjamini– Hochberg-adjusted P value < 0.05), analyzed using the search tool for retrieval of interacting genes or proteins (STRING) database. A graph was constructed using Cytoscape software and filtered to represent only IRS2 interacting cluster. Each node is colored by z-normalized expression value and the edges represent its associated protein interactions. (F) Bar-plot demonstrating the number of interactions for the top 30 most interacting proteins (only IRS2 interacting cluster). The color of the bar represents associated GO terms; red, EMT; green, MAPK; violet, WNT.

### IRS2 amplification increases mitochondrial activity and rewires metabolism of CRC cells

Metabolic adaptations permit tumor cells to metastasize to and thrive in the brain and IRS2 regulates the PI3K/AKT/mTOR metabolic pathway [31]. We, therefore, examined whether IRS2 alters CRC cells’ metabolism. We first selected 70 MSigDB KEGG metabolism-specific gene sets and performed GSEA-P on the transcriptomic analysis [32] of SW403^sh-NS^ and SW403^sh-IRS2^ cells grown under regular conditions. The analysis demonstrated that among the pathways that were significantly decreased upon IRS2 silencing was the KEGG oxidative phosphorylation (OXPHOS) gene set (p-value<0.05, FDR & q-values<0.25; Fig. 4A, B). Using the transcriptomic data, we generated a heat map of the 21 significantly differentially expressed OXPHOS associated genes (p-value<0.05) and noted down-regulation of all these genes in SW403^sh-IRS2^ cells compared to the control cells (Fig. 4C). The majority of these genes were related to respiratory complex I - NADH dehydrogenase (Fig. 5D). These global transcriptomic data were verified by qRT-PCR analysis of SW403^sh-NS^ and SW403^sh-IRS2^ cells grown under regular conditions (Fig. 4E). The same trend was observed in LS513^sh-IRS2^ cells and the opposite in HCT116^IRS2^ cells (Fig. S2). Similar patterns were also observed among SW403^sh-NS^ and SW403^sh-IRS2^ cells grown in HA CM (Fig. 4F). These data suggest an association between IRS2 expression and mitochondrial activity.

**Figure 4:**
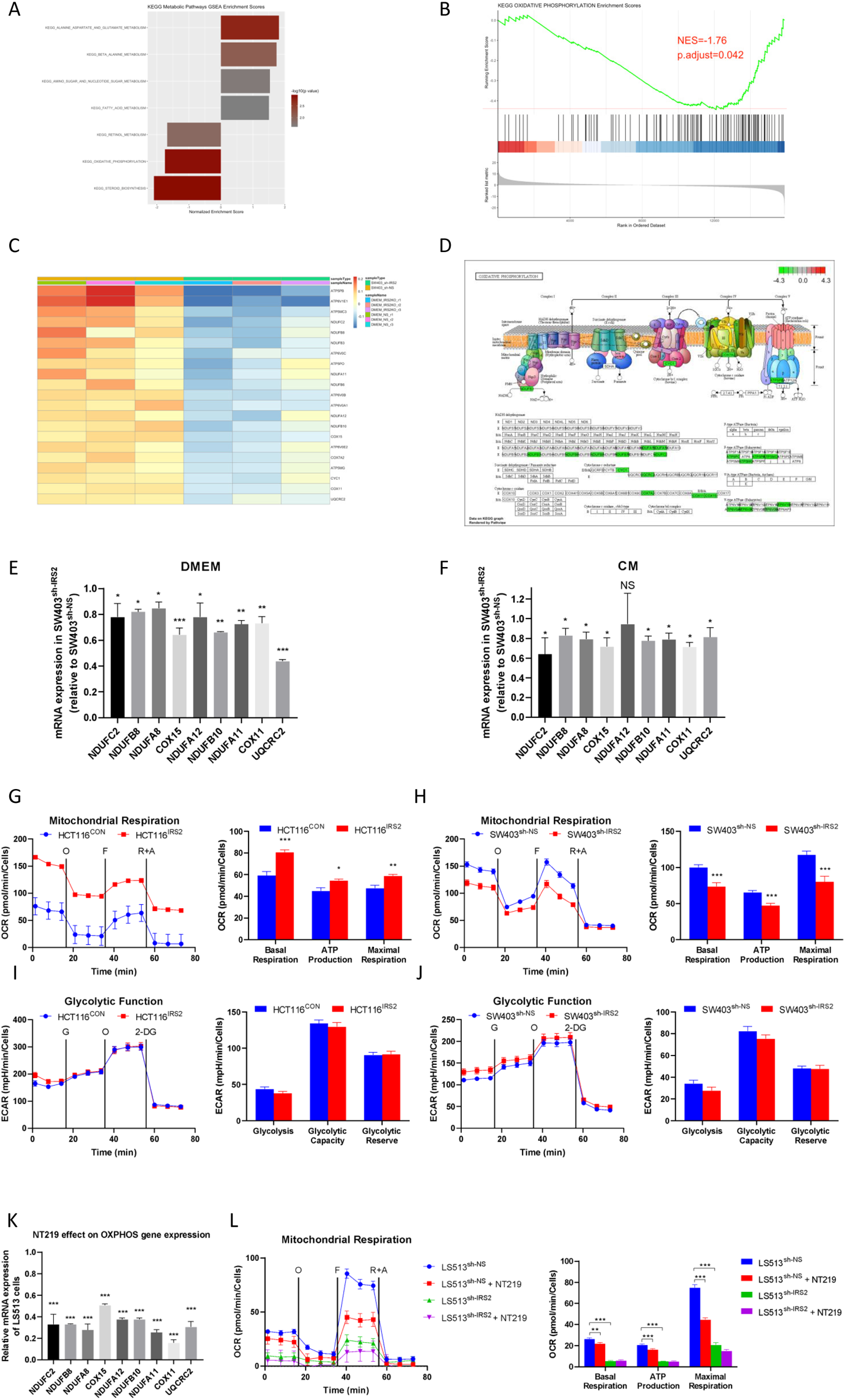
IRS2 amplification increased mitochondrial activity. (A) Pre-ranked GSEA demonstrating significantly altered KEGG metabolism pathways (P value < 0.05, FDR & q-values < 0.25) in SW403sh-IRS2 compared with SW403sh-NS cells. The normalized enrichment score (NES) forms the x-axis. (B) GSEA enrichment plot showing significant expression of the KEGG OXPHOS gene set in SW403sh-IRS2 compared with SW403sh-NS cells. NES and FDR (calculated using Benjamini– Hochberg method) are listed on the plot. (C) Heatmap of significantly differentially expressed genes (p-value<0.05) related to oxidative phosphorylation in SW403sh-IRS2 compared with SW403sh-NS cells. (D) KEGG pathway illustration of oxidative phosphorylation in human, rendered by Pathview R package. Significantly down-regulated genes are labeled by green. No up-regulated genes met the criteria for statistical significance. (E) Verification of chosen down-regulated genes related to oxidative phosphorylation in SW403sh-IRS2 compared to SW403sh-NS cells in DMEM. (F) Validation of chosen down-regulated genes related to oxidative phosphorylation in SW403sh-IRS2 compared to SW403sh-NS cells in HA CM. (G, H) Mitochondrial respiration was studied by monitoring oxygen consumption rate (OCR) using Seahorse Cell Mito Stress Test kit. (G) HCT116CON or HCT116IRS2 or (H) SW403sh-NS or SW403sh-IRS2 were seeded at a density of 10,000 cells per well. Mitochondrial respiration was measured under basal conditions followed by the sequential addition of oligomycin - “O”, FCCP - “F”, rotenone and antimycin A - “R+A”. The figure depicts a representative graph output of at least three independent experiments. Statistical analysis was performed using unpaired t test. (I, J) Glycolytic activity was studied by monitoring extracellular acidification rate (ECAR) using Seahorse Glycolysis Stress Test Kit. (I) HCT116CON or HCT116IRS2 or (J) SW403sh-IRS2 or SW403sh-NS were seeded at a density of 10,000 cells per well. Glycolytic activity was measured following sequential addition of glucose – “G”, oligomycin – “O”, and 2-DG. The figure depicts a representative graph output of at least three independent experiments. Statistical analysis was performed using unpaired t test. (K) LS513 cells were treated with NT219 (5µM) or control vehicle (D5W). Expression of genes related to oxidative phosphorylation was determined by qRT-PCR. Statistical analysis was performed using unpaired t test. (L) LS513sh-NS or LS513sh-IRS2 were seeded as in (G, H) and after a day treated with NT219 (5µM) or control vehicle (D5W) for 4 h and mitochondrial respiration was assessed as described above (G, H). The figure depicts a representative graph output of at least three independent experiments. Statistical analysis was performed using unpaired t test.

**Figure 5:**
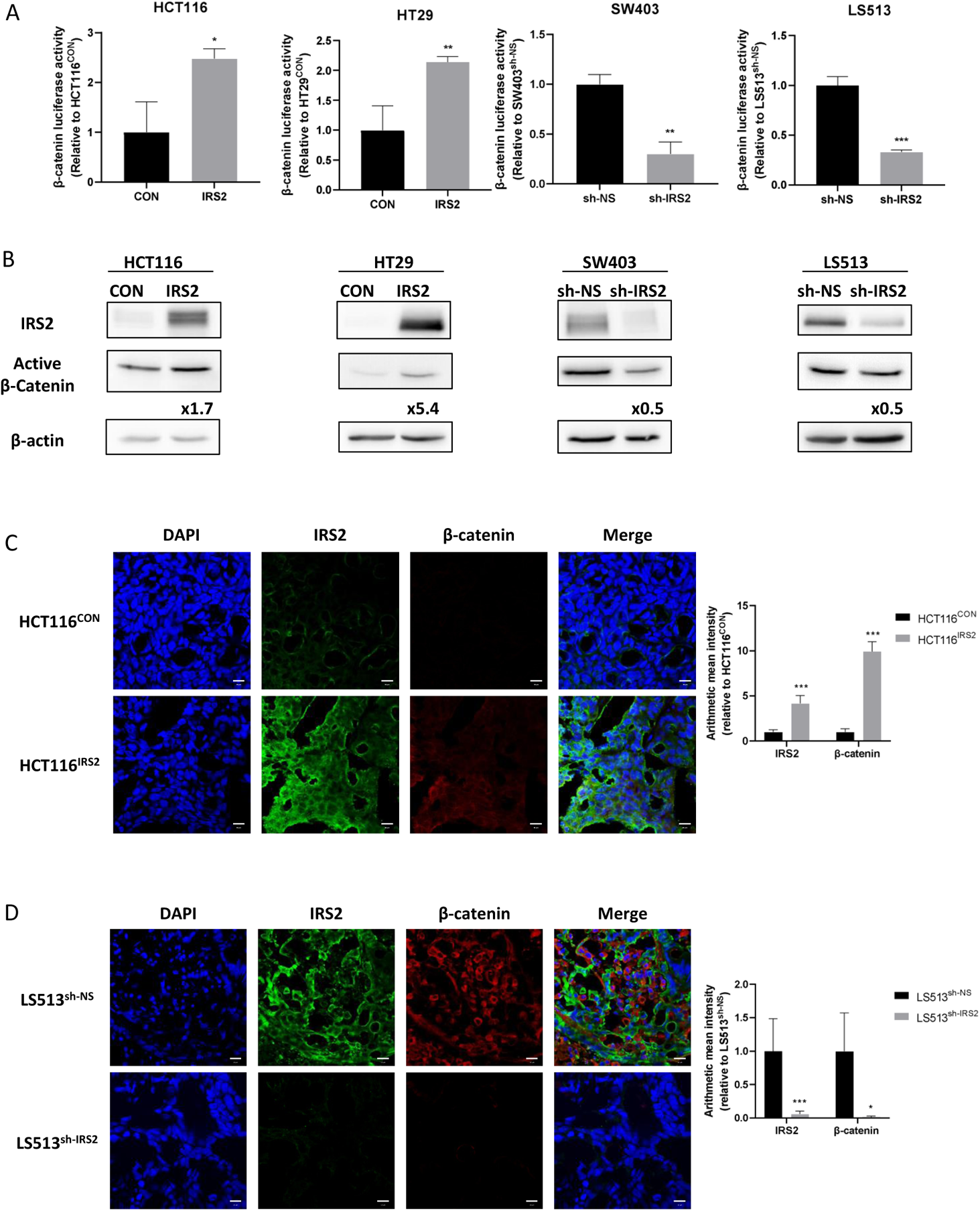
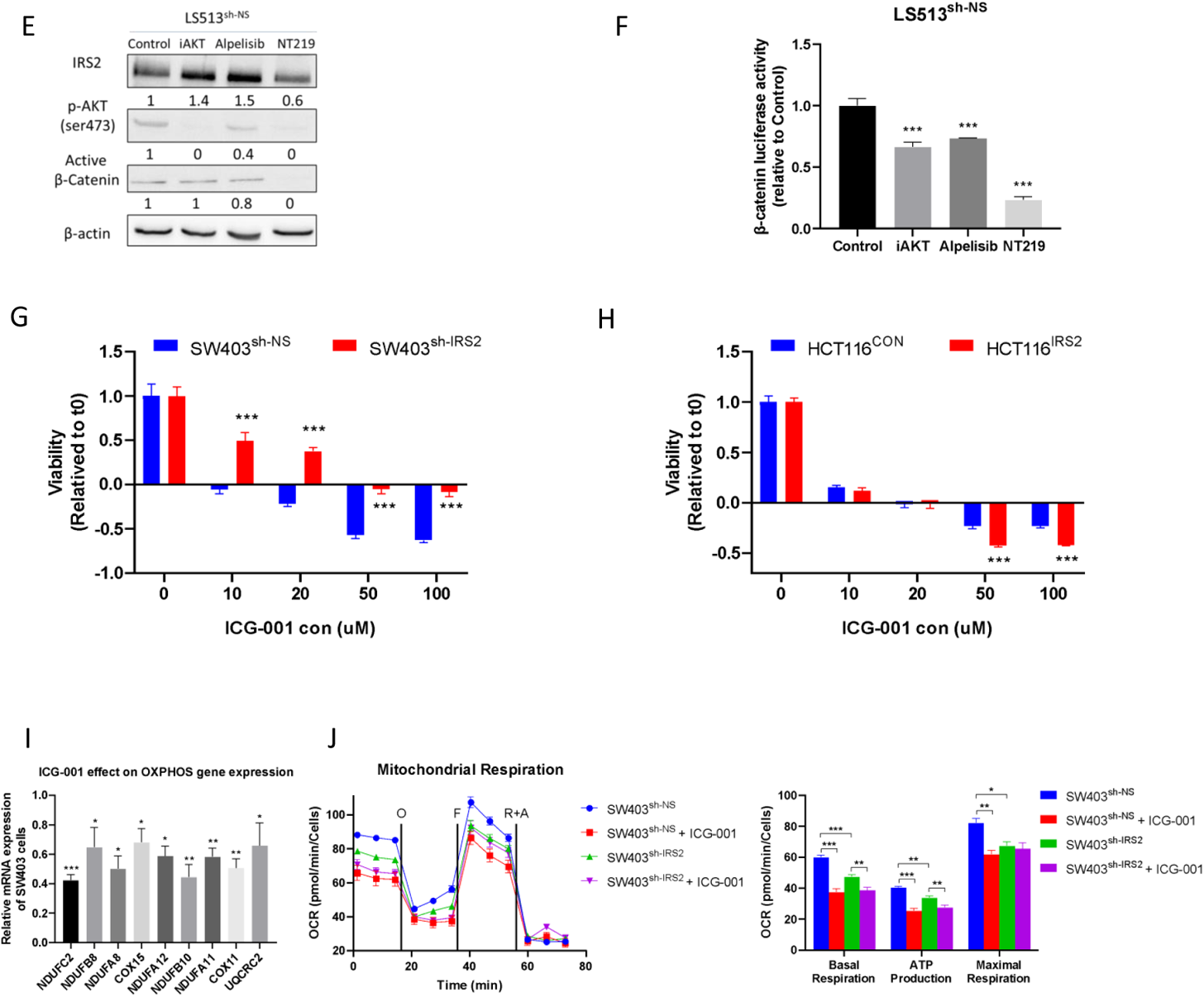
IRS2 rewired AKT and β-catenin pathway of CRC cells in the brain microenvironment. (A) Cells as depicted were transfected with either pTOPFLASH or pFOPFLASH. Luciferase activities were analyzed and normalized to total protein concentration. (B) Cells as depicted were seeded and IRS2 and active β-catenin protein levels were measured using β-actin as a loading control. (C) Mouse HCT116CON or HCT116IRS2 CRC BM tumors described in Fig. 3J were immunostained for IRS2 and β-catenin. Depicted representative images and quantification of proteins expression. Data represent mean ± s.d. Quantification was based on 3 sections from each mouse and 6 mice from each group. Statistical significance was determined using two-way ANOVA test with multiple comparisons adjustment. Scale bars represent 10 µm. (D) Experiment identical to (C), only using mouse LS513sh-NS or LS513sh-IRS2 CRC BM tumors described in Fig. 3K. (E) LS513sh-NS cells were treated with iAKT (1uM), Alpelisib (1uM), NT-219 (5uM), or control (D5W). IRS2 and active β-catenin protein levels were measured using β-actin as a loading control. (F) LS513sh-NS cells were transfected with either pTOPFLASH or pFOPFLASH. A day later cells were treated as in (E), and luciferase activities were analyzed and normalized to total protein concentration. (G) SW403sh-NS or SW403sh-IRS2 or (H) HCT116CON or HCT116IRS2 cells were seeded and 24 h later treated with elevated ICG-001 concentrations. After 72 h, viability was assessed and normalized to time 0 using methylene blue assay. Statistical analysis was performed using two-way ANOVA. (H) SW403 cells were treated with ICG-001 (20uM) or control vehicle (DMSO). Expression of genes related to oxidative phosphorylation was determined by qRT-PCR. Statistical analysis was performed using unpaired t test. (I) SW403sh-NS or SW403sh-IRS2 were seeded at a density of 10,000 cells per well and after a day treated with ICG-001 (20uM) or control vehicle (DMSO) for 4 h. Mitochondrial respiration was measured using Cell Mito Stress Test kit as described in Fig. 5G, H. The figure depicts a representative graph output of at least three independent experiments. Statistical analysis was performed using unpaired t test.

In order to establish the link between IRS2 expression and mitochondrial activity, we first measured mitochondrial respiration and glycolysis by Seahorse extracellular flux analysis in HCT116^IRS2^ cells or SW403^sh-IRS2^ cells compared to control cells. HCT116^IRS2^ exhibited increased basal mitochondrial respiration (by 36%, p<0.0005), ATP production (by 21%, p<0.05), and maximal respiration (by 24%, p<0.005) compared to control cells (Fig. 4G). The opposite was observed in SW403^sh-IRS2^, where a reduction in basal mitochondrial respiration (by 26%, p<0.0005), ATP production (by 28%, p<0.0005), and maximal respiration (by 32%, p<0.0005) was noted (Fig. 4H). Glycolysis, glycolytic capacity, and glycolytic reserve were, however, unaffected by IRS2 expression in both cell types (Fig. 4I, J). Moreover, viability of IRS2-expressing cells was less affected by 2-DG, a glucose analog, that inhibits glycolysis (Fig. S2). Collectively, these data suggest that IRS2 shifts the metabolic activity of CRC cells to rely more on OXPHOS rather than glycolysis.

In order to further establish the role of IRS2 in shifting metabolic activity toward OXPHOS, we used the first-in-class IRS2 inhibitor NT219, which triggers serine phosphorylation and subsequent degradation of IRS2 [33][34]. We observed that treatment with NT219 inhibited LS513 cells’ IRS2 expression and viability in a dose-dependent manner (Fig. S2). Importantly, NT219 reduced expression of OXPHOS genes (Fig. 4K), reduced basal mitochondrial respiration (by 17%, p<0.005), ATP production (by 21%, p<0.0005), and maximal respiration (by 40%, p<0.0005) in LS513^sh-NS^, but not in the LS513^sh-IRS2^, which do not express IRS2 (Fig. 4L). Notably, when cancer cells are deprived of glucose they switch from glycolysis to mitochondrial respiration [35], and cells that adjust better to the brain interstitium’s lower glucose levels [36] will survive and grow faster. These results suggest that IRS2 permits CRC cells adaptation to the brain environment by increasing mitochondrial respiration.

### IRS2 activates AKT and β-catenin pathways in CRC cells in the brain microenvironment

In order to investigate the mechanism of action of IRS2 in the brain environment, we first focused on the immediate IRS2 downstream PI3K/AKT/mTOR pathway [10]. We examined the effects of HA CM on manipulated HCT116 and SW403 cells, compared to IGF-1 (positive control) or serum-free media (SFM, negative control) and noted that HA CM enhanced AKT phosphorylation in HCT116^IRS2^ cells compared to control cells (Fig. S3) and the opposite in SW403^sh-IRS2^ cells (Fig. S3). These results suggested that the increased tumorigenic phenotype of IRS2-overexpressing CRC cells in the brain microenvironment relies, at least partly, on PI3K/AKT activation. Hence, we hypothesized that inhibition of these proteins will be more detrimental to cells overexpressing IRS2. Therefore, we treated cells grown in either HA CM or control media with an AKT1/2 inhibitor (iAKT) (Fig. S3) or PI3K inhibitor (Alpelisib) (Fig. S3). Both treatments significantly reduced proliferation of HCT116^IRS2^ cells compared to HCT116^CON^, but only in the presence of HA CM (Fig. S3), where the AKT pathway is activated (Fig. S3). In accordance with these results, the inhibitory effect of iAKT or Alpelisib on proliferation was more pronounced in SW403^sh-NS^ compared to SW403^sh-IRS2^, again in the presence of HA CM (Fig. S3). As the effect of AKT inhibition was relatively subtle, we hypothesized that additional mechanisms might be involved in mediating IRS2 activity.

The transcriptomic study of SW403^sh-NS^ and SW403^sh-IRS2^ cells indicated enrichment of the Wnt/β-catenin pathway (Fig. 3B). In order to examine the involvement of this pathway in mediating IRS2 activity, we first examined β-catenin expression and transcriptional activity in IRS2-expressing cells compared to control cells. IRS2 expression was directly associated with β-catenin expression and transcriptional activity (Fig. 5A, B). Furthermore, we were also able to show that IRS2 expression was directly correlated with β-catenin expression in the brain environment by staining the CRC BM tumors shown in Fig. 2D, E (Fig. 5C, D).

The connection between the PI3K/AKT/mTOR and β-catenin pathways is complex [37]. In order to study the interaction between IRS2 and β-catenin, we compared inhibition of PI3K/AKT/mTOR pathway to direct inhibition of IRS2 and explored their impact on β-catenin expression and transcriptional activity. Direct inhibition of PI3K/AKT/mTOR pathway using either iAKT or Alpelisib decreased active β-catenin expression by only 0% or 20%, respectively (Fig. 5E), and β-catenin transcriptional activity by only 30% (Fig. 5F). On the other hand, NT219 diminished the active β-catenin expression as well as activity by 80% (Fig. 5E, F). These data indicate IRS2 as a potent activator of the β-catenin pathway and suggest this activity to be mostly independent of the PI3K/AKT/mTOR pathway. If indeed IRS2 pro-proliferative effect is mediated by the β-catenin pathway, inhibition of β-catenin is expected to avert the impact of IRS2. In order to test this, we examined the effects of ICG-001, a β-catenin inhibitor, on the viability of the manipulated cells. Indeed, ICG-001 reduced proliferation of SW403^sh-NS^ by 100%, whereas a reduction of only 50% was observed in SW403^sh-IRS2^ cells (Fig. 5G, p<0.0005). A similar trend was observed in the reverse model of HCT116^IRS2^ (Fig. 5H). Finally, we aimed to evaluate the role of β-catenin in regulating cellular metabolism in IRS2-amplified cells. ICG-001 reduced OXPHOS genes expression in SW403^IRS2^ (Fig. 5I) and had more impact on the basal mitochondrial respiration, ATP production, and maximal respiration of SW403^sh-NS^ as compared to SW403^sh-IRS2^ (Fig. 5J). These results suggest the involvement of IRS2 in modulating OXPHOS through β-catenin.

### Combination of 5-FU and NT219 inhibited CRC BM and extended animal survival

We aimed to employ the data accumulated thus far in order to develop a novel treatment strategy for CRC BM. 5-FU-based therapies are the mainstay for BM treatment; however, BM usually develop at a late stage of CRC, following the development of 5-FU resistance [38]. One of the 5-FU resistance mechanisms is the activation of the β-catenin pathway [39], and our data indicate a role for IRS2 in activating the β-catenin in CRC cells (Fig. 5A, B). We, therefore, hypothesized that IRS2 inhibition might sensitize CRC cells to 5-FU. In accordance with this hypothesis, treatment with NT219 and 5-FU synergistically decreased LS513 cells proliferation *in vitro* by 73% (Fig. 6A, Fig. S4, p<0.005). Moreover, while 5-FU elevated β-catenin expression under HA CM by 80%, NT219 diminished both the 5-FU-induced and the basal level of the β-catenin expression in LS513 cells (Fig. 6B). We then evaluated the therapeutic potential of combined treatment with 5-FU and NT219 *in vivo* using our CRC BM mouse model. To this end, LS513 cells were injected intracranially and a day later mice were treated with either 5-FU, NT219, their combination or a control vehicle (Fig. 6C). While neither compound had a significant effect, the combination of 5-FU and NT219 significantly and synergistically inhibited the formation of brain metastasis by 98% (Fig. 6D, Fig. S4, p<0.005) and extended median survival of the mice by 50% (Fig. 6E, p=0.0057). These results suggest that inhibition of IRS2 may sensitize CRC cells to 5-FU, and the approach to patients with BM may be significantly impacted by agents such as NT219.

**Figure 6:**
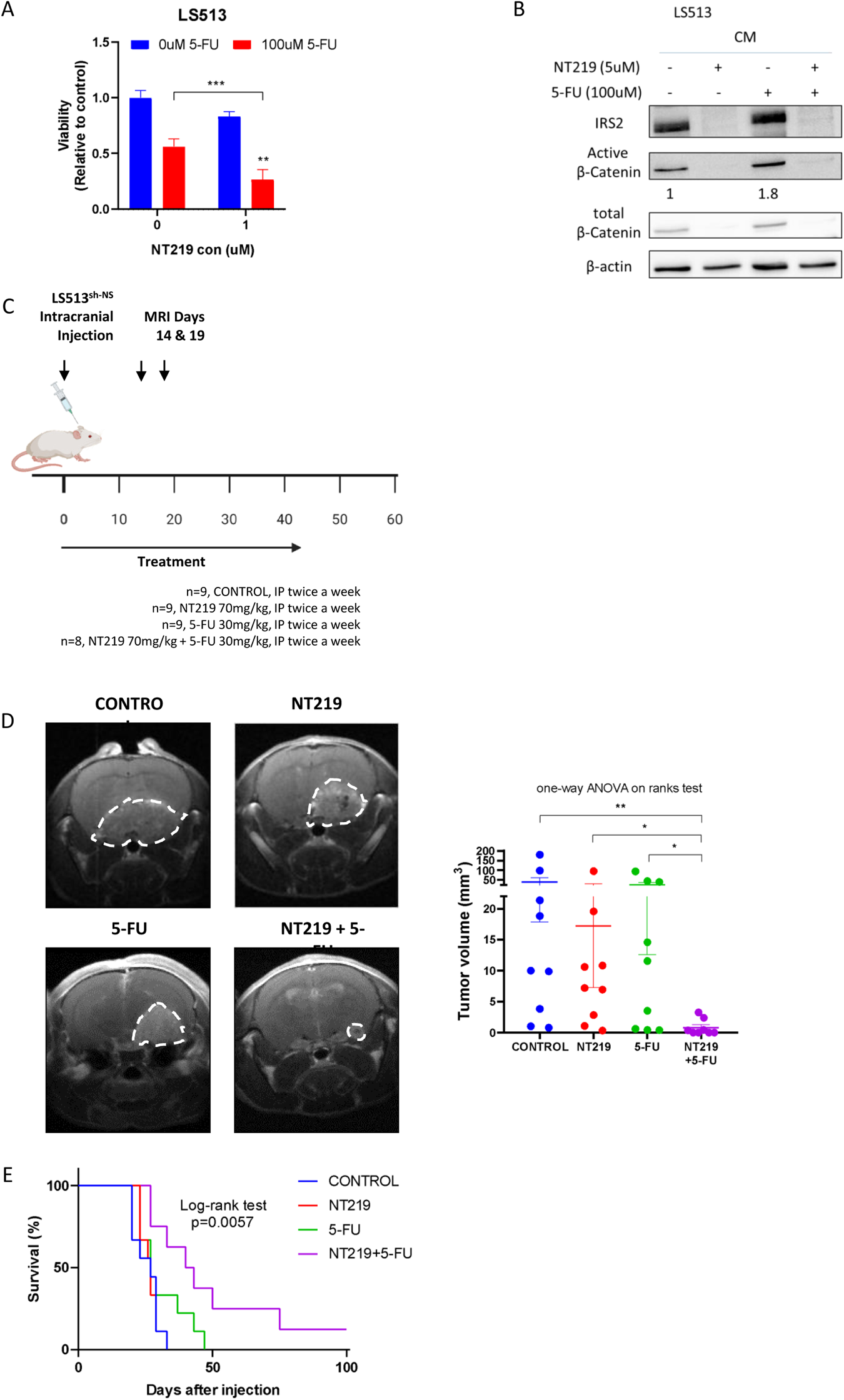
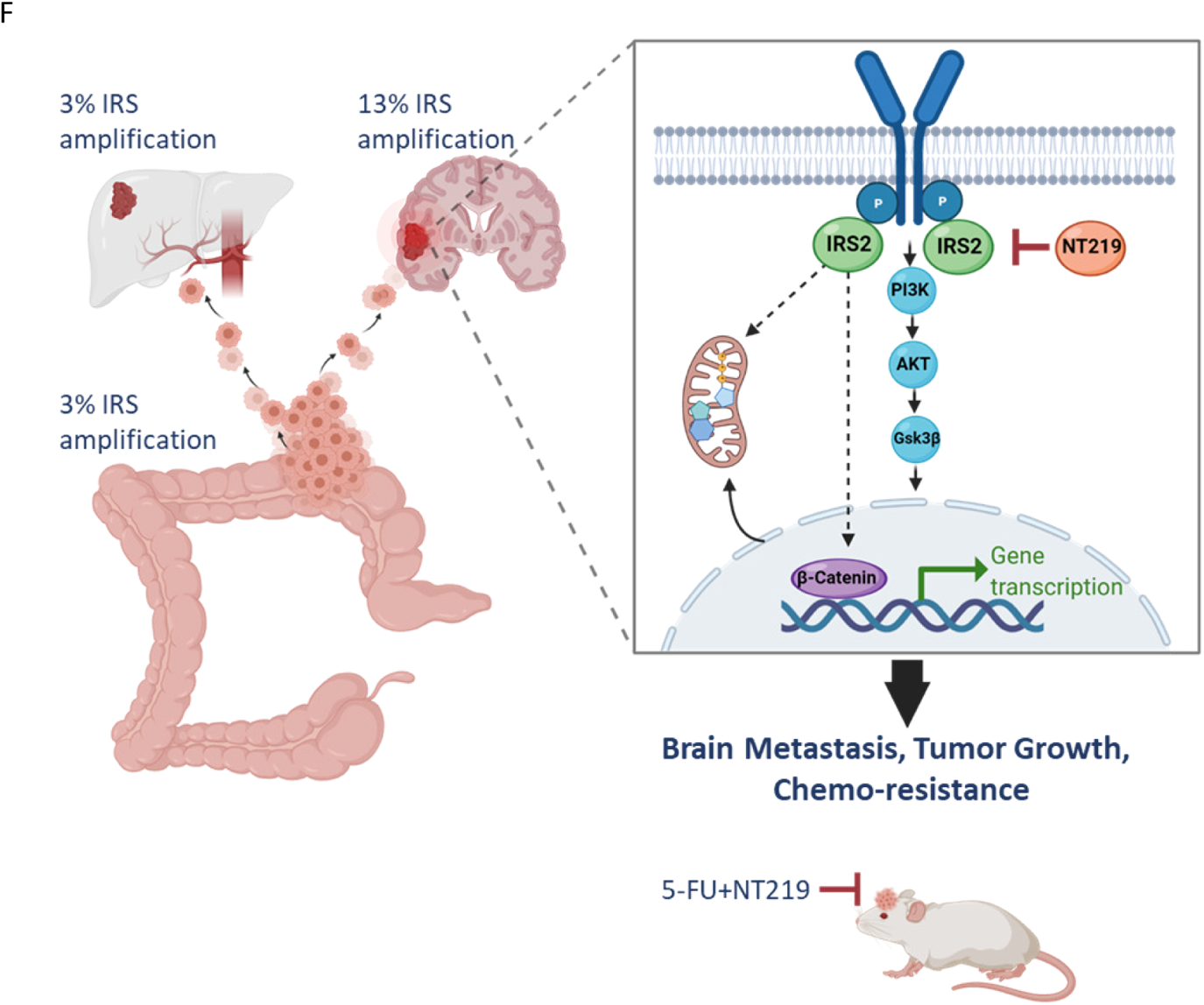
Combination of 5-FU and NT219 inhibited CRC BM and extended animal survival. (A) LS513 cells were seeded and a day later medium was replaced with HA CM and cells were treated for 24 h with either NT219 (5uM), 5-FU (100uM) or their combination. IRS2, pAKT (ser473), and active β-catenin protein levels were measured using β-actin as a loading control. (B) LS513 cells were seeded and a day later treated with either 5-FU (100uM) or NT219 (1uM) or their combination. After 72 h, viability was assessed using methylene blue assay. Statistical analysis was performed using one-way ANOVA. Bliss Synergy Score was assessed using the formula (Ea + Eb - Ea × Eb), Ea = fractional inhibition obtained by NT219, Eb = fractional inhibition obtained by 5-FU. (C) Experimental design of combined treatment of 5-FU and NT219 in vivo using intracranial LS513sh-NS cells CRC BM mouse model. (D) Mice were intracranially injected with LS513sh-NS cells and MRI T1-weighted gadolinium contrast was conducted on days 14 and 19 post cells implantation. Images of a representative mouse brain from each group on day 19 is depicted. Tumor volume was quantified using MRIcro software. Statistical analysis was performed using one-way ANOVA on ranks test. (E) Kaplan–Meyer survival curve of mice was generated. Log-rank (Mantel-Cox) test was used. 5-FU, fluorouracil. (F) A graphical illustration depicting the role IRS2 amplification plays in adaptation of CRC cells in the brain microenvironment.

## DISCUSSION

BM from CRC represent up to 8% of all diagnosed BM, and their incidence is rising due to the improvement in metastatic CRC treatment and longer survival [40]. Improving our understanding of the features of CRC BM is a critical first step to facilitate the development of new, more effective therapeutic approaches for treatment or prevention. Our global analysis of GA is the largest cohort analyzed to date of primary tumors, CRC BM, and other CRC metastatic sites, providing new insights into the pathogenesis of these tumors. Importantly, a number of these novel findings provide the rationale for testing new clinical strategies, including the targeting of IRS2, to improve the outcomes of patients with CRC BM.

An essential step in understanding the formation of CRC BM is identifying GA, which may serve as critical players in adaptation to the new environment. In recent years, attempts to discover GA in CRC BM have been made; however, they were based on a small number of clinical samples [41][42][43][44]. We noted highly statistically significant differences in the prevalence of GA between CRC BM, local tumor, and other metastatic sites (Fig. 1). As the differences ranged from 0% to 7.6%, these differences could only be detected using a very large dataset. Importantly, although small, these differences may also be of major biological implications.

GA can occur as early as within the primary tumor, giving rise to metastatic precursor cells. In contrast, some GA are enriched at later time points of the metastatic cascade, when cells colonize and adapt to a new tissue microenvironment. Moreover, GA could arise from treatment pressure, which can influence therapy resistance and metastatic trajectories. As our dataset did not include matched local and metastatic samples, we could not differentiate between those possible conditions.

The putative role of IRS2 in tumor metastasis is supported by other studies. A metastatic phenotype, conferring tumor cells with the ability to invade and survive in foreign environments, has been identified in IRS2-overexpressing mammary tumors [45]. Therefore, the upregulation of IRS2 levels and activity may contribute to tumor metastasis [45], as opposed to the effects of IRS2 gene deletion [46]. The relationship between IRS2 expression and colorectal cancer has not been explored in detail. It has been demonstrated that an increase in IRS2 expression is associated with disease progression through the stages of colorectal carcinoma formation [17]. IRS2 copy number gain was observed to be sensitive to the IGF-1R/IR inhibitor BMS-754807 and could be used as a predictive biomarker in response to this inhibitor in colorectal cancer cell lines [18]. Consistent with other studies, our study clearly indicated IRS2 role in promoting aggressiveness, as evident by colony formation, migration, invasion, and 3D sphere formation (Fig. 2). Importantly, IRS2 enhanced tumorigenicity of CRC cells in *in vitro* system mimicking the brain microenvironment (Fig. 2).

To directly address whether IRS2 amplification invokes a favorable brain adaptation, an intracranial CRC BM mouse model was employed (Fig. 2). There are several limitations to consider regarding the intracranial injection. First, intracranial injections do not mirror the metastatic cascade in that they completely bypass the development of metastatic properties within the primary tumor, intravasation into the bloodstream, and penetration through the BBB [47]. Second, direct injecting into the brain induces inflammation, which may confound findings associated with neuroinflammation. Yet, this model allows testing what tumor-derived factors mediate growth within the brain, how changes in the metastatic microenvironment alter cancer growth at the site, and the effectiveness of novel therapeutic strategies on the growth of established lesions. Given that there is no spontaneous CRC BM mice model and low incidence of experimental models [38][39], the intracranial technique is the best option to assess BM development, growth, and response to therapies. Consistent with the *in vitro* experiments, IRS2-overexpressed cells generated larger brain lesions while silencing IRS2 dramatically decreased tumor outgrowth (Fig. 2). Hence, the observation that IRS2 amplification enhanced aggressive phenotype of CRC cells may provide an explanation for the increased occurrence of this GA in brain metastasis samples.

In order to reveal a mechanism through which IRS2 amplification mediates brain predilection, we conducted an unbiased transcriptomic screen, using RNAseq, of SW403^sh-NS^ (IRS2-amplified CRC cells) compared to SW403^sh-IRS2^. We noted differential expression of genes regulating migration, adhesion, and cell communication, including C-KIT, FGF2, and NOTCH3 (Fig. 3). Expression of these genes, associated with aggressive tumors, is expected to allow CRC cells to survive and penetrate the brain. However, we sought to reveal pathways that are unique to CRC, and that may specifically allow adaptation to the brain environment. We revealed significant enrichment of the oxidative phosphorylation (OXPHOS) and the Wnt/β-catenin pathway expression (Fig. 4, 5).

In order to survive and grow, extravasated cancer cells should adjust to the lower glucose levels in the brain interstitium [36]. Notably, when cancer cells are deprived of glucose, they switch from glycolysis to mitochondrial respiration [35]. Metastatic breast cancer cells have been shown to be less dependent on glucose in the brain and instead utilize mitochondrial respiration for energy production and antioxidant defense [50][36]. Moreover, molecular profiling shows enzymes involved in oxidative phosphorylation are enriched in melanoma brain metastases [32]. Treatment with IACS-010759, a novel mitochondrial complex I inhibitor, improved survival and decreased the development of brain metastases in BRAF/MEK inhibitor-resistant mice [32]. In a metastatic melanoma PDX study, β-sitosterol, an electron transport chain complex I inhibitor, effectively reduced oxidative phosphorylation and prevented the development of brain metastases [51]. Interestingly, in this study, the majority of up-regulated genes were related to respiratory complex I - NADH dehydrogenase (Fig. 4D). Indeed, IRS2-expressing CRC cells relied more on oxidative phosphorylation rather than glycolysis (Fig. 4).

It has been documented that IRS1 and Wnt/β-catenin signaling tightly linked with each other [52][53][54]. A recent study has reported that IRS1/2 regulates Wnt/β-catenin signaling via stabilizing Dvl2 to promote EMT and cell proliferation [55]. In this study, we found that IRS2 was associated with elevated active β-catenin expression and transcriptional activity in different CRC cell lines (Fig. 3, 5). Direct inhibition of IRS2 by NT219 decreased the β-catenin expression (Fig. 5).

There is an ever-increasing body of evidence demonstrating that WNT/β-catenin signaling regulates cellular metabolism in tumors. However, how this regulation is controlled is still poorly understood and likely to be highly context-dependent. For instance, in triple-negative breast cancer cells, WNT5B was shown to be able to control the expression of the OXPHOS-related genes for cytochrome c1 and the ATP synthase γ subunit through the canonical WNT pathway [56]. In contrast, in a different study on breast cancer, WNT/β-catenin signaling increased aerobic glycolysis through suppression of mitochondrial respiration by reducing the transcription of the gene for cytochrome c oxidase, which is an integral enzyme of the ETC and thus essential for OXPHOS [57]. In our study, we aimed to show that IRS-2-induced β-catenin activation enhances OXPHOS. Indeed, inhibition of β-catenin using NT219 or direct β-catenin inhibitor (ICG-001) dose-dependently inhibited IRS2-expressing CRC cells viability and OXPHOS genes expression (Fig. 4, 5), suggesting the involvement of IRS2 in modulating OXPHOS through β-catenin.

5-FU-based therapies, such as FOLFOX (5-FU, leucovorin, and oxaliplatin) or FOLFIRI (5-FU, leucovorin, and irinotecan), have been used as the standard therapy for advanced CRC [58]. Although 5-FU-based therapies have conferred survival benefits, nearly half of metastatic CRC patients are resistant to them [59]. A recent study found that WNT/β-catenin pathway activation confers 5-FU resistance of CRC cells [39]. In our study, we showed that combination of 5-FU and NT219 treatment inhibited the formation of CRC BM and extended animal survival (Fig. 6). To our knowledge, this is the first example of successful preclinical repurposing of a combination of drugs to treat colorectal-associated brain metastases.

Taken together, this study indicates the unique genomic profile of CRC BM and implies IRS2 amplification role in promoting CRC BM. These effects may be mediated, at least in part, by modulation of the β-catenin and OXPHOS pathway. Approaching CRC BM may benefit clinically from the inhibition of the IRS pathway with agents such as NT219.

