## Supplementary Figure legend for "Genomic alterations drive brain metastases formation in colorectal cancer: The role of IRS2"

**Supplementary Fig. 1: Generation of IRS2 overexpressed stably single clones of HCT116, HT29, and SW480 cells**

(A) HCT116, (B) HT29, and (C) SW480 cells were infected with pReceiver-Lv247-IRS2 or pReceiver-Lv247-Empty control vectors and selected with puromycin (0.75µg/ml). Cell lysates were prepared for western blot, and IRS2 protein level were evaluated. Depicted two clones cultured routinely separately for each cell line and mixed prior to each experiment.

**Supplementary Fig. 2: IRS2 amplification increases mitochondrial activity**

Verification of chosen down-regulated genes related to oxidative phosphorylation in (A) $SW403^{sh\text{-}NS}$ compared to $SW403^{sh\text{-}IRS2}$ cells, and (B) $HCT116^{CON}$ compared to $HCT116^{IRS2}$ in DMEM.

(C) $HCT116^{CON}$ or $HCT116^{IRS2}$ or (D) $SW403^{sh\text{-}IRS2}$ or $SW403^{sh\text{-}NS}$ were seeded and 24 h later treated with 2-DG (20mM). After 72 h, viability was assessed using methylene blue assay. Statistical analysis was performed using two-way ANOVA.

(E) LS513 cells were seeded and 24 h later treated with elevated NT219 concentrations. IRS2 protein levels were measured using β-actin as a loading control.

(F) LS513 cells were seeded and 24 h later treated with elevated NT219 concentrations. After 72 h, viability was assessed and normalized to time 0 using methylene blue assay. Statistical analysis was performed using one-way ANOVA.

**Supplementary Fig. 3: IRS2 rewired AKT pathway of CRC cells in the brain microenvironment**

(A) $HCT116^{CON}$ or $HCT116^{IRS2}$ or (B) $SW403^{sh\text{-}NS}$ or $SW403^{sh\text{-}IRS2}$ were seeded and a day later starved for 24h in serum-free medium (SFM). After overnight, cells were stimulated with SFM or IGF-1 (100ng/ul) or HA CM for 15 minutes. IRS2, pAKT (ser473), and tAKT protein levels were measured using β-actin as a loading control.

(C) $HCT116^{CON}$ or $HCT116^{IRS2}$ or (D) $SW403^{sh\text{-}NS}$ or $SW403^{sh\text{-}IRS2}$ were seeded and 24 h later treated with HA CM or SFM (control media) containing 1000nM iAKT or control (without iAKT). IRS2, pAKT (ser473), and tAKT protein levels were measured using β-actin as a loading control.

(E, F) Experiment identical to (C, D), only using Alpelisib.

(G) $HCT116^{CON}$ or $HCT116^{IRS2}$ or (H) $SW403^{sh\text{-}NS}$ or $SW403^{sh\text{-}IRS2}$ were seeded and 24 h later treated with HA CM or SFM (control media) containing 500nM or 1000nM iAKT or control (without iAKT). Viability was assessed 72h later, each group relative to itself without iAKT using methylene blue assay. Statistical analysis was performed using two-way ANOVA.

(I, J) Experiment identical to (G, H), only using Alpelisib.

**Supplementary Fig. 4: Combination of 5-FU and NT219 works in synergy *in vitro* and *in vivo***

(A, B) Bliss score was assessed for the *in vitro* (described in Fig. 7A) and *in vivo* (described in Fig. 7D) studies using the formula (Ea + Eb - Ea x Eb), Ea = fractional inhibition obtained by NT219, Eb = fractional inhibition obtained by 5-FU. Excess over Bliss (*eob*) is calculated by the observed combined effect compared with the Bliss score. A positive, negative, or null value, is used to determine a synergistic, antagonistic or no interaction, respectively.
