## Supplementary Figures for "Genomic alterations drive brain metastases formation in colorectal cancer: The role of IRS2"

**Supplementary Fig. 1:** Generation of IRS2 overexpressed stably single clones of HCT116, HT29, and SW480 cells

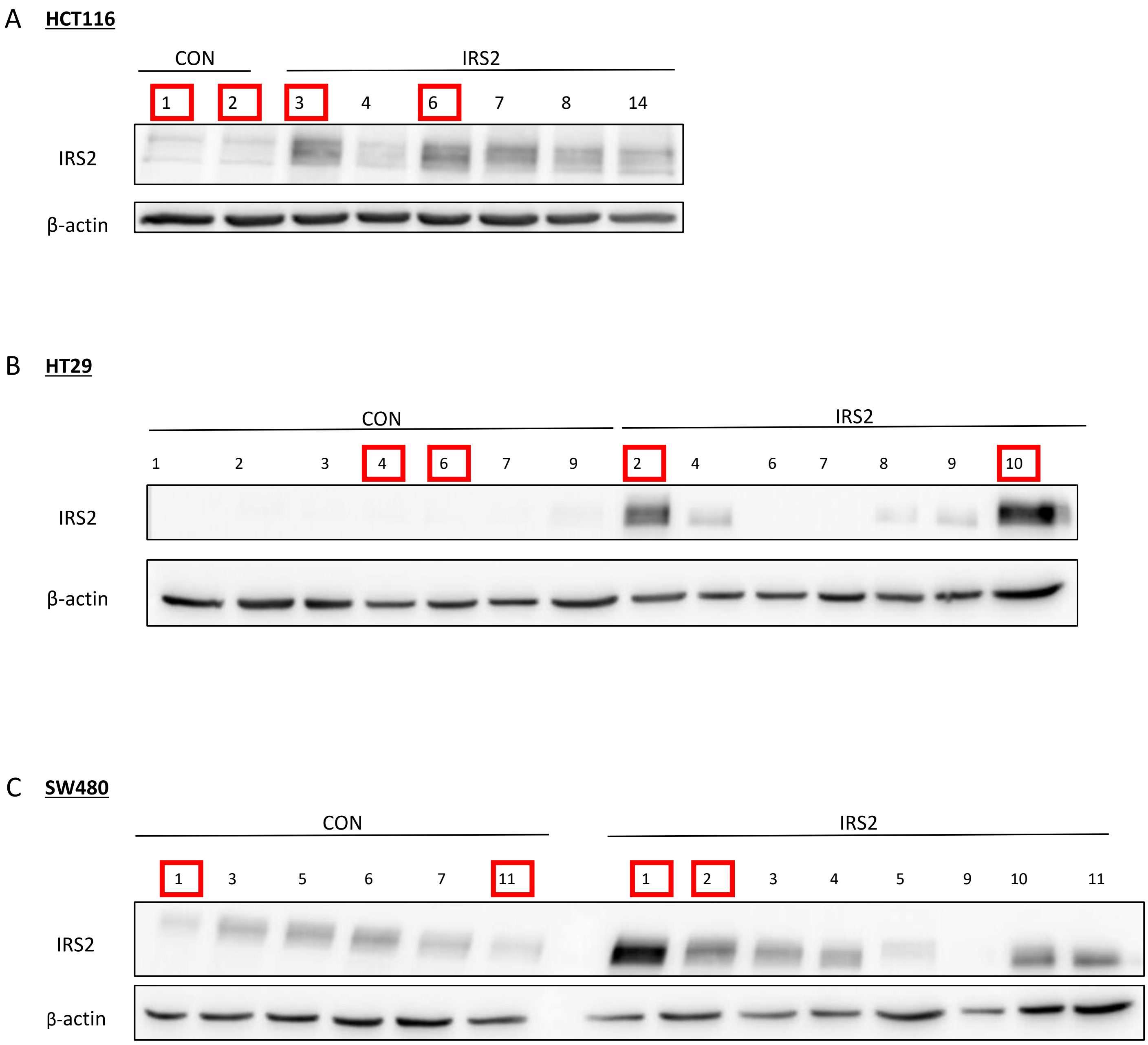

**Supplementary Fig. 2:** IRS2 amplification increases mitochondrial activity

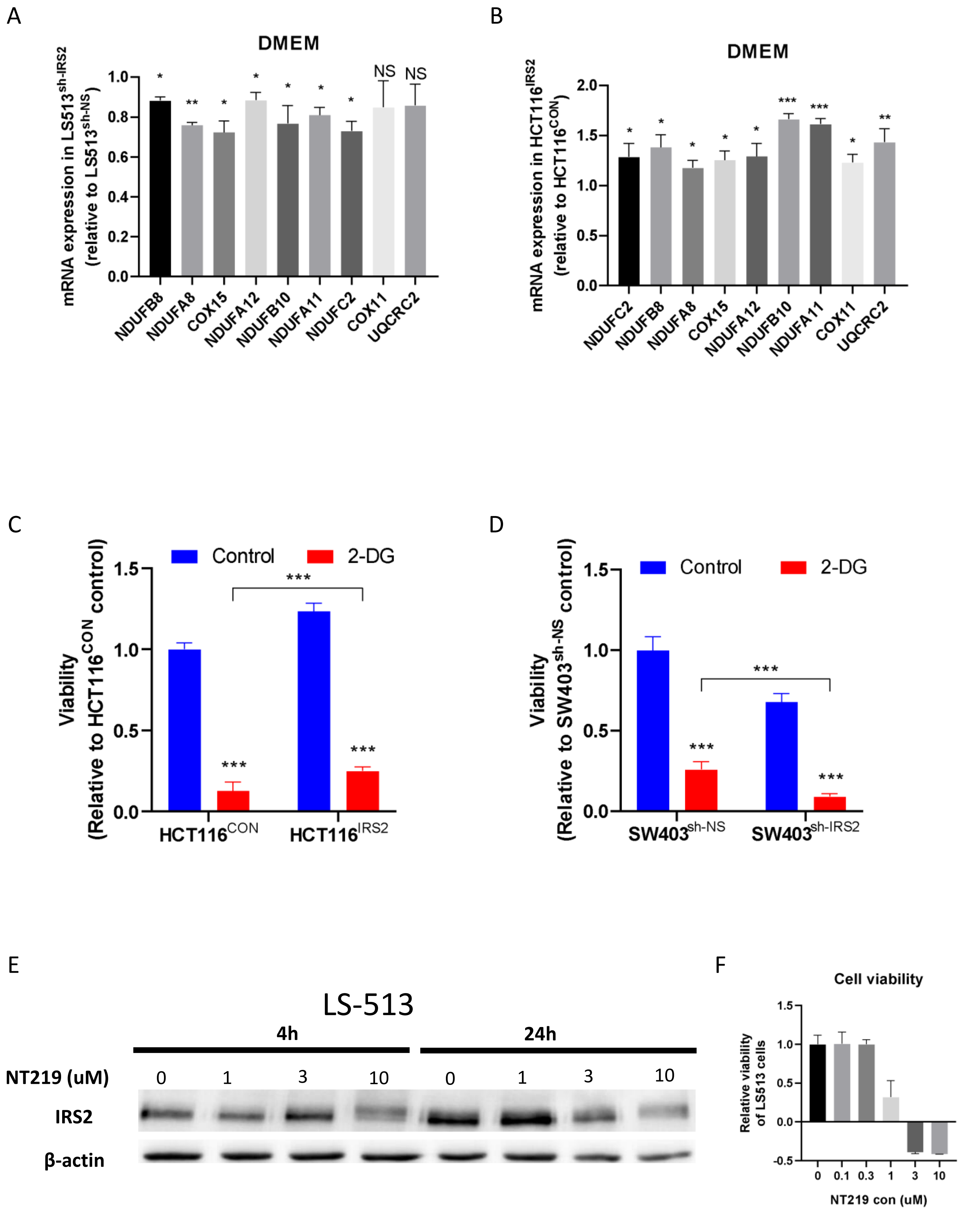

**Supplementary Fig. 3:** IRS2 rewired AKT pathway of CRC cells in the brain microenvironment

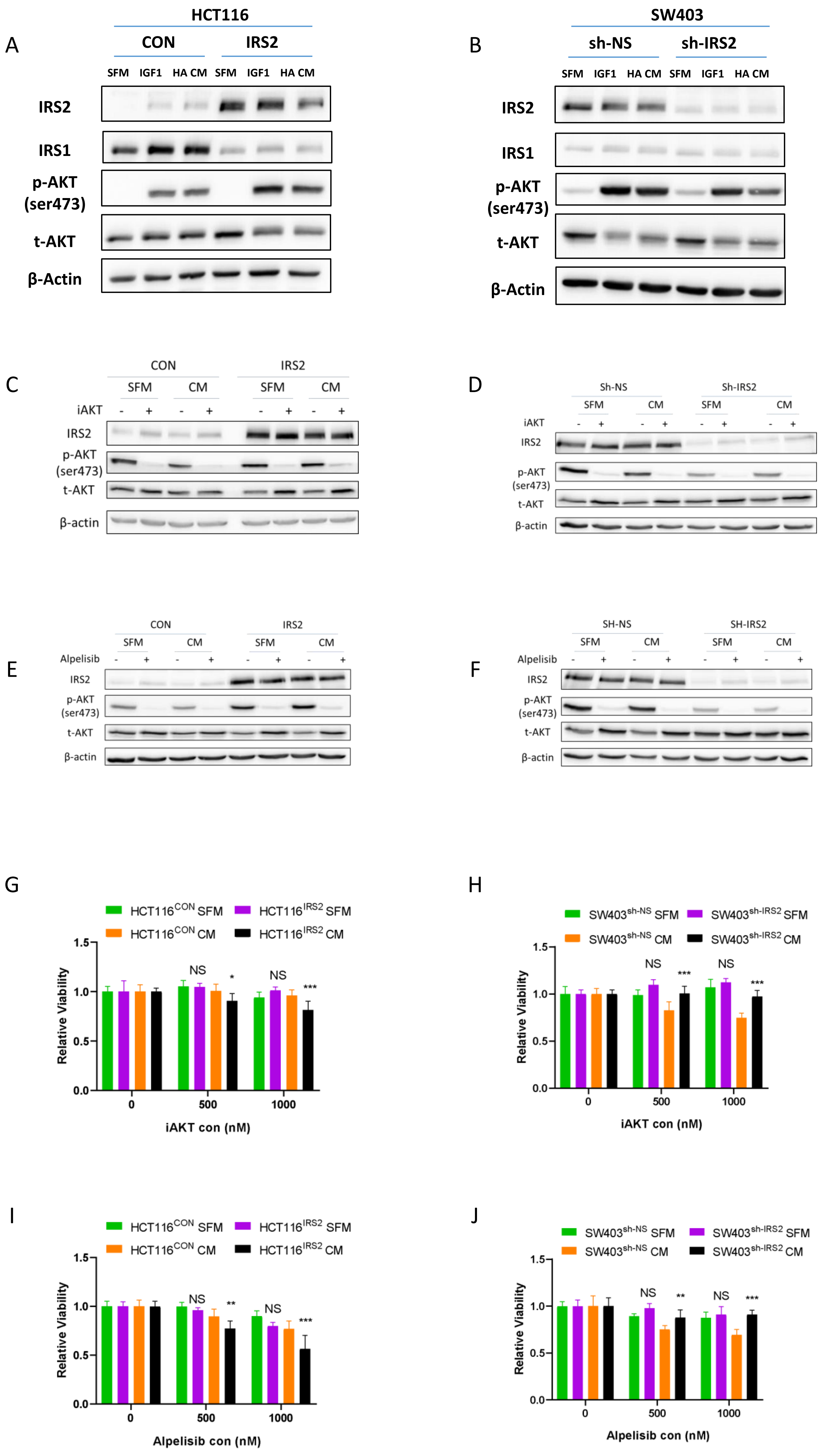

**Supplementary Fig. 4:** Combination of 5-FU and NT219 works in synergy *in vitro* and *in vivo*

A

| 5-FU vs 5-FU + NT219 (*in vitro*) | | | |
|---|---|---|---|
| Calculation | Result | | Conclusion |
| Bliss (Ea + Eb - Ea x Eb) | 0.5352 | - | - |
| Ea = fractional inhibition obtained by NT219 (1uM) | - | 0.17 | - |
| Eb = fractional inhibition obtained by 5-FU (100uM) | - | 0.44 | - |
| E a + b observed | 0.73 | >Ebliss | Synergistic |
| Excess over Bliss | 0.19 | | - |

B

| 5-FU vs 5-FU + NT219 (*in vivo*) | | | |
|---|---|---|---|
| Calculation | Result | | Conclusion |
| Bliss (Ea + Eb - Ea x Eb) | 0.730961 | - | - |
| Ea = fractional inhibition obtained by NT219 (70mg/kg) | - | 0.5516669 | - |
| Eb = fractional inhibition obtained by 5-FU (30mg/kg) | - | 0.3999128 | - |
| E a + b observed | 0.978889 | >Ebliss | Synergistic |
| Excess over Bliss | 0.25 | | - |
