## Supplementary Table 1 for "Genomic alterations drive brain metastases formation in colorectal cancer: The role of IRS2"

### Table S1: Gene lists for Version 3 (v3) of the Foundati

#### Version 3: exonic capture (n=323 genes)

| HUGO SYMBOL | HUGO SYMBOL | HUGO SYMBOL | HUGO SYMBOL | HUGO SYMBOL |
|---|---|---|---|---|
| *ABL1* | *BRCA2* | *CTNNA1* | *AS (TNFRSF* | *HIST1H2AG* |
| *ACTB* | *BRIP1* | *CTNNB1* | *FBXW7* | *HIST1H2AL* |
| *AKT1* | *BTG2* | *CUX1* | *FGF10* | *HIST1H2BC* |
| *AKT2* | *BTK* | *DAXX* | *FGF14* | *HIST1H2BK* |
| *AKT3* | *CARD11* | *DDR2* | *FGF19* | *HIST1H2BO* |
| *ALK* | *CBFB* | *DNM2* | *FGF23* | *HIST1H3B* |
| *APC* | *CBL* | *DNMT3A* | *FGF3* | *HRAS* |
| *APH1A* | *CCND1* | *DOT1L* | *FGF4* | *HSP90AA1* |
| *AR* | *CCND2* | *DTX1* | *FGF6* | *ID3* |
| *ARAF* | *CCND3* | *DUSP2* | *FGFR1* | *IDH1* |
| *ARFRP1* | *CCNE1* | *DUSP9* | *FGFR2* | *IDH2* |
| *ARHGAP26 (GRAF)* | *CD22* | *ECT2L* | *FGFR3* | *IGF1R* |
| *ARID1A* | *CD274* | *EED* | *FGFR4* | *IKBKE* |
| *ARID2* | *CD58* | *EGFR* | *FLCN* | *IKZF1* |
| *ASXL1* | *CD70* | *SY (c11orf3* | *FLT1* | *IL7R* |
| *ATM* | *CD79A* | *EP300* | *FLT3* | *INHBA* |
| *ATR* | *CD79B* | *EPHA3* | *FLT4* | *PP5D (SHIP-* |
| *ATRX* | *CDC73* | *EPHA5* | *FOXL2* | *IRF1* |
| *AURKA* | *CDH1* | *EPHA7* | *FOXO1* | *IRF4* |
| *AURKB* | *CDK12* | *EPHB1* | *FOXO3* | *IRF8* |
| *AXL* | *CDK4* | *ERBB2* | *GATA1* | *IRS2* |
| *B2M* | *CDK6* | *ERBB3* | *GATA2* | *JAK1* |
| *BAP1* | *CDK8* | *ERBB4* | *GATA3* | *JAK2* |
| *BARD1* | *CDKN1B* | *ERG* | *D4 (C17orf3* | *JAK3* |
| *BCL10* | *CDKN2A* | *ESR1* | *GNA11* | *JARID2* |
| *BCL11B* | *CDKN2B* | *ETV6* | *GNA13* | *JUN* |
| *BCL2* | *CDKN2C* | *EZH2* | *GNAQ* | *T6A (MYST* |
| *BCL2L2* | *CEBPA* | *M123B (WT* | *GNAS* | *KDM2B* |
| *BCL6* | *CHEK1* | *FAM46C* | *GPR124* | *KDM4C* |
| *BCL7A* | *CHEK2* | *FANCA* | *GRIN2A* | *KDM5A* |
| *BCOR* | *CIC* | *FANCC* | *GSK3B* | *KDM5C* |
| *BCORL1* | *CREBBP* | *FANCD2* | *HDAC4* | *KDM6A* |
| *BIRC3* | *CRKL* | *FANCE* | *HDAC7* | *KDR* |
| *BLM* | *CRLF2* | *FANCF* | *HGF* | *KEAP1* |
| *BRAF* | *CSF1R* | *FANCG* | *HIST1H1C* | *KIT* |
| *BRCA1* | *CTCF* | *FANCL* | *HIST1H1E* | *KLHL6* |

#### Version 3: Select intronic capture for rearrangement analysis (n=24 g

| HUGO SYMBOL | HUGO SYMBOL | HUGO SYMBOL | HUGO SYMBOL | HUGO SYMBOL |
|---|---|---|---|---|
| *ALK* | *BCR* | *ETV1* | *ETV6* | *JAK1* |

| | | | | |
|---|---|---|---|---|
| *BCL2* | *BRAF* | *ETV4* | *EWSR1* | *JAK2* |
| *BCL6* | *EGFR* | *ETV5* | *FGFR2* | *MLL* |

**ionOne Assay**

| HUGO SYMBOL | HUGO SYMBOL | HUGO SYMBOL | HUGO SYMBOL |
|---|---|---|---|
| *KRAS* | *NFE2L2* | *PTPN11* | *STAG2* |
| *LEF1* | *NFKBIA* | *PTPN2* | *STAT3* |
| *LRP1B* | *NKX2-1* | *TPN6 (SHP-1* | *STAT4* |
| *LRRK2* | *NOTCH1* | *PRO (PTPRC* | *STAT6* |
| *MAGED1* | *NOTCH2* | *RAD21* | *STK11* |
| *MAP2K1* | *NPM1* | *RAD50* | *SUFU* |
| *MAP2K2* | *NRAS* | *RAD51* | *SUZ12* |
| *MAP2K4* | *NT5C2* | *RAF1* | *TCF3* |
| *MAP3K1* | *NTRK1* | *RARA* | *TET2* |
| *MAP3K14* | *NTRK2* | *RB1* | *TGFBR2* |
| *MAP3K7* | *NTRK3* | *RELN* | *B4XP8 (TMSL3)* |
| *MCL1* | *NUP93* | *RET* | *TNFAIP3* |
| *MDM2* | *NUP98* | *RICTOR* | *TNFRSF11A* |
| *MDM4* | *P2RY8* | *RNF43* | *TNFRSF14* |
| *MED12* | *PAG1* | *ROS1* | *TNFRSF17* |
| *MEF2B* | *PAK3* | *RPTOR* | *TOP1* |
| *MEF2C* | *PALB2* | *RUNX1* | *TP53* |
| *MEN1* | *PAX5* | *SETBP1* | *TRAF2* |
| *MET* | *PBRM1* | *SETD2* | *TRAF3* |
| *MITF* | *PCLO* | *SF3B1* | *TRAF5* |
| *MLH1* | *PDCD1LG2* | *SGK1* | *TSC1* |
| *MLL* | *PDGFRA* | *SMAD2* | *TSC2* |
| *MLL2* | *PDGFRB* | *SMAD4* | *TSHR* |
| *MPL* | *PDK1* | *SMARCA1* | *TYK2* |
| *MRE11A* | *PHF6* | *SMARCA4* | *U2AF1* |
| *MSH2* | *PIK3CA* | *SMARCB1* | *U2AF2* |
| *MSH6* | *PIK3CG* | *SMC1A* | *VHL* |
| *MTOR* | *PIK3R1* | *SMC3* | *WISP3* |
| *MUTYH* | *PIK3R2* | *SMO* | *WT1* |
| *MYC* | *PIM1* | *SOCS1* | *XBP1* |
| *MYCL1* | *PPP2R1A* | *SOX10* | *XPO1* |
| *MYCN* | *PRDM1* | *SOX2* | *ZMYM3* |
| *MYD88* | *PRKAR1A* | *SPEN* | *ZNF217* |
| *NCSTN* | *PRKDC* | *SPOP* | *ZNF703* |
| *NF1* | *PTCH1* | *SRC* | *ZRSR2* |
| *NF2* | *PTEN* | *SRSF2* | |

**genes)**

| HUGO SYMBOL | HUGO SYMBOL | HUGO SYMBOL | HUGO SYMBOL |
|---|---|---|---|
| *MYC* | *PDGFRB* | *RARA* | *ROS1* |

| | | | |
|---|---|---|---|
| *NTRK1* | *RAF1* | *RET* | *TMPRSS2* |
| *PDGFRA* | | | |
