## Supplementary Table 2 for "Genomic alterations drive brain metastases formation in colorectal cancer: The role of IRS2"

### Table S2: Gene lists for Version 5 (v5) of the FoundationOne Ass

#### Version 5: exonic capture (n=395)

| HUGO SYMBOL | HUGO SYMBOL | HUGO SYMBOL | HUGO SYMBOL | HUGO SYMBOL | HUGO SYMBOL | HUGO SYMBOL | HUGO SYMBOL | HUGO SYMBOL |
|---|---|---|---|---|---|---|---|---|
| *ABL1* | *BRD4* | *CRLF2* | *FANCF* | *GLI1* | *KDM5A* | *MST1R* | *PHLPP2* | *RB1* |
| *ABL2* | *BRIP1* | *CSF1R* | *FANCG* | *GNA11* | *KDM5C* | *MTOR* | *PIK3C2B* | *RBM10* |
| *ACVR1B* | *BTG1* | *CTCF* | *FANCI* | *GNA13* | *KDM6A* | *MUTYH* | *PIK3C2G* | *REL* |
| *AKT1* | *BTK* | *CTNNA1* | *FANCL* | *GNAQ* | *KDR* | *MYC* | *PIK3C3* | *RET* |
| *AKT2* | *11orf30 (EMS* | *CTNNB1* | *FANCM* | *GNAS* | *KEAP1* | *YCL (MYCL1)* | *PIK3CA* | *RICTOR* |
| *AKT3* | *CARD11* | *CUL3* | *FAS* | *GPR124* | *KEL* | *MYCN* | *PIK3CB* | *RNF43* |
| *ALK* | *CASP8* | *CUL4A* | *FAT1* | *GREM1* | *KIT* | *MYD88* | *PIK3CG* | *ROS1* |
| *ALOX12B* | *CBFB* | *CUL4B* | *FAT3* | *GRIN2A* | *KLHL6* | *NBN* | *PIK3R1* | *RPA1* |
| *ER1 (FAM123* | *CBL* | *CYLD* | *FBXW7* | *GRM3* | *KMT2A (MLL)* | *NCOR1* | *PIK3R2* | *RPTOR* |
| *APC* | *CCND1* | *CYP17A1* | *FGF10* | *GSK3B* | *MT2C (MLL3)* | *NF1* | *PLCG2* | *RUNX1* |
| *APCDD1* | *CCND2* | *DAXX* | *FGF12* | *H3F3A* | *MT2D (MLL2)* | *NF2* | *PMS2* | *RUNX1T1* |
| *AR* | *CCND3* | *DDR1* | *FGF14* | *HGF* | *KRAS* | *NFE2L2* | *PNRC1* | *SDHA* |
| *ARAF* | *CCNE1* | *DDR2* | *FGF19* | *HLA-A* | *LMO1* | *NFKBIA* | *POLD1* | *SDHB* |
| *ARFRP1* | *CD274* | *DICER1* | *FGF23* | *HLA-B* | *LRP1B* | *NKX2-1* | *POLE* | *SDHC* |
| *ARID1A* | *CD79A* | *DIS3* | *FGF3* | *HLA-C* | *LRP6* | *NOTCH1* | *PPARG* | *SDHD* |
| *ARID1B* | *CD79B* | *DNMT3A* | *FGF4* | *HNF1A* | *LTK* | *NOTCH2* | *PPP2R1A* | *SETD2* |
| *ARID2* | *CDC73* | *DOT1L* | *FGF6* | *HOXB13* | *LYN* | *NOTCH3* | *PRDM1* | *SF3B1* |
| *ASXL1* | *CDH1* | *EGFR* | *FGF7* | *HRAS* | *LZTR1* | *NOTCH4* | *PREX2* | *SH2B3* |
| *ATM* | *CDH2* | *EP300* | *FGFR1* | *HSD3B1* | *MAGI2* | *NPM1* | *PRKAR1A* | *SLIT2* |
| *ATR* | *CDH20* | *EPHA3* | *FGFR2* | *HSP90AA1* | *MAP2K1* | *NRAS* | *PRKCI* | *SMAD2* |
| *ATRX* | *CDH5* | *EPHA5* | *FGFR3* | *IDH1* | *MAP2K2* | *NSD1* | *PRKDC* | *SMAD3* |
| *AURKA* | *CDK12* | *EPHA6* | *FGFR4* | *IDH2* | *MAP2K4* | *NTRK1* | *PRSS1* | *SMAD4* |
| *AURKB* | *CDK4* | *EPHA7* | *FH* | *IGF1* | *MAP3K1* | *NTRK2* | *PRSS8* | *SMARCA4* |
| *AXIN1* | *CDK6* | *EPHB1* | *FLCN* | *IGF1R* | *MAP3K13* | *NTRK3* | *PTCH1* | *SMARCB1* |
| *AXL* | *CDK8* | *EPHB4* | *FLT1* | *IGF2* | *MCL1* | *NUDT1* | *PTCH2* | *SMARCD1* |
| *BACH1* | *CDKN1A* | *EPHB6* | *FLT3* | *IGF2R* | *MDM2* | *NUP93* | *PTEN* | *SMO* |
| *BAP1* | *CDKN1B* | *ERBB2* | *FLT4* | *IKBKE* | *MDM4* | *PAK3* | *PTPN11* | *SNCAIP* |
| *BARD1* | *CDKN2A* | *ERBB3* | *FOXL2* | *IKZF1* | *MED12* | *PAK7* | *PTPRD* | *SOCS1* |
| *BCL2* | *CDKN2B* | *ERBB4* | *FOXP1* | *IL7R* | *MEF2B* | *PALB2* | *QKI* | *SOX10* |
| *BCL2A1* | *CDKN2C* | *ERCC4* | *FRS2* | *INHBA* | *MEN1* | *PARK2* | *RAC1* | *SOX2* |
| *BCL2L1* | *CEBPA* | *ERG* | *FUBP1* | *INPP4B* | *MERTK* | *PARP1* | *RAD50* | *SOX9* |
| *BCL2L2* | *CHD2* | *ERRFI1* | *GABRA6* | *INSR* | *MET* | *PARP2* | *RAD51* | *SPEN* |
| *BCL6* | *CHD4* | *ESR1* | *GALNT12* | *IRF2* | *MITF* | *PARP3* | *D51B (RAD51* | *SPOP* |
| *BCOR* | *CHEK1* | *EZH2* | *GATA1* | *IRF4* | *MKNK1* | *PARP4* | *RAD51C* | *SPTA1* |
| *BCORL1* | *CHEK2* | *FAM175A* | *GATA2* | *IRS2* | *MKNK2* | *PAX5* | *D51D (RAD51* | *SRC* |
| *BLM* | *CHUK* | *FAM46C* | *GATA3* | *JAK1* | *MLH1* | *PBRM1* | *RAD52* | *STAG2* |
| *BMPR1A* | *CIC* | *FANCA* | *GATA4* | *JAK2* | *MPL* | *PDCD1LG2* | *RAD54L* | *STAT3* |
| *BRAF* | *CRBN* | *FANCC* | *GATA6* | *JAK3* | *MRE11A* | *PDGFRA* | *RAF1* | *STAT4* |
| *BRCA1* | *CREBBP* | *FANCD2* | *GEN1* | *JUN* | *MSH2* | *PDGFRB* | *RANBP2* | *STK11* |
| *BRCA2* | *CRKL* | *FANCE* | *D4 (C17orf39* | *AT6A (MYST3* | *MSH6* | *PDK1* | *RARA* | *SUFU* |

#### Version 5: Select intronic capture for rearrangement analysis (n=31 gene

| HUGO SYMBOL | HUGO SYMBOL | HUGO SYMBOL | HUGO SYMBOL | HUGO SYMBOL | HUGO SYMBOL | HUGO SYMBOL | HUGO SYMBOL | HUGO SYMBOL |
|---|---|---|---|---|---|---|---|---|
| *ALK* | *BRCA1* | *EGFR* | *ETV5* | *FGFR1* | *KIT* | *MYB* | *NTRK1* | *RAF1* |
| *BCL2* | *BRCA2* | *ETV1* | *ETV6* | *FGFR2* | *KMT2A (MLL)* | *MYC* | *NTRK2* | *RARA* |
| *BCR* | *BRD4* | *ETV4* | *EWSR1* | *FGFR3* | *MSH2* | *NOTCH2* | *PDGFRA* | *RET* |
| *BRAF* | | | | | | | | |

**ay**

| HUGO SYMBOL |
|---|
| *SYK* |
| *TAF1* |
| *TBX3* |
| *TEK* |
| *TERC* |
| *T (promoter only)* |
| *TET2* |
| *TGFBR2* |
| *TIPARP* |
| *TNF* |
| *TNFAIP3* |
| *TNFRSF14* |
| *TNKS* |
| *TNKS2* |
| *TOP1* |
| *TOP2A* |
| *TP53* |
| *TP53BP1* |
| *TRRAP* |
| *TSC1* |
| *TSC2* |
| *TSHR* |
| *TYRO3* |
| *U2AF1* |
| *VEGFA* |
| *VHL* |
| *WISP3* |
| *WT1* |
| *XPO1* |
| *XRCC2* |
| *XRCC3* |
| *ZBTB2* |
| *ZNF217* |
| *ZNF703* |
| *ZNRF3* |

**s)**

| HUGO SYMBOL |
| --- |
| *ROS1* |
| *RSPO2* |
| *TMPRSS2* |
