## Supplementary Table 6 for "Genomic alterations drive brain metastases formation in colorectal cancer: The role of IRS2"

| ENTREZID | ENSEMBL | padj | pvalue | stat | lfcSE | log2FoldCh | baseMean | |
|---|---|---|---|---|---|---|---|---|
| 6405 | ENSG00000( | 0.007851 | 0.00126 | -3.22489 | 0.502017 | -1.61895 | 15.60459 | SEMA3F |
| 90293 | ENSG00000( | 0.000165 | 1.46E-05 | -4.33505 | 0.299038 | -1.29634 | 43.132 | KLHL13 |
| 843 | ENSG00000( | 0.001144 | 0.00013 | -3.82582 | 0.54667 | -2.09146 | 14.55942 | CASP10 |
| 6376 | ENSG00000( | 8.22E-05 | 6.62E-06 | 4.505396 | 0.539622 | 2.431212 | 16.9015 | CX3CL1 |
| 90273 | ENSG00000( | 0.065812 | 0.01659 | 2.395678 | 0.506669 | 1.213817 | 14.44991 | CEACAM21 |
| 4145 | ENSG00000( | 3.22E-06 | 1.88E-07 | -5.2107 | 0.301936 | -1.5733 | 45.76704 | MATK |
| 9254 | ENSG00000( | 0.00037 | 3.61E-05 | -4.13121 | 0.35882 | -1.48236 | 30.68257 | CACNA2D2 |
| 9235 | ENSG00000( | 6.92E-19 | 6.84E-21 | 9.376256 | 0.23171 | 2.172568 | 90.59143 | IL32 |
| 23542 | ENSG00000( | 9.96E-11 | 2.73E-12 | -6.99078 | 0.209739 | -1.46624 | 94.45365 | MAPK8IP2 |
| 25900 | ENSG00000( | 3.88E-09 | 1.33E-10 | -6.42333 | 0.231737 | -1.48852 | 72.42878 | IFFO1 |
| 80199 | ENSG00000( | 3.91E-07 | 1.91E-08 | -5.62009 | 0.237393 | -1.33417 | 64.62131 | FUZ |
| 30845 | ENSG00000( | 0.005455 | 0.000818 | -3.34661 | 0.390249 | -1.30601 | 26.66711 | EHD3 |
| 147948 | ENSG00000( | 0.024263 | 0.004803 | -2.81995 | 1.033557 | -2.91458 | 4.90388 | ZNF582 |
| 1373 | ENSG00000( | 9.37E-08 | 4.08E-09 | -5.88072 | 0.248179 | -1.45947 | 67.31709 | CPS1 |
| 57476 | ENSG00000( | 0.018027 | 0.003338 | -2.93474 | 0.460193 | -1.35055 | 19.43098 | GRAMD1B |
| 330 | ENSG00000( | 3.04E-16 | 3.81E-18 | 8.68445 | 0.233101 | 2.024358 | 85.47681 | BIRC3 |
| 9099 | ENSG00000( | 0.000291 | 2.76E-05 | -4.19258 | 0.311324 | -1.30525 | 41.38978 | USP2 |
| 2324 | ENSG00000( | 9.43E-11 | 2.58E-12 | -6.99881 | 0.229371 | -1.60532 | 72.23607 | FLT4 |
| 1462 | ENSG00000( | 6.62E-11 | 1.76E-12 | -7.05212 | 0.222327 | -1.56788 | 82.22027 | VCAN |
| 5874 | ENSG00000( | 0.047123 | 0.010929 | -2.54497 | 0.767054 | -1.95213 | 7.216125 | RAB27B |
| 2042 | ENSG00000( | 0.000104 | 8.60E-06 | -4.44975 | 0.404755 | -1.80106 | 24.08636 | EPHA3 |
| 2824 | ENSG00000( | 0.000887 | 9.76E-05 | -3.89643 | 0.266452 | -1.03821 | 51.63202 | GPM6B |
| 80243 | ENSG00000( | 0.025659 | 0.005128 | -2.79889 | 0.741962 | -2.07667 | 7.494909 | PREX2 |
| 10659 | ENSG00000( | 1.20E-16 | 1.47E-18 | -8.7916 | 0.252942 | -2.22376 | 70.83228 | CELF2 |
| 4254 | ENSG00000( | 3.97E-05 | 2.95E-06 | -4.67403 | 0.295002 | -1.37885 | 49.28815 | KITLG |
| 3604 | ENSG00000( | 0.000426 | 4.25E-05 | 4.093382 | 0.446039 | 1.825808 | 24.28835 | TNFRSF9 |
| 4052 | ENSG00000( | 2.86E-06 | 1.65E-07 | -5.23527 | 0.221929 | -1.16186 | 73.75752 | LTBP1 |
| 60529 | ENSG00000( | 0.003318 | 0.000455 | -3.5061 | 0.354494 | -1.24289 | 29.95415 | ALX4 |
| 26999 | ENSG00000( | 1.50E-15 | 2.02E-17 | -8.4929 | 0.138179 | -1.17354 | 191.7058 | CYFIP2 |
| 10758 | ENSG00000( | 0.0012 | 0.000138 | -3.81224 | 0.445638 | -1.69888 | 20.95972 | TRAF3IP2 |
| 5218 | ENSG00000( | 0.001595 | 0.000191 | -3.7303 | 0.417267 | -1.55653 | 26.80129 | CDK14 |
| 586 | ENSG00000( | 2.74E-23 | 1.81E-25 | -10.4298 | 0.171167 | -1.78524 | 139.6288 | BCAT1 |
| 11178 | ENSG00000( | 0.046904 | 0.010849 | -2.54751 | 0.913171 | -2.32631 | 5.276992 | LZTS1 |
| 2983 | ENSG00000( | 1.29E-07 | 5.81E-09 | -5.82228 | 0.314261 | -1.82971 | 39.54404 | GUCY1B1 |
| 22998 | ENSG00000( | 8.86E-07 | 4.61E-08 | -5.46583 | 0.382519 | -2.09079 | 28.40001 | LIMCH1 |
| 10402 | ENSG00000( | 0.000901 | 9.95E-05 | -3.89192 | 0.386433 | -1.50397 | 26.09426 | ST3GAL6 |
| 4804 | ENSG00000( | 8.72E-31 | 3.13E-33 | -12.0105 | 0.112162 | -1.34713 | 299.1698 | NGFR |
| 2139 | ENSG00000( | 0.007315 | 0.001157 | -3.24936 | 0.473841 | -1.53968 | 17.95962 | EYA2 |
| 2065 | ENSG00000( | 4.64E-22 | 3.53E-24 | 10.14385 | 0.138862 | 1.408597 | 208.4374 | ERBB3 |
| 2009 | ENSG00000( | 0.000206 | 1.86E-05 | -4.28143 | 0.23818 | -1.01975 | 64.94675 | EML1 |
| 2261 | ENSG00000( | 2.53E-20 | 2.23E-22 | -9.73064 | 0.175239 | -1.70519 | 133.8921 | FGFR3 |
| 54549 | ENSG00000( | 0.00024 | 2.21E-05 | -4.24217 | 0.35803 | -1.51882 | 29.80185 | SDK2 |
| 6343 | ENSG00000( | 5.29E-09 | 1.85E-10 | -6.37358 | 0.226796 | -1.4455 | 76.15182 | SCT |
| 6710 | ENSG00000( | 0.001866 | 0.000229 | -3.68511 | 0.295618 | -1.08939 | 40.31887 | SPTB |
| 6196 | ENSG00000( | 0.000126 | 1.08E-05 | -4.40078 | 0.330425 | -1.45413 | 34.86134 | RPS6KA2 |
| 6596 | ENSG00000( | 1.09E-29 | 4.11E-32 | -11.7957 | 0.140909 | -1.66212 | 212.3653 | HLTF |
| 140469 | ENSG00000( | 2.75E-11 | 6.99E-13 | 7.179645 | 0.18517 | 1.329453 | 116.0508 | MYO3B |
| 55117 | ENSG00000( | 4.79E-21 | 4.04E-23 | -9.9031 | 0.157121 | -1.55598 | 157.5021 | SLC6A15 |
| 27330 | ENSG00000( | 3.19E-08 | 1.28E-09 | -6.07018 | 0.359217 | -2.18051 | 34.1801 | RPS6KA6 |

| | | | | | | | | |
|---|---|---|---|---|---|---|---|---|
| 55679 | ENSG0000( | 0.032284 | 0.00683 | -2.70504 | 0.809147 | -2.18877 | 6.424628 | LIMS2 |
| 2121 | ENSG0000( | 0.000141 | 1.21E-05 | -4.37553 | 0.365483 | -1.59918 | 29.25542 | EVC |
| 80216 | ENSG0000( | 0.032495 | 0.006882 | -2.70248 | 0.389807 | -1.05345 | 24.47438 | ALPK1 |
| 10129 | ENSG0000( | 4.11E-05 | 3.08E-06 | -4.66511 | 0.243549 | -1.13618 | 72.68081 | FRY |
| 4854 | ENSG0000( | 5.26E-06 | 3.22E-07 | -5.11023 | 0.205678 | -1.05106 | 86.38431 | NOTCH3 |
| 59277 | ENSG0000( | 0.047147 | 0.010944 | -2.54448 | 0.571877 | -1.45513 | 11.31264 | NTN4 |
| 10512 | ENSG0000( | 0.005482 | 0.000823 | -3.34482 | 0.497526 | -1.66413 | 16.2174 | SEMA3C |
| 119 | ENSG0000( | 2.08E-15 | 2.89E-17 | -8.45078 | 0.191901 | -1.62172 | 102.6323 | ADD2 |
| 55034 | ENSG0000( | 0.000934 | 0.000104 | -3.8814 | 0.269737 | -1.04696 | 53.51301 | MOCOS |
| 8786 | ENSG0000( | 0.016436 | 0.002998 | 2.967975 | 0.354472 | 1.052063 | 30.99714 | RGS11 |
| 55796 | ENSG0000( | 2.69E-11 | 6.79E-13 | -7.18353 | 0.236545 | -1.69923 | 71.34495 | MBNL3 |
| 83992 | ENSG0000( | 0.010001 | 0.001665 | -3.14435 | 0.54487 | -1.71326 | 13.0711 | CTTNBP2 |
| 5915 | ENSG0000( | 0.001413 | 0.000167 | -3.76503 | 0.573814 | -2.16043 | 12.72434 | RARB |
| 273 | ENSG0000( | 0.019139 | 0.003591 | -2.912 | 0.496743 | -1.44651 | 14.97652 | AMPH |
| 7289 | ENSG0000( | 3.96E-54 | 4.23E-57 | 15.92523 | 0.085429 | 1.360483 | 565.1002 | TULP3 |
| 1015 | ENSG0000( | 0.601195 | 0.362034 | 0.911497 | 1.193546 | 1.087914 | 2.95893 | CDH17 |
| 26002 | ENSG0000( | 1.78E-06 | 9.73E-08 | -5.33171 | 0.397704 | -2.12044 | 27.07213 | MOXD1 |
| 285220 | ENSG0000( | 0.026538 | 0.005355 | -2.78484 | 0.536738 | -1.49473 | 13.42314 | EPHA6 |
| 8671 | ENSG0000( | 0.00746 | 0.001187 | -3.24193 | 0.524625 | -1.7008 | 17.03545 | SLC4A4 |
| 5317 | ENSG0000( | 0.426971 | 0.210926 | 1.251023 | 0.815062 | 1.019662 | 5.751964 | PKP1 |
| 93664 | ENSG0000( | 0.002329 | 0.000297 | -3.6182 | 0.439359 | -1.58969 | 19.18893 | CADPS2 |
| 1310 | ENSG0000( | 0.046575 | 0.010759 | -2.55044 | 1.553591 | -3.96234 | 2.577676 | COL19A1 |
| 23136 | ENSG0000( | 3.38E-20 | 3.02E-22 | -9.69986 | 0.19212 | -1.86354 | 107.8791 | EPB41L3 |
| 56256 | ENSG0000( | 0.000792 | 8.58E-05 | -3.92771 | 0.68377 | -2.68565 | 10.78837 | SERTAD4 |
| 9422 | ENSG0000( | 4.61E-06 | 2.78E-07 | -5.13755 | 0.337849 | -1.73572 | 38.35562 | ZNF264 |
| 3732 | ENSG0000( | 1.09E-23 | 6.76E-26 | 10.52307 | 0.096396 | 1.014387 | 437.6855 | CD82 |
| 2122 | ENSG0000( | 1.33E-07 | 6.00E-09 | -5.81686 | 0.340858 | -1.98272 | 34.71419 | MECOM |
| 231 | ENSG0000( | 1.29E-50 | 1.71E-53 | -15.3971 | 0.117102 | -1.80303 | 305.4231 | AKR1B1 |
| 22795 | ENSG0000( | 0.007669 | 0.001224 | -3.23317 | 0.409915 | -1.32533 | 25.08317 | NID2 |
| 79822 | ENSG0000( | 0.009197 | 0.001509 | -3.17301 | 0.736521 | -2.33699 | 7.942164 | ARHGAP28 |
| 23409 | ENSG0000( | 0.202633 | 0.07213 | 1.798294 | 0.625588 | 1.124992 | 9.650826 | SIRT4 |
| 29965 | ENSG0000( | 2.28E-12 | 4.91E-14 | -7.53416 | 0.205465 | -1.54801 | 88.7062 | CDIP1 |
| 3383 | ENSG0000( | 2.24E-12 | 4.83E-14 | 7.53641 | 0.165942 | 1.250604 | 155.4868 | ICAM1 |
| 8646 | ENSG0000( | 0.039956 | 0.008888 | -2.61635 | 0.492234 | -1.28785 | 14.88547 | CHRD |
| 4897 | ENSG0000( | 0.00531 | 0.000791 | -3.35598 | 0.426209 | -1.43035 | 24.34691 | NRCAM |
| 79776 | ENSG0000( | 1.92E-09 | 6.37E-11 | -6.53477 | 0.273649 | -1.78823 | 63.39186 | ZFHX4 |
| 84502 | ENSG0000( | 0.436945 | 0.218257 | 1.231176 | 0.889022 | 1.094542 | 4.810853 | JPH4 |
| 57556 | ENSG0000( | 3.65E-10 | 1.11E-11 | -6.79205 | 0.227469 | -1.54498 | 77.21674 | SEMA6A |
| 1592 | ENSG0000( | 0.008455 | 0.001375 | -3.19989 | 0.614005 | -1.96475 | 10.76531 | CYP26A1 |
| 53904 | ENSG0000( | 0.006015 | 0.00092 | -3.3139 | 0.568634 | -1.88439 | 12.77372 | MYO3A |
| 23430 | ENSG0000( | 0.047213 | 0.010965 | 2.543797 | 1.284504 | 3.267518 | 4.008084 | TPSD1 |
| 56952 | ENSG0000( | 6.14E-07 | 3.11E-08 | -5.53522 | 0.220295 | -1.21938 | 73.79584 | PRTFDC1 |
| 9424 | ENSG0000( | 1.08E-05 | 7.08E-07 | 4.959065 | 0.256633 | 1.272661 | 64.96444 | KCNK6 |
| 4113 | ENSG0000( | 0.006526 | 0.001011 | -3.28733 | 0.536783 | -1.76458 | 15.73449 | MAGEB2 |
| 27443 | ENSG0000( | 1.67E-05 | 1.13E-06 | -4.86719 | 0.261775 | -1.27411 | 52.1621 | CECR2 |
| 2953 | ENSG0000( | 0.259588 | 0.101648 | 1.636914 | 1.555634 | 2.54644 | 2.533775 | GSTT2 |
| 23331 | ENSG0000( | 8.92E-09 | 3.23E-10 | -6.28713 | 0.276736 | -1.73988 | 50.94481 | TTC28 |
| 11144 | ENSG0000( | 0.000232 | 2.12E-05 | -4.25157 | 0.353596 | -1.50334 | 30.41929 | DMC1 |
| 8542 | ENSG0000( | 0.103813 | 0.02986 | 2.171946 | 1.340216 | 2.910877 | 3.219052 | APOL1 |
| 54207 | ENSG0000( | 0.123671 | 0.037424 | 2.081112 | 0.734999 | 1.529616 | 7.39411 | KCNK10 |

| | | | | | | | | |
|---|---|---|---|---|---|---|---|---|
| 1690 | ENSG00000( | 4.87E-12 | 1.10E-13 | -7.42825 | 0.20568 | -1.52784 | 92.33876 | COCH |
| 55195 | ENSG00000( | 0.259857 | 0.101868 | 1.635864 | 0.781995 | 1.279238 | 6.546535 | CCDC198 |
| 4318 | ENSG00000( | 0.323946 | 0.140018 | 1.475722 | 1.047814 | 1.546282 | 3.72018 | MMP9 |
| 140730 | ENSG00000( | 4.34E-07 | 2.15E-08 | -5.59934 | 0.297037 | -1.66321 | 44.10832 | RIMS4 |
| 10398 | ENSG00000( | 3.80E-05 | 2.80E-06 | 4.684964 | 0.238224 | 1.116069 | 66.53223 | MYL9 |
| 1472 | ENSG00000( | 0.020349 | 0.00389 | -2.88694 | 0.942838 | -2.72192 | 5.490652 | CST4 |
| 29906 | ENSG00000( | 0.56308 | 0.324779 | 0.984685 | 1.249504 | 1.230369 | 2.594453 | ST8SIA5 |
| 284217 | ENSG00000( | 0.000379 | 3.71E-05 | -4.12462 | 0.291063 | -1.20053 | 47.14372 | LAMA1 |
| 8715 | ENSG00000( | 0.017314 | 0.003182 | -2.94955 | 0.721201 | -2.12722 | 8.393978 | NOL4 |
| 412 | ENSG00000( | 0.023835 | 0.004706 | -2.82647 | 0.595134 | -1.68213 | 11.36765 | STS |
| 5973 | ENSG00000( | 0.106539 | 0.030939 | 2.15786 | 0.634452 | 1.369059 | 9.471597 | RENBP |
| 2245 | ENSG00000( | 5.63E-08 | 2.36E-09 | -5.97088 | 0.24849 | -1.48371 | 63.30751 | FGD1 |
| 56271 | ENSG00000( | 2.14E-21 | 1.73E-23 | -9.98759 | 0.178087 | -1.77866 | 124.4023 | BEX4 |
| 54 | ENSG00000( | 0.281398 | 0.114117 | 1.579955 | 0.848367 | 1.340382 | 5.389146 | ACP5 |
| 6299 | ENSG00000( | 3.71E-05 | 2.72E-06 | -4.69061 | 0.29993 | -1.40686 | 42.25978 | SALL1 |
| 9455 | ENSG00000( | 3.31E-06 | 1.95E-07 | -5.20445 | 0.215056 | -1.11925 | 86.67309 | HOMER2 |
| 5157 | ENSG00000( | 0.04283 | 0.009697 | -2.58645 | 0.625706 | -1.61836 | 9.549989 | PDGFRL |
| 760 | ENSG00000( | 1.15E-06 | 6.12E-08 | -5.41519 | 0.219414 | -1.18817 | 79.19735 | CA2 |
| 27121 | ENSG00000( | 7.00E-147 | 4.41E-151 | 26.18072 | 0.05601 | 1.466385 | 1474.003 | DKK4 |
| 7227 | ENSG00000( | 1.38E-06 | 7.42E-08 | -5.38081 | 0.379258 | -2.04071 | 28.94873 | TRPS1 |
| 4741 | ENSG00000( | 2.82E-08 | 1.12E-09 | -6.09133 | 0.221631 | -1.35003 | 82.22984 | NEFM |
| 7991 | ENSG00000( | 2.54E-11 | 6.39E-13 | -7.19188 | 0.224761 | -1.61645 | 80.36939 | TUSC3 |
| 10382 | ENSG00000( | 1.93E-11 | 4.77E-13 | -7.2318 | 0.191973 | -1.38831 | 109.6663 | TUBB4A |
| 259307 | ENSG00000( | 0.054416 | 0.013042 | 2.482611 | 0.433971 | 1.077382 | 20.61419 | IL4I1 |
| 10148 | ENSG00000( | 0.283416 | 0.115153 | 1.575447 | 0.812801 | 1.280525 | 5.872285 | EBI3 |
| 973 | ENSG00000( | 0.514317 | 0.279047 | 1.082463 | 1.021729 | 1.105983 | 3.544556 | CD79A |
| 54854 | ENSG00000( | 0.57978 | 0.340846 | 0.952496 | 1.252581 | 1.193078 | 2.449752 | FAM83E |
| 7980 | ENSG00000( | 0.008628 | 0.001404 | -3.19374 | 0.539827 | -1.72406 | 13.8832 | TFPI2 |
| 8701 | ENSG00000( | 0.025746 | 0.005153 | -2.79729 | 0.703225 | -1.96712 | 8.496214 | DNAH11 |
| 1687 | ENSG00000( | 0.038852 | 0.008553 | -2.62943 | 0.431016 | -1.13333 | 20.81306 | GSDME |
| 1124 | ENSG00000( | 7.18E-23 | 5.02E-25 | 10.33263 | 0.108549 | 1.121596 | 319.9543 | CHN2 |
| 158471 | ENSG00000( | 6.21E-05 | 4.84E-06 | -4.57177 | 0.515365 | -2.35613 | 17.61367 | PRUNE2 |
| 23189 | ENSG00000( | 0.000974 | 0.000109 | -3.87019 | 0.30704 | -1.18831 | 41.45119 | KANK1 |
| 8777 | ENSG00000( | 1.09E-09 | 3.51E-11 | -6.6235 | 0.258769 | -1.71396 | 57.39018 | MPDZ |
| 169792 | ENSG00000( | 0.015066 | 0.002702 | -2.99976 | 0.473041 | -1.41901 | 16.75294 | GLIS3 |
| 6456 | ENSG00000( | 0.00044 | 4.41E-05 | -4.08476 | 0.687075 | -2.80654 | 10.39164 | SH3GL2 |
| 2625 | ENSG00000( | 1.10E-09 | 3.55E-11 | -6.62164 | 0.212395 | -1.40641 | 81.71961 | GATA3 |
| 219699 | ENSG00000( | 1.20E-06 | 6.39E-08 | -5.4075 | 0.213617 | -1.15513 | 84.26983 | UNC5B |
| 3195 | ENSG00000( | 0.002021 | 0.000252 | 3.660478 | 0.314655 | 1.151787 | 39.60635 | TLX1 |
| 56521 | ENSG00000( | 0.001828 | 0.000223 | -3.69087 | 0.540388 | -1.99451 | 14.43323 | DNAJC12 |
| 3384 | ENSG00000( | 0.040083 | 0.008921 | 2.615061 | 0.528496 | 1.382051 | 14.30092 | ICAM2 |
| 8153 | ENSG00000( | 6.56E-05 | 5.16E-06 | -4.55834 | 0.275392 | -1.25533 | 52.8333 | RND2 |
| 83871 | ENSG00000( | 2.53E-05 | 1.79E-06 | -4.77596 | 0.217086 | -1.03679 | 75.65217 | RAB34 |
| 1363 | ENSG00000( | 3.70E-08 | 1.50E-09 | -6.04463 | 0.222862 | -1.34712 | 82.28016 | CPE |
| 8532 | ENSG00000( | 0.036906 | 0.008024 | 2.651074 | 0.411255 | 1.090266 | 25.56274 | CPZ |
| 2743 | ENSG00000( | 0.00099 | 0.000111 | -3.86552 | 0.712807 | -2.75537 | 10.021 | GLRB |
| 23220 | ENSG00000( | 0.474519 | 0.247083 | 1.157464 | 1.029785 | 1.191939 | 3.7768 | DTX4 |
| 8825 | ENSG00000( | 3.18E-09 | 1.08E-10 | -6.45478 | 0.377809 | -2.43868 | 32.65748 | LIN7A |
| 79611 | ENSG00000( | 0.000528 | 5.46E-05 | -4.03486 | 0.471836 | -1.90379 | 19.25219 | ACSS3 |
| 1303 | ENSG00000( | 0.009472 | 0.001562 | -3.16298 | 0.748134 | -2.36633 | 8.938313 | COL12A1 |

| | | | | | | | | |
|---|---|---|---|---|---|---|---|---|
| 6492 | ENSG0000( | 0.019418 | 0.003664 | -2.90569 | 0.660221 | -1.9184 | 9.653035 | SIM1 |
| 2070 | ENSG0000( | 5.10E-08 | 2.12E-09 | -5.98809 | 0.282933 | -1.69423 | 50.48383 | EYA4 |
| 10846 | ENSG0000( | 2.05E-08 | 7.94E-10 | -6.1461 | 0.177987 | -1.09393 | 120.3749 | PDE10A |
| 90632 | ENSG0000( | 2.05E-08 | 7.94E-10 | -6.1461 | 0.177987 | -1.09393 | 120.3749 | LINC00473 |
| 3910 | ENSG0000( | 1.98E-06 | 1.10E-07 | -5.31008 | 0.397811 | -2.11241 | 27.06456 | LAMA4 |
| 9096 | ENSG0000( | 4.67E-10 | 1.44E-11 | -6.75342 | 0.329372 | -2.22439 | 41.00071 | TBX18 |
| 2690 | ENSG0000( | 0.041781 | 0.009382 | -2.59779 | 0.682728 | -1.77359 | 8.305626 | GHR |
| 4015 | ENSG0000( | 6.76E-07 | 3.44E-08 | -5.51731 | 0.349997 | -1.93104 | 34.04663 | LOX |
| 9421 | ENSG0000( | 0.001876 | 0.00023 | -3.68323 | 0.356297 | -1.31232 | 30.44437 | HAND1 |
| 7903 | ENSG0000( | 0.039656 | 0.008779 | -2.62056 | 0.626451 | -1.64165 | 10.17672 | ST8SIA4 |
| 3977 | ENSG0000( | 1.69E-09 | 5.57E-11 | -6.55486 | 0.295846 | -1.93923 | 48.64725 | LIFR |
| 590 | ENSG0000( | 0.003026 | 0.000406 | -3.53606 | 0.451981 | -1.59823 | 18.13681 | BCHE |
| 57415 | ENSG0000( | 4.52E-05 | 3.42E-06 | -4.64369 | 0.421053 | -1.95524 | 22.95021 | C3orf14 |
| 59345 | ENSG0000( | 9.44E-16 | 1.25E-17 | -8.54849 | 0.207527 | -1.77405 | 103.7524 | GNB4 |
| 23150 | ENSG0000( | 0.00084 | 9.18E-05 | -3.91136 | 0.356685 | -1.39512 | 29.61203 | FRMD4B |
| 6364 | ENSG0000( | 0.002936 | 0.000392 | 3.545644 | 0.852407 | 3.022331 | 8.629769 | CCL20 |
| 3676 | ENSG0000( | 1.79E-05 | 1.22E-06 | -4.8524 | 0.330417 | -1.60331 | 35.24283 | ITGA4 |
| 9784 | ENSG0000( | 1.30E-72 | 5.72E-76 | -18.445 | 0.056184 | -1.03632 | 1436.555 | SNX17 |
| 79745 | ENSG0000( | 0.015909 | 0.00288 | -2.98025 | 0.357862 | -1.06652 | 29.32697 | CLIP4 |
| 2591 | ENSG0000( | 0.020889 | 0.004014 | -2.87704 | 0.581579 | -1.67323 | 11.47996 | GALNT3 |
| 33 | ENSG0000( | 0.003241 | 0.000442 | -3.5138 | 0.575242 | -2.02128 | 13.58491 | ACADL |
| 2202 | ENSG0000( | 0.001728 | 0.00021 | -3.7071 | 0.697897 | -2.58717 | 10.05319 | EFEMP1 |
| 3488 | ENSG0000( | 0.000111 | 9.33E-06 | -4.43214 | 0.377808 | -1.6745 | 28.58328 | IGFBP5 |
| 80303 | ENSG0000( | 5.41E-05 | 4.16E-06 | -4.60332 | 0.331947 | -1.52806 | 33.85311 | EFHD1 |
| 7456 | ENSG0000( | 1.89E-05 | 1.29E-06 | -4.84093 | 0.35652 | -1.72589 | 31.72765 | WIPF1 |
| 5341 | ENSG0000( | 0.005687 | 0.000859 | 3.3331 | 0.406737 | 1.355695 | 25.57478 | PLEK |
| 4139 | ENSG0000( | 4.88E-07 | 2.43E-08 | -5.57796 | 0.360555 | -2.01116 | 31.41158 | MARK1 |
| 25823 | ENSG0000( | 0.117544 | 0.035066 | 2.107591 | 1.173126 | 2.47247 | 3.676051 | TPSG1 |
| 23266 | ENSG0000( | 8.00E-12 | 1.89E-13 | -7.35616 | 0.204982 | -1.50788 | 97.04591 | ADGRL2 |
| 3399 | ENSG0000( | 1.81E-27 | 7.97E-30 | -11.3437 | 0.16014 | -1.81659 | 153.449 | ID3 |
| 57821 | ENSG0000( | 0.002819 | 0.000373 | -3.55884 | 0.477164 | -1.69815 | 17.33819 | CCDC181 |
| 1266 | ENSG0000( | 2.37E-24 | 1.37E-26 | -10.6724 | 0.155535 | -1.65993 | 162.5046 | CNN3 |
| 54996 | ENSG0000( | 0.000141 | 1.22E-05 | -4.37433 | 0.51588 | -2.25663 | 17.98772 | 2-Mar |
| 6785 | ENSG0000( | 1.34E-09 | 4.36E-11 | -6.59122 | 0.28643 | -1.88793 | 48.40594 | ELOVL4 |
| 7128 | ENSG0000( | 0.014779 | 0.002642 | 3.00658 | 0.355869 | 1.069947 | 31.67845 | TNFAIP3 |
| 1490 | ENSG0000( | 6.99E-06 | 4.38E-07 | -5.05177 | 0.296869 | -1.49971 | 52.92952 | CCN2 |
| 27253 | ENSG0000( | 0.0001 | 8.32E-06 | -4.45684 | 0.4099 | -1.82686 | 25.44852 | PCDH17 |
| 894 | ENSG0000( | 4.94E-14 | 7.74E-16 | -8.05827 | 0.154374 | -1.24399 | 165.3126 | CCND2 |
| 25927 | ENSG0000( | 0.014909 | 0.00267 | -3.00333 | 0.525659 | -1.57873 | 13.9693 | CNRIP1 |
| 79817 | ENSG0000( | 0.000204 | 1.83E-05 | -4.28424 | 0.406134 | -1.73997 | 23.38102 | MOB3B |
| 63970 | ENSG0000( | 0.09146 | 0.025271 | 2.237239 | 0.951616 | 2.128993 | 5.077605 | TP53AIP1 |
| 84898 | ENSG0000( | 0.002719 | 0.000356 | -3.5709 | 0.480611 | -1.71621 | 17.66862 | PLXDC2 |
| 440073 | ENSG0000( | 0.50451 | 0.271144 | 1.100432 | 1.223467 | 1.346343 | 2.684695 | IQSEC3 |
| 4093 | ENSG0000( | 2.10E-07 | 9.79E-09 | -5.73441 | 0.236002 | -1.35334 | 72.94941 | SMAD9 |
| 83690 | ENSG0000( | 5.68E-06 | 3.49E-07 | -5.09487 | 0.37484 | -1.90976 | 30.31222 | CRISPLD1 |
| 117854 | ENSG0000( | 0.000403 | 3.99E-05 | -4.1079 | 0.428906 | -1.7619 | 22.17065 | TRIM6 |
| 57616 | ENSG0000( | 0.00746 | 0.001187 | -3.24207 | 0.500308 | -1.62203 | 15.33038 | TSHZ3 |
| 9034 | ENSG0000( | 0.366282 | 0.168137 | 1.378216 | 1.300303 | 1.792097 | 2.545362 | CCRL2 |
| 22865 | ENSG0000( | 0.000308 | 2.93E-05 | -4.17876 | 0.50489 | -2.10981 | 16.75377 | SLITRK3 |
| 3208 | ENSG0000( | 0.170581 | 0.057511 | 1.899404 | 0.712138 | 1.352638 | 8.059847 | HPCA |

| | | | | | | | | |
|---|---|---|---|---|---|---|---|---|
| 7291 | ENSG0000( | 0.038852 | 0.008554 | -2.62937 | 0.523634 | -1.37683 | 14.72956 | TWIST1 |
| 23349 | ENSG0000( | 0.266629 | 0.105496 | 1.618773 | 1.241584 | 2.009843 | 2.878467 | PHF24 |
| 170685 | ENSG0000( | 0.001018 | 0.000114 | -3.85774 | 0.415915 | -1.60449 | 24.00901 | NUDT10 |
| 1959 | ENSG0000( | 0.50756 | 0.273657 | 1.09468 | 1.08619 | 1.18903 | 3.109995 | EGR2 |
| 64168 | ENSG0000( | 0.007756 | 0.001241 | -3.22927 | 0.572391 | -1.84841 | 12.18689 | NECAB1 |
| 64333 | ENSG0000( | 0.359843 | 0.163509 | 1.393367 | 0.787787 | 1.097677 | 6.556109 | ARHGAP9 |
| 1757 | ENSG0000( | 5.58E-05 | 4.31E-06 | 4.596012 | 0.238058 | 1.094118 | 71.09277 | SARDH |
| 3759 | ENSG0000( | 0.043626 | 0.009915 | -2.57876 | 1.326982 | -3.42197 | 3.547834 | KCNJ2 |
| 2162 | ENSG0000( | 0.02238 | 0.004364 | -2.85056 | 0.544815 | -1.55303 | 13.46657 | F13A1 |
| 4337 | ENSG0000( | 4.37E-05 | 3.30E-06 | -4.65148 | 0.327095 | -1.52147 | 34.73854 | MOCS1 |
| 11166 | ENSG0000( | 0.003351 | 0.00046 | -3.50297 | 0.63092 | -2.21009 | 10.62061 | SOX21 |
| 644524 | ENSG0000( | 0.014473 | 0.002576 | -3.01422 | 0.454716 | -1.37062 | 19.78138 | NKX2-4 |
| 650 | ENSG0000( | 2.25E-06 | 1.26E-07 | -5.28418 | 0.500241 | -2.64336 | 19.6604 | BMP2 |
| 154796 | ENSG0000( | 5.16E-42 | 1.05E-44 | -14.0281 | 0.095954 | -1.34605 | 578.9724 | AMOT |
| 55906 | ENSG0000( | 5.07E-07 | 2.54E-08 | -5.57066 | 0.406843 | -2.26638 | 28.30924 | ZC4H2 |
| 57110 | ENSG0000( | 0.045093 | 0.010323 | -2.56483 | 1.06859 | -2.74075 | 4.529047 | PLAAT1 |
| 83460 | ENSG0000( | 1.39E-44 | 2.53E-47 | -14.4494 | 0.076968 | -1.11214 | 688.6332 | EMC6 |
| 8910 | ENSG0000( | 0.001183 | 0.000135 | -3.81634 | 0.390372 | -1.48979 | 24.45984 | SGCE |
| 9514 | ENSG0000( | 0.033152 | 0.007055 | 2.694232 | 0.539679 | 1.454022 | 13.46616 | GAL3ST1 |
| 80833 | ENSG0000( | 0.277025 | 0.111511 | 1.591438 | 0.638574 | 1.016251 | 9.671389 | APOL3 |
| 80115 | ENSG0000( | 3.05E-21 | 2.51E-23 | 9.950419 | 0.112235 | 1.11679 | 322.0358 | BAIAP2L2 |
| 51200 | ENSG0000( | 0.025823 | 0.005175 | -2.79593 | 0.886478 | -2.47853 | 5.852187 | CPA4 |
| 93986 | ENSG0000( | 1.93E-05 | 1.33E-06 | -4.83571 | 0.510865 | -2.47039 | 22.42361 | FOXP2 |
| 2318 | ENSG0000( | 2.76E-07 | 1.33E-08 | -5.6826 | 0.1969 | -1.1189 | 104.5653 | FLNC |
| 64101 | ENSG0000( | 0.019099 | 0.003581 | -2.91286 | 0.664305 | -1.93503 | 9.104875 | LRRC4 |
| 3232 | ENSG0000( | 0.00258 | 0.000333 | -3.5883 | 0.546254 | -1.96012 | 14.20238 | HOXD3 |
| 84952 | ENSG0000( | 0.003843 | 0.000542 | -3.45911 | 0.466775 | -1.61463 | 18.12871 | CGNL1 |
| 8854 | ENSG0000( | 2.96E-33 | 9.51E-36 | -12.4807 | 0.181478 | -2.26498 | 130.7999 | ALDH1A2 |
| 3746 | ENSG0000( | 0.039191 | 0.008646 | -2.62574 | 0.451919 | -1.18662 | 17.9004 | KCNC1 |
| 57509 | ENSG0000( | 1.32E-09 | 4.29E-11 | -6.5938 | 0.170755 | -1.12593 | 126.3456 | MTUS1 |
| 11202 | ENSG0000( | 0.253171 | 0.098181 | 1.653736 | 0.657185 | 1.086812 | 8.911982 | KLK8 |
| 3169 | ENSG0000( | 0.000402 | 3.99E-05 | -4.10831 | 0.373315 | -1.53369 | 26.83002 | FOXA1 |
| 1036 | ENSG0000( | 0.002123 | 0.000267 | -3.64536 | 0.82999 | -3.02562 | 7.863677 | CDO1 |
| 9459 | ENSG0000( | 0.00031 | 2.96E-05 | -4.17634 | 0.360644 | -1.50617 | 30.62817 | ARHGEF6 |
| 8226 | ENSG0000( | 0.002327 | 0.000296 | -3.61875 | 0.444266 | -1.60769 | 18.88256 | PUDP |
| 27112 | ENSG0000( | 0.001392 | 0.000164 | -3.76959 | 0.490111 | -1.84752 | 17.35975 | FAM155B |
| 54857 | ENSG0000( | 0.389779 | 0.184205 | 1.327917 | 1.147731 | 1.524092 | 2.925966 | GDPD2 |
| 57631 | ENSG0000( | 7.57E-11 | 2.04E-12 | -7.03145 | 0.244813 | -1.72139 | 66.84955 | LRCH2 |
| 7837 | ENSG0000( | 5.07E-10 | 1.57E-11 | -6.74111 | 0.218483 | -1.47282 | 84.85608 | PXDN |
| 80726 | ENSG0000( | 0.504525 | 0.271233 | 1.100228 | 1.021158 | 1.123507 | 3.622559 | IQCN |
| 25830 | ENSG0000( | 0.005622 | 0.000848 | -3.33676 | 0.547802 | -1.82788 | 13.11562 | SULT4A1 |
| 1289 | ENSG0000( | 0.000732 | 7.86E-05 | -3.94878 | 0.42401 | -1.67432 | 23.70815 | COL5A1 |
| 84929 | ENSG0000( | 0.010866 | 0.001841 | -3.1147 | 0.338763 | -1.05515 | 30.28974 | FIBCD1 |
| 27293 | ENSG0000( | 0.003899 | 0.000551 | -3.45453 | 0.443402 | -1.53175 | 20.02512 | SMPDL3B |
| 2686 | ENSG0000( | 1.81E-07 | 8.27E-09 | -5.76293 | 0.24548 | -1.41469 | 61.71229 | GGT7 |
| 60401 | ENSG0000( | 0.026048 | 0.00523 | -2.79251 | 0.608939 | -1.70047 | 11.04379 | EDA2R |
| 6451 | ENSG0000( | 1.55E-12 | 3.23E-14 | -7.58885 | 0.235323 | -1.78583 | 70.64654 | SH3BGRL |
| 6569 | ENSG0000( | 0.436286 | 0.217814 | 1.232363 | 1.010407 | 1.245189 | 3.80322 | SLC34A1 |
| 23180 | ENSG0000( | 0.007274 | 0.00115 | -3.25105 | 0.717519 | -2.33269 | 8.605767 | RFTN1 |
| 57343 | ENSG0000( | 1.17E-05 | 7.71E-07 | -4.94248 | 0.389514 | -1.92517 | 26.66057 | ZNF304 |

| | | | | | | | | |
|---|---|---|---|---|---|---|---|---|
| 7568 | ENSG0000( | 0.005521 | 0.000831 | -3.34235 | 0.780056 | -2.60722 | 9.915457 | ZNF20 |
| 6538 | ENSG0000( | 0.006027 | 0.000923 | -3.31306 | 0.546518 | -1.81065 | 13.85488 | SLC6A11 |
| 1644 | ENSG0000( | 0.001997 | 0.000248 | 3.663955 | 0.669316 | 2.452343 | 11.05143 | DDC |
| 59352 | ENSG0000( | 3.69E-18 | 3.95E-20 | 9.189476 | 0.139095 | 1.278213 | 208.3351 | LGR6 |
| 23111 | ENSG0000( | 2.00E-23 | 1.30E-25 | -10.4616 | 0.181473 | -1.8985 | 126.5951 | SPART |
| 9365 | ENSG0000( | 0.406258 | 0.195204 | 1.295337 | 1.163143 | 1.506662 | 2.918698 | KL |
| 8471 | ENSG0000( | 2.14E-60 | 1.75E-63 | -16.8197 | 0.090994 | -1.5305 | 652.1117 | IRS4 |
| 55859 | ENSG0000( | 2.98E-10 | 8.92E-12 | -6.82293 | 0.292079 | -1.99284 | 47.6239 | BEX1 |
| 84626 | ENSG0000( | 1.99E-05 | 1.38E-06 | -4.82828 | 0.310523 | -1.49929 | 38.40611 | KRBA1 |
| 83857 | ENSG0000( | 0.000691 | 7.36E-05 | -3.96418 | 0.258115 | -1.02322 | 53.66502 | TMTC1 |
| 2947 | ENSG0000( | 6.56E-10 | 2.05E-11 | -6.70232 | 0.267135 | -1.79043 | 57.73829 | GSTM3 |
| 10451 | ENSG0000( | 0.039753 | 0.008823 | -2.61885 | 0.942979 | -2.46952 | 5.571986 | VAV3 |
| 3601 | ENSG0000( | 3.39E-11 | 8.73E-13 | 7.149224 | 0.148292 | 1.060172 | 171.1137 | IL15RA |
| 6660 | ENSG0000( | 0.031482 | 0.006603 | -2.71624 | 0.745487 | -2.02492 | 8.017379 | SOX5 |
| 85004 | ENSG0000( | 0.030814 | 0.006422 | -2.72543 | 0.804471 | -2.19253 | 6.472452 | RERG |
| 8933 | ENSG0000( | 2.89E-17 | 3.35E-19 | -8.9567 | 0.184869 | -1.65582 | 114.2011 | RTL8C |
| 1825 | ENSG0000( | 2.32E-24 | 1.33E-26 | -10.6751 | 0.146929 | -1.56849 | 182.2447 | DSC3 |
| 1837 | ENSG0000( | 9.63E-10 | 3.06E-11 | -6.6436 | 0.212726 | -1.41327 | 84.84941 | DTNA |
| 187 | ENSG0000( | 0.264293 | 0.104241 | 1.624634 | 1.250611 | 2.031785 | 2.886722 | APLNR |
| 22873 | ENSG0000( | 4.34E-16 | 5.52E-18 | -8.64214 | 0.18433 | -1.593 | 117.4668 | DZIP1 |
| 11095 | ENSG0000( | 3.05E-14 | 4.65E-16 | 8.120316 | 0.146609 | 1.190513 | 192.6492 | ADAMTS8 |
| 4440 | ENSG0000( | 4.87E-08 | 2.02E-09 | -5.99623 | 0.246966 | -1.48086 | 61.64516 | MSI1 |
| 8739 | ENSG0000( | 0.00328 | 0.000448 | -3.50977 | 0.450347 | -1.58061 | 18.47742 | HRK |
| 29969 | ENSG0000( | 5.16E-09 | 1.80E-10 | -6.3775 | 0.30378 | -1.93736 | 47.32456 | MDFIC |
| 22881 | ENSG0000( | 0.000443 | 4.45E-05 | -4.08269 | 0.38079 | -1.55465 | 26.614 | ANKRD6 |
| 2045 | ENSG0000( | 7.67E-12 | 1.80E-13 | -7.36257 | 0.233797 | -1.72135 | 71.07116 | EPHA7 |
| 116986 | ENSG0000( | 0.016173 | 0.002936 | -2.97437 | 0.442377 | -1.31579 | 18.79291 | AGAP2 |
| 5502 | ENSG0000( | 0.049925 | 0.011756 | -2.51938 | 0.728845 | -1.83624 | 7.272781 | PPP1R1A |
| 9053 | ENSG0000( | 7.68E-08 | 3.31E-09 | -5.91538 | 0.218797 | -1.29427 | 79.3949 | MAP7 |
| 5570 | ENSG0000( | 0.035832 | 0.00774 | -2.66319 | 0.490787 | -1.30706 | 15.14858 | PKIB |
| 27345 | ENSG0000( | 0.003597 | 0.000501 | -3.48026 | 0.307344 | -1.06964 | 37.91039 | KCNMB4 |
| 5077 | ENSG0000( | 0.000274 | 2.58E-05 | -4.20803 | 0.440604 | -1.85408 | 21.18012 | PAX3 |
| 80326 | ENSG0000( | 0.036635 | 0.007945 | 2.654381 | 0.483503 | 1.283402 | 17.4709 | WNT10A |
| 10913 | ENSG0000( | 1.04E-92 | 2.61E-96 | 20.82426 | 0.078913 | 1.643296 | 720.2233 | EDAR |
| 9771 | ENSG0000( | 9.55E-06 | 6.15E-07 | -4.98643 | 0.305496 | -1.52333 | 43.50433 | RAPGEF5 |
| 7080 | ENSG0000( | 0.036573 | 0.007926 | -2.65522 | 0.654586 | -1.73807 | 8.899013 | NKX2-1 |
| 2626 | ENSG0000( | 3.17E-06 | 1.85E-07 | -5.21362 | 0.310623 | -1.61947 | 40.52074 | GATA4 |
| 51759 | ENSG0000( | 4.76E-51 | 5.99E-54 | -15.4649 | 0.068824 | -1.06435 | 844.4339 | C9orf78 |
| 5168 | ENSG0000( | 0.042227 | 0.009521 | -2.59277 | 0.592328 | -1.53577 | 10.64138 | ENPP2 |
| 84709 | ENSG0000( | 0.020212 | 0.003851 | -2.8901 | 0.601293 | -1.7378 | 11.33902 | MGARP |
| 23213 | ENSG0000( | 0.011222 | 0.001914 | -3.10322 | 0.514695 | -1.59721 | 15.75528 | SULF1 |
| 4091 | ENSG0000( | 2.16E-22 | 1.59E-24 | -10.2214 | 0.118341 | -1.20961 | 276.7237 | SMAD6 |
| 80031 | ENSG0000( | 0.007401 | 0.001174 | -3.24523 | 0.511984 | -1.66151 | 14.47387 | SEMA6D |
| 51196 | ENSG0000( | 0.000473 | 4.80E-05 | -4.06538 | 0.247445 | -1.00596 | 58.32572 | PLCE1 |
| 57520 | ENSG0000( | 0.035688 | 0.007703 | -2.66484 | 0.519965 | -1.38562 | 14.43031 | HECW2 |
| 30061 | ENSG0000( | 0.042198 | 0.009509 | -2.59321 | 0.459156 | -1.19069 | 17.09053 | SLC40A1 |
| 10144 | ENSG0000( | 8.98E-09 | 3.26E-10 | -6.2859 | 0.298975 | -1.87933 | 48.41782 | FAM13A |
| 51191 | ENSG0000( | 1.26E-07 | 5.62E-09 | -5.82778 | 0.315799 | -1.84041 | 42.21606 | HERC5 |
| 57575 | ENSG0000( | 3.28E-19 | 3.16E-21 | -9.45733 | 0.183299 | -1.73352 | 121.7603 | PCDH10 |
| 2247 | ENSG0000( | 0.001111 | 0.000126 | -3.83346 | 0.334022 | -1.28046 | 32.97577 | FGF2 |

| | | | | | | | | |
|---|---|---|---|---|---|---|---|---|
| 8654 | ENSG00000( | 0.002611 | 0.000339 | -3.58353 | 0.344811 | -1.23564 | 32.06692 | PDE5A |
| 901 | ENSG00000( | 6.54E-11 | 1.74E-12 | -7.05415 | 0.17123 | -1.20788 | 125.6699 | CCNG2 |
| 2201 | ENSG00000( | 3.69E-17 | 4.37E-19 | -8.92726 | 0.194104 | -1.73281 | 107.4244 | FBN2 |
| 144402 | ENSG00000( | 3.61E-06 | 2.14E-07 | -5.18712 | 0.322877 | -1.6748 | 39.10047 | CPNE8 |
| 144165 | ENSG00000( | 0.004896 | 0.00072 | -3.38199 | 0.373444 | -1.26298 | 29.6025 | PRICKLE1 |
| 1280 | ENSG00000( | 3.44E-08 | 1.38E-09 | -6.0573 | 0.241479 | -1.46271 | 66.46881 | COL2A1 |
| 121227 | ENSG00000( | 0.001096 | 0.000124 | -3.83791 | 0.341319 | -1.30995 | 33.85505 | LRIG3 |
| 122060 | ENSG00000( | 1.68E-24 | 9.19E-27 | -10.7095 | 0.16853 | -1.80487 | 157.1242 | SLAIN1 |
| 4857 | ENSG00000( | 0.000746 | 8.03E-05 | -3.94359 | 0.467033 | -1.84179 | 17.94832 | NOVA1 |
| 145407 | ENSG00000( | 0.034981 | 0.007517 | -2.67303 | 0.432825 | -1.15695 | 20.06691 | ARMH4 |
| 1583 | ENSG00000( | 0.333425 | 0.146067 | 1.453564 | 1.133523 | 1.647648 | 3.146866 | CYP11A1 |
| 145864 | ENSG00000( | 3.03E-06 | 1.76E-07 | -5.22277 | 0.360267 | -1.88159 | 32.94195 | HAPLN3 |
| 6457 | ENSG00000( | 2.10E-06 | 1.17E-07 | -5.29877 | 0.485358 | -2.5718 | 20.40118 | SH3GL3 |
| 1009 | ENSG00000( | 0.26286 | 0.103394 | 1.628618 | 0.669453 | 1.090283 | 8.960036 | CDH11 |
| 64762 | ENSG00000( | 0.003901 | 0.000552 | -3.45423 | 0.32676 | -1.12871 | 34.3099 | GAREM1 |
| 80000 | ENSG00000( | 2.68E-12 | 5.83E-14 | -7.51191 | 0.242966 | -1.82514 | 74.83408 | GREB1L |
| 84225 | ENSG00000( | 0.101472 | 0.028995 | 2.183552 | 0.517215 | 1.129366 | 14.51477 | ZMYND15 |
| 5596 | ENSG00000( | 0.048256 | 0.011271 | -2.53417 | 1.185461 | -3.00416 | 3.843044 | MAPK4 |
| 4089 | ENSG00000( | 5.02E-13 | 9.48E-15 | -7.74611 | 0.229321 | -1.77634 | 77.40567 | SMAD4 |
| 8792 | ENSG00000( | 0.019189 | 0.003603 | -2.91097 | 0.388403 | -1.13063 | 23.19271 | TNFRSF11A |
| 4542 | ENSG00000( | 3.25E-17 | 3.81E-19 | 8.94247 | 0.221479 | 1.980566 | 89.07133 | MYO1F |
| 83872 | ENSG00000( | 0.021642 | 0.004185 | -2.86389 | 0.94875 | -2.71711 | 5.497902 | HMCN1 |
| 57198 | ENSG00000( | 6.75E-14 | 1.09E-15 | -8.01616 | 0.127571 | -1.02263 | 227.5944 | ATP8B2 |
| 151449 | ENSG00000( | 0.023822 | 0.004701 | -2.82684 | 0.439292 | -1.24181 | 19.72383 | GDF7 |
| 165545 | ENSG00000( | 0.178017 | 0.060948 | 1.873871 | 1.421239 | 2.663219 | 2.852438 | DQX1 |
| 114805 | ENSG00000( | 3.63E-06 | 2.15E-07 | -5.18606 | 0.372927 | -1.93402 | 28.61572 | GALNT13 |
| 339768 | ENSG00000( | 0.011887 | 0.002043 | -3.08385 | 0.462439 | -1.42609 | 17.71092 | ESPNL |
| 57574 | ENSG00000( | 6.80E-08 | 2.90E-09 | -5.93742 | 0.476675 | -2.83022 | 22.47077 | MARCHF4 |
| 27303 | ENSG00000( | 0.004495 | 0.000651 | -3.40942 | 0.56473 | -1.9254 | 12.68113 | RBMS3 |
| 729085 | ENSG00000( | 0.018489 | 0.003447 | -2.92476 | 0.633923 | -1.85407 | 10.13296 | GASK1A |
| 23066 | ENSG00000( | 4.11E-08 | 1.68E-09 | -6.02581 | 0.288763 | -1.74003 | 46.55166 | CAND2 |
| 66000 | ENSG00000( | 0.033764 | 0.007203 | -2.68732 | 0.729231 | -1.95967 | 8.25916 | TMEM108 |
| 9353 | ENSG00000( | 1.16E-52 | 1.38E-55 | -15.7058 | 0.124475 | -1.95497 | 274.4481 | SLIT2 |
| 6622 | ENSG00000( | 0.01397 | 0.002469 | -3.02708 | 0.393475 | -1.19108 | 25.19673 | SNCA |
| 56034 | ENSG00000( | 1.06E-05 | 6.92E-07 | -4.96362 | 0.290916 | -1.44399 | 44.93029 | PDGFC |
| 23635 | ENSG00000( | 9.82E-07 | 5.14E-08 | -5.44644 | 0.309268 | -1.68441 | 40.86793 | SSBP2 |
| 91975 | ENSG00000( | 0.002626 | 0.000341 | -3.5818 | 0.452595 | -1.6211 | 20.16328 | ZNF300 |
| 23500 | ENSG00000( | 0.000125 | 1.07E-05 | -4.40302 | 0.421746 | -1.85695 | 22.65818 | DAAM2 |
| 6581 | ENSG00000( | 2.59E-14 | 3.89E-16 | 8.142 | 0.15661 | 1.275117 | 176.9487 | SLC22A3 |
| 221806 | ENSG00000( | 0.0011 | 0.000125 | -3.83665 | 0.588477 | -2.25778 | 12.64154 | VWDE |
| 9586 | ENSG00000( | 0.000106 | 8.85E-06 | -4.44344 | 0.312958 | -1.39061 | 40.86992 | CREB5 |
| 3561 | ENSG00000( | 0.000168 | 1.48E-05 | 4.330929 | 0.388774 | 1.683752 | 27.12801 | IL2RG |
| 7552 | ENSG00000( | 6.69E-11 | 1.79E-12 | -7.05027 | 0.219011 | -1.54409 | 92.40799 | ZNF711 |
| 55086 | ENSG00000( | 1.53E-25 | 7.68E-28 | -10.9369 | 0.153995 | -1.68423 | 163.5616 | RADX |
| 84443 | ENSG00000( | 0.019748 | 0.003737 | -2.89955 | 0.795833 | -2.30756 | 6.794892 | FRMPD3 |
| 139818 | ENSG00000( | 7.38E-14 | 1.21E-15 | 8.003745 | 0.136661 | 1.093802 | 202.4945 | DOCK11 |
| 2719 | ENSG00000( | 5.50E-13 | 1.05E-14 | -7.73323 | 0.282463 | -2.18435 | 52.90751 | GPC3 |
| 2040 | ENSG00000( | 6.89E-08 | 2.96E-09 | -5.93389 | 0.240115 | -1.42482 | 63.60026 | STOM |
| 92211 | ENSG00000( | 1.05E-06 | 5.55E-08 | -5.43272 | 0.389597 | -2.11657 | 29.20573 | CDHR1 |
| 9118 | ENSG00000( | 6.21E-06 | 3.86E-07 | -5.07559 | 0.334932 | -1.69998 | 39.00062 | INA |

| | | | | | | | | |
|---|---|---|---|---|---|---|---|---|
| 10205 | ENSG0000( | 4.42E-89 | 1.39E-92 | 20.40902 | 0.056912 | 1.161523 | 1290.473 | MPZL2 |
| 57158 | ENSG0000( | 0.001495 | 0.000177 | -3.74945 | 0.311876 | -1.16936 | 37.54634 | JPH2 |
| 64405 | ENSG0000( | 0.45904 | 0.234413 | 1.189068 | 1.130017 | 1.343667 | 3.398067 | CDH22 |
| 23284 | ENSG0000( | 0.048705 | 0.011398 | -2.53026 | 0.443985 | -1.1234 | 17.95943 | ADGRL3 |
| 116372 | ENSG0000( | 0.000494 | 5.06E-05 | -4.05282 | 0.495508 | -2.0082 | 20.11754 | LYPD1 |
| 2823 | ENSG0000( | 0.001706 | 0.000206 | -3.71108 | 1.040824 | -3.86258 | 6.672535 | GPM6A |
| 116966 | ENSG0000( | 0.019227 | 0.003613 | -2.91014 | 0.385523 | -1.12193 | 24.81652 | WDR17 |
| 341640 | ENSG0000( | 2.96E-06 | 1.72E-07 | -5.2278 | 0.270964 | -1.41654 | 49.7611 | FREM2 |
| 7068 | ENSG0000( | 0.000391 | 3.85E-05 | -4.11605 | 0.416002 | -1.71228 | 25.41184 | THRB |
| 1909 | ENSG0000( | 0.02173 | 0.004209 | -2.86208 | 0.592741 | -1.69647 | 12.14886 | EDNRA |
| 4306 | ENSG0000( | 0.018133 | 0.003364 | -2.93238 | 0.584681 | -1.71451 | 11.78686 | NR3C2 |
| 2313 | ENSG0000( | 0.005723 | 0.000865 | -3.33112 | 0.512952 | -1.70871 | 14.75965 | FLI1 |
| 148534 | ENSG0000( | 8.73E-05 | 7.10E-06 | -4.49081 | 0.305239 | -1.37077 | 41.45215 | TLCD4 |
| 5457 | ENSG0000( | 1.21E-14 | 1.78E-16 | -8.23614 | 0.205028 | -1.68864 | 99.42505 | POU4F1 |
| 2977 | ENSG0000( | 0.030496 | 0.006346 | -2.72935 | 0.86417 | -2.35862 | 6.068668 | GUCY1A2 |
| 2697 | ENSG0000( | 2.80E-18 | 2.92E-20 | -9.22169 | 0.182172 | -1.67993 | 116.4179 | GJA1 |
| 5352 | ENSG0000( | 3.10E-25 | 1.62E-27 | -10.869 | 0.161427 | -1.75455 | 153.5112 | PLOD2 |
| 55203 | ENSG0000( | 0.008881 | 0.00145 | -3.18447 | 0.559187 | -1.78071 | 12.67429 | LGI2 |
| 22925 | ENSG0000( | 0.028846 | 0.005919 | 2.752225 | 1.59905 | 4.400945 | 2.217757 | PLA2R1 |
| 5789 | ENSG0000( | 0.000132 | 1.13E-05 | -4.39115 | 0.428139 | -1.88002 | 21.98576 | PTPRD |
| 781 | ENSG0000( | 3.11E-06 | 1.81E-07 | -5.21786 | 0.324891 | -1.69524 | 36.55843 | CACNA2D1 |
| 223117 | ENSG0000( | 0.002686 | 0.000351 | -3.57464 | 0.462147 | -1.65201 | 17.99502 | SEMA3D |
| 125206 | ENSG0000( | 0.42378 | 0.208077 | 1.258872 | 0.896164 | 1.128155 | 4.851299 | SLC5A10 |
| 26289 | ENSG0000( | 4.67E-08 | 1.93E-09 | -6.0039 | 0.295389 | -1.77349 | 45.71752 | AK5 |
| 54538 | ENSG0000( | 0.510278 | 0.275628 | 1.090193 | 1.009223 | 1.100248 | 3.58728 | ROBO4 |
| 64090 | ENSG0000( | 0.008897 | 0.001453 | 3.183849 | 0.440151 | 1.401375 | 20.69383 | GAL3ST2 |
| 7345 | ENSG0000( | 1.50E-49 | 2.17E-52 | -15.2319 | 0.116114 | -1.76864 | 317.7033 | UCHL1 |
| 728239 | ENSG0000( | 0.049462 | 0.011603 | -2.524 | 0.586977 | -1.48153 | 42.66355 | MAGED4 |
| 140578 | ENSG0000( | 0.039057 | 0.008614 | -2.627 | 0.910384 | -2.39158 | 5.456452 | CHODL |
| 9510 | ENSG0000( | 0.000143 | 1.24E-05 | -4.37028 | 0.408305 | -1.78441 | 24.08289 | ADAMTS1 |
| 23228 | ENSG0000( | 0.001503 | 0.000178 | -3.74774 | 0.32466 | -1.21674 | 36.0686 | PLCL2 |
| 51560 | ENSG0000( | 0.00157 | 0.000188 | -3.73498 | 0.325609 | -1.21615 | 34.38117 | RAB6B |
| 56776 | ENSG0000( | 0.003338 | 0.000458 | -3.50429 | 0.504448 | -1.76773 | 15.40045 | FMN2 |
| 286097 | ENSG0000( | 0.000716 | 7.66E-05 | -3.95494 | 0.757835 | -2.99719 | 9.136323 | MICU3 |
| 23426 | ENSG0000( | 0.026073 | 0.005238 | -2.79199 | 0.520448 | -1.45309 | 14.34268 | GRIP1 |
| 4325 | ENSG0000( | 0.00134 | 0.000156 | -3.78079 | 0.495878 | -1.87481 | 16.97893 | MMP16 |
| 9508 | ENSG0000( | 0.000384 | 3.78E-05 | -4.1205 | 0.469452 | -1.93438 | 19.26554 | ADAMTS3 |
| 7102 | ENSG0000( | 0.034779 | 0.007465 | -2.67536 | 0.599353 | -1.60348 | 12.14963 | TSPAN7 |
| 6854 | ENSG0000( | 0.011139 | 0.001897 | -3.10596 | 0.528611 | -1.64184 | 13.69871 | SYN2 |
| 3815 | ENSG0000( | 0.000332 | 3.19E-05 | -4.15916 | 0.320122 | -1.33144 | 35.25411 | KIT |
| 90139 | ENSG0000( | 2.35E-05 | 1.65E-06 | -4.79243 | 0.44926 | -2.15304 | 22.00294 | TSPAN18 |
| 7780 | ENSG0000( | 0.150379 | 0.048467 | 1.973249 | 0.755441 | 1.490672 | 7.964492 | SLC30A2 |
| 11013 | ENSG0000( | 0.004254 | 0.00061 | -3.42718 | 0.603577 | -2.06857 | 11.95702 | TMSB15A |
| 22808 | ENSG0000( | 3.66E-06 | 2.17E-07 | -5.18428 | 0.294908 | -1.52889 | 46.46655 | MRAS |
| 81035 | ENSG0000( | 2.41E-06 | 1.36E-07 | -5.27009 | 0.394076 | -2.07682 | 29.47571 | COLEC12 |
| 653361 | ENSG0000( | 0.003838 | 0.000541 | 3.459697 | 0.84813 | 2.934272 | 8.239042 | NCF1 |
| 4359 | ENSG0000( | 4.67E-06 | 2.83E-07 | 5.134685 | 0.224093 | 1.150649 | 78.38937 | MPZ |
| 6493 | ENSG0000( | 0.000145 | 1.26E-05 | -4.36719 | 0.264938 | -1.15703 | 53.3577 | SIM2 |
| 79190 | ENSG0000( | 0.030926 | 0.006453 | -2.72383 | 0.753955 | -2.05364 | 7.554741 | IRX6 |
| 875 | ENSG0000( | 7.36E-22 | 5.70E-24 | -10.097 | 0.109932 | -1.10998 | 379.6124 | CBS |

| | | | | | | | | |
|---|---|---|---|---|---|---|---|---|
| 54742 | ENSG0000( | 0.186204 | 0.064513 | 1.848618 | 1.728113 | 3.19462 | 2.090134 | LY6K |
| 58191 | ENSG0000( | 2.50E-11 | 6.27E-13 | 7.194525 | 0.143157 | 1.029944 | 184.2088 | CXCL16 |
| 9965 | ENSG0000( | 6.62E-27 | 3.12E-29 | 11.22357 | 0.118402 | 1.328889 | 277.8957 | FGF19 |
| 22977 | ENSG0000( | 0.117985 | 0.035235 | 2.105647 | 0.499462 | 1.05169 | 14.52214 | AKR7A3 |
| 10630 | ENSG0000( | 4.44E-11 | 1.15E-12 | -7.11068 | 0.312728 | -2.22371 | 43.67065 | PDPN |
| 9077 | ENSG0000( | 0.488794 | 0.258761 | 1.129325 | 1.08058 | 1.220326 | 3.175492 | DIRAS3 |
| 343450 | ENSG0000( | 0.026277 | 0.005291 | -2.78875 | 0.636749 | -1.77573 | 9.550798 | KCNT2 |
| 4921 | ENSG0000( | 2.64E-14 | 4.00E-16 | -8.13864 | 0.212125 | -1.72641 | 86.36694 | DDR2 |
| 57216 | ENSG0000( | 5.20E-05 | 3.99E-06 | -4.61213 | 0.265553 | -1.22476 | 50.89794 | VANGL2 |
| 79762 | ENSG0000( | 9.01E-07 | 4.69E-08 | -5.46281 | 0.371968 | -2.03199 | 30.70873 | C1orf115 |
| 132671 | ENSG0000( | 0.022768 | 0.004454 | -2.84405 | 0.717876 | -2.04168 | 8.664245 | SPATA18 |
| 1520 | ENSG0000( | 1.75E-09 | 5.78E-11 | 6.549215 | 0.409333 | 2.680811 | 34.82109 | CTSS |
| 317649 | ENSG0000( | 7.79E-05 | 6.26E-06 | -4.51729 | 0.259298 | -1.17132 | 58.73917 | EIF4E3 |
| 11167 | ENSG0000( | 5.02E-38 | 1.17E-40 | -13.3511 | 0.139494 | -1.86239 | 208.7112 | FSTL1 |
| 25802 | ENSG0000( | 0.01167 | 0.002003 | -3.08974 | 0.994208 | -3.07185 | 5.553796 | LMOD1 |
| 166336 | ENSG0000( | 0.008904 | 0.001455 | -3.18349 | 0.44327 | -1.41115 | 19.47222 | PRICKLE2 |
| 5806 | ENSG0000( | 0.020889 | 0.004014 | -2.87705 | 0.944968 | -2.71872 | 5.444504 | PTX3 |
| 54502 | ENSG0000( | 0.000568 | 5.89E-05 | -4.01736 | 0.387831 | -1.55806 | 25.81588 | RBM47 |
| 2919 | ENSG0000( | 0.1036 | 0.029785 | 2.172934 | 0.571538 | 1.241915 | 12.07882 | CXCL1 |
| 116832 | ENSG0000( | 0.006027 | 0.000923 | -3.31298 | 0.431127 | -1.42831 | 20.20118 | RPL39L |
| 9464 | ENSG0000( | 0.018756 | 0.003506 | -2.91946 | 0.886768 | -2.58889 | 6.150228 | HAND2 |
| 79884 | ENSG0000( | 2.75E-12 | 5.99E-14 | -7.50822 | 0.235849 | -1.77081 | 70.61899 | MAP9 |
| 2982 | ENSG0000( | 0.047123 | 0.01093 | -2.54494 | 0.480559 | -1.22299 | 17.14863 | GUCY1A1 |
| 55314 | ENSG0000( | 0.001448 | 0.000171 | -3.75802 | 0.442724 | -1.66377 | 20.22539 | TMEM144 |
| 10085 | ENSG0000( | 4.58E-06 | 2.76E-07 | -5.13914 | 0.355992 | -1.8295 | 32.8758 | EDIL3 |
| 84059 | ENSG0000( | 0.002042 | 0.000255 | -3.65761 | 0.423427 | -1.54873 | 22.75522 | ADGRV1 |
| 651746 | ENSG0000( | 0.02111 | 0.004066 | -2.87299 | 0.451101 | -1.29601 | 17.5576 | ANKRD33B |
| 133396 | ENSG0000( | 0.454971 | 0.231551 | 1.196374 | 0.842221 | 1.007611 | 5.134725 | IL31RA |
| 221833 | ENSG0000( | 0.004586 | 0.000668 | -3.40248 | 0.616506 | -2.09765 | 10.7275 | SP8 |
| 23462 | ENSG0000( | 1.21E-54 | 1.22E-57 | -16.003 | 0.102062 | -1.63329 | 393.9261 | HEY1 |
| 6469 | ENSG0000( | 1.11E-08 | 4.09E-10 | 6.25058 | 0.221773 | 1.386212 | 78.73504 | SHH |
| 107 | ENSG0000( | 6.56E-09 | 2.33E-10 | -6.33808 | 0.321858 | -2.03996 | 44.04341 | ADCY1 |
| 157753 | ENSG0000( | 0.039673 | 0.008797 | -2.61983 | 0.734147 | -1.92334 | 7.673755 | TMEM74 |
| 216 | ENSG0000( | 0.011061 | 0.00188 | -3.10857 | 1.454612 | -4.52177 | 3.736375 | ALDH1A1 |
| 158158 | ENSG0000( | 3.89E-09 | 1.34E-10 | -6.42259 | 0.244769 | -1.57205 | 72.58254 | RASEF |
| 26050 | ENSG0000( | 7.38E-14 | 1.21E-15 | -8.0032 | 0.216585 | -1.73337 | 91.71585 | SLITRK5 |
| 120114 | ENSG0000( | 0.004697 | 0.000685 | -3.39536 | 0.689598 | -2.34143 | 9.51981 | FAT3 |
| 84889 | ENSG0000( | 0.009876 | 0.001637 | -3.14915 | 0.683851 | -2.15355 | 10.10208 | SLC7A3 |
| 63876 | ENSG0000( | 0.003673 | 0.000513 | -3.47395 | 0.491615 | -1.70784 | 16.95643 | PKNOX2 |
| 83592 | ENSG0000( | 0.121191 | 0.036444 | 2.091938 | 0.526502 | 1.101409 | 14.86075 | AKR1E2 |
| 196740 | ENSG0000( | 0.045617 | 0.010486 | -2.55939 | 0.644308 | -1.64903 | 9.414975 | VSTM4 |
| 143379 | ENSG0000( | 0.007077 | 0.001112 | -3.26065 | 0.547013 | -1.78362 | 12.96522 | C10orf82 |
| 79841 | ENSG0000( | 0.131514 | 0.040467 | 2.048949 | 0.510001 | 1.044966 | 15.05956 | AGBL2 |
| 341346 | ENSG0000( | 7.57E-11 | 2.04E-12 | 7.031502 | 0.253441 | 1.782072 | 66.30833 | SMCO2 |
| 79789 | ENSG0000( | 0.020036 | 0.003806 | -2.89379 | 0.377302 | -1.09183 | 25.93632 | CLMN |
| 154810 | ENSG0000( | 6.64E-18 | 7.31E-20 | -9.12293 | 0.184905 | -1.68687 | 114.5314 | AMOTL1 |
| 255394 | ENSG0000( | 0.015172 | 0.002725 | -2.99721 | 0.602548 | -1.80596 | 11.21601 | TCP11L2 |
| 83700 | ENSG0000( | 2.85E-13 | 5.21E-15 | -7.82167 | 0.172832 | -1.35183 | 130.737 | JAM3 |
| 112937 | ENSG0000( | 0.0011 | 0.000125 | -3.83646 | 0.547945 | -2.10217 | 16.76135 | GLB1L3 |
| 81693 | ENSG0000( | 0.233919 | 0.087963 | 1.706245 | 0.749868 | 1.279459 | 7.222792 | AMN |

| | | | | | | | | |
|---|---|---|---|---|---|---|---|---|
| 2200 | ENSG0000( | 3.22E-05 | 2.33E-06 | -4.72289 | 0.280641 | -1.32544 | 45.80623 | FBN1 |
| 654429 | ENSG0000( | 0.491238 | 0.260642 | 1.124875 | 0.973461 | 1.095022 | 4.149463 | LRTM2 |
| 1152 | ENSG0000( | 1.74E-144 | 2.20E-148 | -25.9427 | 0.051951 | -1.34775 | 2021.888 | CKB |
| 7275 | ENSG0000( | 1.81E-18 | 1.84E-20 | -9.27119 | 0.187267 | -1.73619 | 113.2558 | TUB |
| 283659 | ENSG0000( | 1.48E-08 | 5.56E-10 | -6.20244 | 0.270901 | -1.68025 | 52.77658 | PRTG |
| 23423 | ENSG0000( | 1.88E-19 | 1.78E-21 | -9.51729 | 0.172401 | -1.64079 | 133.1255 | TMED3 |
| 3948 | ENSG0000( | 0.003064 | 0.000413 | -3.5316 | 0.655611 | -2.31536 | 10.51609 | LDHC |
| 6447 | ENSG0000( | 0.313925 | 0.133515 | 1.500384 | 0.949521 | 1.424646 | 4.188997 | SCG5 |
| 4130 | ENSG0000( | 7.74E-10 | 2.45E-11 | -6.67631 | 0.218188 | -1.45669 | 80.47397 | MAP1A |
| 64127 | ENSG0000( | 0.031644 | 0.006645 | 2.71415 | 0.390271 | 1.059255 | 26.46395 | NOD2 |
| 94274 | ENSG0000( | 0.004193 | 0.0006 | -3.43173 | 0.626478 | -2.1499 | 10.84825 | PPP1R14A |
| 21 | ENSG0000( | 8.24E-08 | 3.57E-09 | -5.90307 | 0.176293 | -1.04067 | 115.0546 | ABCA3 |
| 146434 | ENSG0000( | 0.025823 | 0.005175 | -2.79591 | 0.735741 | -2.05707 | 7.486246 | ZNF597 |
| 222584 | ENSG0000( | 0.021875 | 0.004242 | -2.85955 | 0.82472 | -2.35833 | 6.956271 | FAM83B |
| 1281 | ENSG0000( | 0.031903 | 0.006719 | -2.71045 | 0.633201 | -1.71626 | 14.1044 | COL3A1 |
| 167127 | ENSG0000( | 0.006618 | 0.001027 | -3.28292 | 0.678511 | -2.22749 | 10.23517 | UGT3A2 |
| 753 | ENSG0000( | 0.001805 | 0.00022 | -3.69449 | 0.444278 | -1.64138 | 19.99195 | LDLRAD4 |
| 53353 | ENSG0000( | 0.003073 | 0.000415 | -3.53064 | 0.722254 | -2.55002 | 9.118518 | LRP1B |
| 255743 | ENSG0000( | 0.023635 | 0.004651 | -2.83029 | 0.587678 | -1.6633 | 11.02048 | NPNT |
| 25849 | ENSG0000( | 0.002812 | 0.000371 | -3.55987 | 0.441269 | -1.57086 | 19.41201 | PARM1 |
| 90362 | ENSG0000( | 0.005562 | 0.000838 | -3.34006 | 0.581194 | -1.94122 | 11.90689 | FAM110B |
| 55130 | ENSG0000( | 0.037953 | 0.008308 | -2.63927 | 0.731612 | -1.93093 | 8.273392 | ARMC4 |
| 126669 | ENSG0000( | 0.551566 | 0.313579 | 1.00774 | 1.024919 | 1.032852 | 3.439601 | SHE |
| 6335 | ENSG0000( | 0.01801 | 0.003332 | -2.93533 | 0.564706 | -1.6576 | 12.32146 | SCN9A |
| 6383 | ENSG0000( | 4.52E-15 | 6.48E-17 | -8.35611 | 0.216629 | -1.81018 | 85.65551 | SDC2 |
| 83987 | ENSG0000( | 1.38E-09 | 4.52E-11 | -6.58579 | 0.242858 | -1.59941 | 65.14421 | CCDC8 |
| 9839 | ENSG0000( | 3.63E-05 | 2.66E-06 | -4.69559 | 0.354312 | -1.6637 | 33.10427 | ZEB2 |
| 123591 | ENSG0000( | 0.024312 | 0.00482 | -2.8188 | 0.861855 | -2.42939 | 6.529637 | TMEM266 |
| 5099 | ENSG0000( | 8.26E-29 | 3.32E-31 | -11.6183 | 0.161532 | -1.87673 | 157.1188 | PCDH7 |
| 5028 | ENSG0000( | 2.71E-07 | 1.30E-08 | -5.68632 | 0.367441 | -2.08939 | 30.59779 | P2RY1 |
| 161725 | ENSG0000( | 0.003391 | 0.000467 | -3.49927 | 0.688689 | -2.40991 | 9.939378 | OTUD7A |
| 214 | ENSG0000( | 6.75E-19 | 6.63E-21 | -9.37952 | 0.169944 | -1.594 | 135.2663 | ALCAM |
| 57484 | ENSG0000( | 4.77E-07 | 2.37E-08 | -5.58235 | 0.352144 | -1.96579 | 32.79723 | RNF150 |
| 9720 | ENSG0000( | 0.166308 | 0.055579 | 1.914323 | 0.909045 | 1.740207 | 4.942749 | CCDC144A |
| 166614 | ENSG0000( | 0.010298 | 0.001727 | -3.13351 | 0.454871 | -1.42534 | 18.22791 | DCLK2 |
| 164832 | ENSG0000( | 4.64E-06 | 2.80E-07 | -5.13616 | 0.214122 | -1.09977 | 78.48609 | LONRF2 |
| 1000 | ENSG0000( | 1.33E-34 | 4.01E-37 | -12.7305 | 0.150513 | -1.9161 | 179.123 | CDH2 |
| 133418 | ENSG0000( | 3.03E-21 | 2.48E-23 | -9.95172 | 0.195052 | -1.9411 | 108.2716 | EMB |
| 10736 | ENSG0000( | 2.07E-05 | 1.44E-06 | -4.81995 | 0.226119 | -1.08988 | 69.61067 | SIX2 |
| 2850 | ENSG0000( | 1.05E-09 | 3.37E-11 | -6.62938 | 0.312143 | -2.06931 | 44.0784 | GPR27 |
| 10761 | ENSG0000( | 0.028562 | 0.005848 | -2.75617 | 0.461851 | -1.27294 | 18.3439 | PLAC1 |
| 90161 | ENSG0000( | 1.93E-16 | 2.39E-18 | -8.73725 | 0.188322 | -1.64541 | 126.4479 | HS6ST2 |
| 26108 | ENSG0000( | 2.00E-05 | 1.38E-06 | -4.82717 | 0.291821 | -1.40867 | 42.50977 | PYGO1 |
| 5569 | ENSG0000( | 0.004011 | 0.000569 | -3.44577 | 0.436133 | -1.50281 | 19.88812 | PKIA |
| 116844 | ENSG0000( | 0.11681 | 0.034737 | 2.111408 | 1.572014 | 3.319163 | 2.218828 | LRG1 |
| 4884 | ENSG0000( | 6.78E-07 | 3.45E-08 | -5.51671 | 0.198248 | -1.09368 | 92.78458 | NPTX1 |
| 3777 | ENSG0000( | 0.430622 | 0.213468 | 1.244084 | 0.849417 | 1.056747 | 5.279 | KCNK3 |
| 23566 | ENSG0000( | 0.03729 | 0.008127 | -2.64675 | 0.390909 | -1.03464 | 24.98231 | LPAR3 |
| 9427 | ENSG0000( | 0.010847 | 0.001837 | -3.11544 | 0.495874 | -1.54486 | 15.28723 | ECEL1 |
| 203859 | ENSG0000( | 2.61E-08 | 1.03E-09 | -6.10408 | 0.275968 | -1.68453 | 50.17804 | ANO5 |

| | | | | | | | | |
|---|---|---|---|---|---|---|---|---|
| 10517 | ENSG0000( | 0.392135 | 0.18565 | 1.323558 | 1.649366 | 2.183032 | 2.101384 | FBXW10 |
| 27023 | ENSG0000( | 0.003692 | 0.000516 | -3.4721 | 0.66627 | -2.31336 | 10.24099 | FOXB1 |
| 3400 | ENSG0000( | 1.23E-23 | 7.74E-26 | -10.5104 | 0.166009 | -1.74483 | 146.7665 | ID4 |
| 7177 | ENSG0000( | 0.216648 | 0.079123 | 1.755796 | 0.840965 | 1.476563 | 6.422492 | TPSAB1 |
| 10231 | ENSG0000( | 0.036384 | 0.007878 | -2.65725 | 0.786073 | -2.08879 | 6.764116 | RCAN2 |
| 124221 | ENSG0000( | 0.00312 | 0.000423 | -3.52513 | 0.837739 | -2.95314 | 7.69182 | PRSS30P |
| 348938 | ENSG0000( | 0.032054 | 0.006765 | 2.708202 | 0.415928 | 1.126416 | 22.18998 | NIPAL4 |
| 90288 | ENSG0000( | 0.52299 | 0.286929 | 1.064884 | 1.132617 | 1.206105 | 3.096958 | EFCAB12 |
| 3222 | ENSG0000( | 0.048959 | 0.01146 | 2.528344 | 0.49228 | 1.244653 | 16.61115 | HOXC5 |
| 23491 | ENSG0000( | 0.00035 | 3.39E-05 | -4.14547 | 0.317058 | -1.31436 | 36.18028 | CES3 |
| 3642 | ENSG0000( | 0.039339 | 0.008691 | -2.62397 | 0.884847 | -2.32181 | 6.070624 | INSM1 |
| 5797 | ENSG0000( | 0.001144 | 0.00013 | -3.82575 | 0.413885 | -1.58342 | 21.92606 | PTPRM |
| 1464 | ENSG0000( | 0.009584 | 0.001583 | -3.15902 | 0.585699 | -1.85023 | 11.83327 | CSPG4 |
| 26032 | ENSG0000( | 2.81E-08 | 1.12E-09 | -6.09195 | 0.240874 | -1.46739 | 65.47478 | SUSD5 |
| 353189 | ENSG0000( | 0.049454 | 0.011589 | -2.5244 | 0.551929 | -1.39329 | 12.45168 | SLCO4C1 |
| 8722 | ENSG0000( | 0.001631 | 0.000196 | -3.72366 | 0.514028 | -1.91407 | 15.32632 | CTSF |
| 170261 | ENSG0000( | 3.26E-05 | 2.36E-06 | -4.71972 | 0.217434 | -1.02623 | 75.97964 | ZCCHC12 |
| 26047 | ENSG0000( | 0.000232 | 2.12E-05 | -4.2515 | 0.304818 | -1.29593 | 40.4115 | CNTNAP2 |
| 133022 | ENSG0000( | 0.031007 | 0.006476 | -2.72266 | 1.037165 | -2.82385 | 4.740613 | TRAM1L1 |
| 9435 | ENSG0000( | 1.30E-12 | 2.65E-14 | -7.6143 | 0.151285 | -1.15193 | 165.2548 | CHST2 |
| 147409 | ENSG0000( | 0.05292 | 0.012611 | 2.494577 | 0.808703 | 2.017373 | 6.719163 | DSG4 |
| 1674 | ENSG0000( | 0.164448 | 0.054688 | 1.921343 | 0.831889 | 1.598343 | 6.128855 | DES |
| 92369 | ENSG0000( | 0.014164 | 0.002511 | -3.02202 | 0.7355 | -2.22269 | 8.212889 | SPSB4 |
| 388403 | ENSG0000( | 0.027614 | 0.005623 | -2.769 | 0.444274 | -1.23019 | 19.12838 | YPEL2 |
| 22843 | ENSG0000( | 2.15E-06 | 1.20E-07 | -5.29308 | 0.34952 | -1.85004 | 34.69475 | PPM1E |
| 1466 | ENSG0000( | 2.76E-22 | 2.07E-24 | -10.1959 | 0.13688 | -1.39561 | 212.0878 | CSRP2 |
| 254295 | ENSG0000( | 1.73E-08 | 6.53E-10 | -6.17697 | 0.315979 | -1.95179 | 40.89261 | PHYHD1 |
| 1139 | ENSG0000( | 0.026888 | 0.005436 | -2.77999 | 0.414569 | -1.1525 | 27.22142 | CHRNA7 |
| 285598 | ENSG0000( | 2.36E-07 | 1.11E-08 | -5.71253 | 0.177897 | -1.01624 | 117.2176 | ARL10 |
| 317671 | ENSG0000( | 0.015103 | 0.00271 | -2.99881 | 0.372059 | -1.11573 | 26.05967 | RFESD |
| 374403 | ENSG0000( | 0.221332 | 0.081642 | 1.741239 | 0.579308 | 1.008714 | 11.41575 | TBC1D10C |
| 2877 | ENSG0000( | 0.077323 | 0.020346 | 2.319904 | 0.52269 | 1.212591 | 13.79158 | GPX2 |
| 664 | ENSG0000( | 9.17E-05 | 7.50E-06 | -4.47904 | 0.282332 | -1.26458 | 49.37997 | BNIP3 |
| 220441 | ENSG0000( | 0.016841 | 0.003089 | -2.95874 | 0.95511 | -2.82592 | 6.014386 | RNF152 |
| 10409 | ENSG0000( | 0.00947 | 0.00156 | -3.16326 | 0.554195 | -1.75306 | 12.85258 | BASP1 |
| 8325 | ENSG0000( | 1.03E-05 | 6.70E-07 | -4.96987 | 0.295676 | -1.46947 | 46.26053 | FZD8 |
| 203111 | ENSG0000( | 0.001411 | 0.000166 | -3.76568 | 0.73742 | -2.77689 | 10.05306 | ERICH5 |
| 122970 | ENSG0000( | 0.01963 | 0.003709 | 2.901882 | 0.367965 | 1.06779 | 28.12695 | ACOT4 |
| 56475 | ENSG0000( | 0.000373 | 3.65E-05 | -4.12874 | 0.565431 | -2.33452 | 14.65297 | RPRM |
| 79605 | ENSG0000( | 0.003757 | 0.000527 | -3.46654 | 0.456512 | -1.58252 | 18.35331 | PGBD5 |
| 222183 | ENSG0000( | 0.012379 | 0.00214 | 3.070105 | 0.41781 | 1.28272 | 22.08321 | SRRM3 |
| 56975 | ENSG0000( | 1.28E-10 | 3.57E-12 | -6.95343 | 0.154354 | -1.07329 | 152.0034 | FAM20C |
| 85460 | ENSG0000( | 0.00041 | 4.07E-05 | -4.10351 | 0.303572 | -1.24571 | 40.28579 | ZNF518B |
| 340152 | ENSG0000( | 0.113385 | 0.033426 | 2.126927 | 0.659174 | 1.402016 | 8.976657 | ZC3H12D |
| 441151 | ENSG0000( | 0.368613 | 0.169563 | 1.37361 | 0.96448 | 1.32482 | 4.632466 | TMEM151I |
| 2731 | ENSG0000( | 8.76E-19 | 8.71E-21 | -9.35062 | 0.183067 | -1.71179 | 117.6921 | GLDC |
| 258 | ENSG0000( | 0.000124 | 1.06E-05 | -4.40528 | 0.680622 | -2.99833 | 11.67338 | AMBN |
| 767 | ENSG0000( | 5.63E-16 | 7.30E-18 | -8.61005 | 0.189131 | -1.62843 | 110.9812 | CA8 |
| 2066 | ENSG0000( | 0.033823 | 0.007219 | -2.68656 | 0.81309 | -2.18442 | 6.428527 | ERBB4 |
| 4094 | ENSG0000( | 1.53E-21 | 1.23E-23 | -10.0215 | 0.180844 | -1.81233 | 122.4481 | MAF |

| | | | | | | | | |
|---|---|---|---|---|---|---|---|---|
| 115207 | ENSG0000( | 5.16E-42 | 1.07E-44 | -14.0266 | 0.118617 | -1.6638 | 286.4583 | KCTD12 |
| 23302 | ENSG0000( | 0.00013 | 1.11E-05 | -4.39432 | 0.317377 | -1.39466 | 38.549 | WSCD1 |
| 388394 | ENSG0000( | 0.024487 | 0.004863 | -2.81598 | 0.856947 | -2.41314 | 6.307788 | RPRML |
| 51268 | ENSG0000( | 0.546666 | 0.309211 | 1.016879 | 1.206822 | 1.227193 | 2.54058 | PIPOX |
| 10692 | ENSG0000( | 0.619316 | 0.380155 | 0.87761 | 1.151514 | 1.01058 | 2.820425 | RRH |
| 2619 | ENSG0000( | 0.000124 | 1.05E-05 | -4.40679 | 0.330271 | -1.45544 | 42.77236 | GAS1 |
| 168620 | ENSG0000( | 0.006986 | 0.001094 | 3.265233 | 0.34943 | 1.14097 | 31.55995 | BHLHA15 |
| 8334 | ENSG0000( | 1.65E-12 | 3.47E-14 | -7.5794 | 0.150696 | -1.14219 | 159.6813 | H2AC6 |
| 284129 | ENSG0000( | 0.000391 | 3.86E-05 | -4.1155 | 0.338768 | -1.3942 | 32.92234 | SLC26A11 |
| 124842 | ENSG0000( | 2.58E-28 | 1.09E-30 | 11.5166 | 0.103856 | 1.19607 | 362.3534 | TMEM132I |
| 2837 | ENSG0000( | 0.014568 | 0.002596 | -3.01192 | 0.497423 | -1.4982 | 15.37913 | UTS2R |
| 168544 | ENSG0000( | 0.015959 | 0.002891 | -2.97909 | 0.889027 | -2.64849 | 6.324114 | ZNF467 |
| 84282 | ENSG0000( | 0.000394 | 3.89E-05 | -4.11378 | 0.411723 | -1.69374 | 22.87209 | RNF135 |
| 6448 | ENSG0000( | 1.92E-08 | 7.41E-10 | -6.15711 | 0.245157 | -1.50946 | 67.00545 | SGSH |
| 284451 | ENSG0000( | 0.26268 | 0.10323 | 1.62939 | 1.474173 | 2.402004 | 2.352866 | ODF3L2 |
| 55228 | ENSG0000( | 4.24E-14 | 6.56E-16 | -8.07847 | 0.205513 | -1.66023 | 92.75995 | PNMA8A |
| 8633 | ENSG0000( | 7.80E-07 | 4.01E-08 | -5.49038 | 0.335 | -1.83928 | 36.4804 | UNC5C |
| 23641 | ENSG0000( | 5.19E-10 | 1.61E-11 | -6.73735 | 0.268296 | -1.8076 | 54.80627 | LDOC1 |
| 55137 | ENSG0000( | 0.000524 | 5.42E-05 | -4.03694 | 0.391924 | -1.58218 | 25.1541 | FIGN |
| 284359 | ENSG0000( | 0.627167 | 0.388289 | 0.862724 | 1.263641 | 1.090174 | 2.343122 | IZUMO1 |
| 402415 | ENSG0000( | 0.61633 | 0.377857 | 0.881852 | 1.227536 | 1.082504 | 2.384171 | XKRX |
| 201305 | ENSG0000( | 0.515619 | 0.280353 | 1.079528 | 1.370691 | 1.479699 | 2.144806 | SPNS3 |
| 6304 | ENSG0000( | 2.50E-05 | 1.77E-06 | -4.7783 | 0.282641 | -1.35054 | 48.45843 | SATB1 |
| 6546 | ENSG0000( | 6.51E-06 | 4.07E-07 | -5.06568 | 0.40134 | -2.03306 | 25.97293 | SLC8A1 |
| 18 | ENSG0000( | 1.97E-49 | 2.98E-52 | 15.2113 | 0.0793 | 1.206253 | 671.3862 | ABAT |
| 1482 | ENSG0000( | 5.18E-13 | 9.84E-15 | -7.74135 | 0.252539 | -1.95499 | 66.0256 | NKX2-5 |
| 10082 | ENSG0000( | 7.72E-07 | 3.96E-08 | -5.49277 | 0.344636 | -1.89301 | 34.17748 | GPC6 |
| 150221 | ENSG0000( | 0.016397 | 0.002984 | -2.96938 | 0.459551 | -1.36458 | 17.06312 | RIMBP3C |
| 147372 | ENSG0000( | 0.023178 | 0.004552 | -2.83715 | 0.960267 | -2.72442 | 5.429316 | CCBE1 |
| 5367 | ENSG0000( | 0.010936 | 0.001854 | -3.11271 | 0.912584 | -2.84061 | 6.066779 | PMCH |
| 29933 | ENSG0000( | 0.205236 | 0.073393 | 1.790382 | 1.409229 | 2.523057 | 2.566058 | GPR132 |
| 54039 | ENSG0000( | 4.67E-10 | 1.44E-11 | -6.7539 | 0.180882 | -1.22166 | 109.6386 | PCBP3 |
| 1232 | ENSG0000( | 0.380797 | 0.177699 | 1.347875 | 0.836431 | 1.127404 | 5.457211 | CCR3 |
| 90187 | ENSG0000( | 1.53E-05 | 1.03E-06 | -4.8859 | 0.357027 | -1.7444 | 32.21376 | EMILIN3 |
| 256933 | ENSG0000( | 0.3629 | 0.165693 | 1.386175 | 1.545324 | 2.14209 | 2.073634 | NPB |
| 6899 | ENSG0000( | 5.43E-15 | 7.86E-17 | -8.33338 | 0.256798 | -2.14 | 64.50615 | TBX1 |
| 55803 | ENSG0000( | 0.008512 | 0.001385 | -3.19774 | 0.383932 | -1.22772 | 25.82148 | ADAP2 |
| 5101 | ENSG0000( | 1.13E-22 | 8.16E-25 | -10.2858 | 0.186177 | -1.91498 | 119.1139 | PCDH9 |
| 5587 | ENSG0000( | 0.001733 | 0.000211 | -3.70606 | 0.385536 | -1.42882 | 25.5542 | PRKD1 |
| 5454 | ENSG0000( | 6.56E-09 | 2.32E-10 | -6.33822 | 0.192971 | -1.22309 | 110.4965 | POU3F2 |
| 1850 | ENSG0000( | 0.001646 | 0.000198 | -3.72117 | 0.5744 | -2.13744 | 12.59895 | DUSP8 |
| 642273 | ENSG0000( | 0.115073 | 0.034134 | 2.118489 | 0.729488 | 1.545413 | 7.419068 | FAM110C |
| 11274 | ENSG0000( | 4.53E-05 | 3.44E-06 | -4.64289 | 0.358733 | -1.66556 | 32.9899 | USP18 |
| 6092 | ENSG0000( | 0.00136 | 0.000159 | -3.7761 | 0.610246 | -2.30435 | 12.40142 | ROBO2 |
| 27124 | ENSG0000( | 0.002731 | 0.000357 | -3.56963 | 0.351526 | -1.25482 | 29.71071 | INPP5J |
| 54033 | ENSG0000( | 0.021875 | 0.004242 | -2.85956 | 0.588334 | -1.68238 | 10.90166 | RBM11 |
| 93349 | ENSG0000( | 0.011906 | 0.002048 | 3.083135 | 0.361323 | 1.114006 | 29.27174 | SP140L |
| 25840 | ENSG0000( | 0.000105 | 8.77E-06 | -4.44541 | 0.291521 | -1.29593 | 47.5801 | METTL7A |
| 285513 | ENSG0000( | 1.68E-05 | 1.14E-06 | -4.86569 | 0.399354 | -1.94313 | 25.27749 | GPRIN3 |
| 5592 | ENSG0000( | 0.002835 | 0.000375 | -3.55684 | 0.43571 | -1.54975 | 20.07525 | PRKG1 |

| | | | | | | | | |
|---|---|---|---|---|---|---|---|---|
| 5087 | ENSG0000( | 2.39E-08 | 9.38E-10 | -6.11954 | 0.246998 | -1.51152 | 61.37506 | PBX1 |
| 23532 | ENSG0000( | 8.33E-13 | 1.63E-14 | -7.67689 | 0.239203 | -1.83634 | 73.02053 | PRAME |
| 56479 | ENSG0000( | 0.004012 | 0.00057 | -3.44561 | 0.453204 | -1.56157 | 18.19803 | KCNQ5 |
| 1E+08 | ENSG0000( | 0.404019 | 0.193798 | 1.299426 | 1.05751 | 1.374155 | 3.403251 | HDHD5-AS: |
| 374899 | ENSG0000( | 0.04836 | 0.011299 | -2.53332 | 0.53479 | -1.3548 | 12.95484 | ZNF829 |
| 84856 | ENSG0000( | 0.016072 | 0.002915 | -2.97661 | 0.430017 | -1.27999 | 20.76688 | LINC00839 |
| 8660 | ENSG0000( | 4.48E-34 | 1.38E-36 | -12.6335 | 0.109459 | -1.38285 | 321.2319 | IRS2 |
| 64757 | ENSG0000( | 1.64E-10 | 4.65E-12 | -6.91598 | 0.21889 | -1.51384 | 78.83948 | 1-Mar |
| 4675 | ENSG0000( | 0.020621 | 0.003946 | -2.88246 | 0.483102 | -1.39252 | 16.22498 | NAP1L3 |
| 27445 | ENSG0000( | 0.000402 | 3.99E-05 | -4.10827 | 0.383657 | -1.57617 | 28.54817 | PCLO |
| 84109 | ENSG0000( | 0.0031 | 0.00042 | -3.52709 | 0.600779 | -2.119 | 13.05487 | QRFPR |
| 2248 | ENSG0000( | 1.13E-13 | 1.96E-15 | 7.944131 | 0.1353 | 1.074844 | 219.5215 | FGF3 |
| 1282 | ENSG0000( | 5.04E-06 | 3.06E-07 | -5.11941 | 0.34851 | -1.78417 | 33.78241 | COL4A1 |
| 148398 | ENSG0000( | 2.23E-23 | 1.46E-25 | -10.4504 | 0.196145 | -2.0498 | 118.7192 | SAMD11 |
| 389421 | ENSG0000( | 3.72E-17 | 4.43E-19 | -8.92576 | 0.165518 | -1.47738 | 137.9327 | LIN28B |
| 3006 | ENSG0000( | 5.81E-14 | 9.25E-16 | -8.03644 | 0.137116 | -1.10192 | 199.2955 | H1-2 |
| 7373 | ENSG0000( | 0.000628 | 6.64E-05 | -3.98894 | 0.851692 | -3.39735 | 8.574025 | COL14A1 |
| 138649 | ENSG0000( | 2.27E-05 | 1.59E-06 | -4.79938 | 0.333131 | -1.59882 | 42.46445 | ANKRD19P |
| 5133 | ENSG0000( | 0.246367 | 0.09469 | 1.671162 | 0.940549 | 1.57181 | 4.527676 | PDCD1 |
| 84570 | ENSG0000( | 0.000259 | 2.42E-05 | -4.22207 | 0.426518 | -1.80079 | 21.7187 | COL25A1 |
| 344901 | ENSG0000( | 0.000448 | 4.52E-05 | -4.07922 | 0.437019 | -1.7827 | 20.50544 | OSTN |
| 389206 | ENSG0000( | 7.99E-07 | 4.12E-08 | -5.48581 | 0.25567 | -1.40256 | 56.80266 | BEND4 |
| 120892 | ENSG0000( | 0.027635 | 0.005629 | -2.76865 | 0.67262 | -1.86225 | 8.703902 | LRRK2 |
| 401494 | ENSG0000( | 0.635876 | 0.397523 | 0.846054 | 1.294913 | 1.095567 | 2.407596 | HACD4 |
| 257177 | ENSG0000( | 0.449178 | 0.227133 | 1.207778 | 3.579642 | 4.323413 | 2.094704 | CFAP126 |
| 80320 | ENSG0000( | 0.000257 | 2.40E-05 | 4.224036 | 0.248005 | 1.047582 | 64.38897 | SP6 |
| 1364 | ENSG0000( | 6.10E-88 | 2.30E-91 | 20.27134 | 0.05436 | 1.101954 | 1663.72 | CLDN4 |
| 79633 | ENSG0000( | 0.002332 | 0.000297 | -3.61762 | 0.465631 | -1.68447 | 17.94374 | FAT4 |
| 148213 | ENSG0000( | 0.020084 | 0.00382 | -2.89269 | 0.505691 | -1.46281 | 16.94549 | ZNF681 |
| 94027 | ENSG0000( | 0.38244 | 0.179091 | 1.343558 | 1.025207 | 1.377425 | 4.042554 | CGB7 |
| 55190 | ENSG0000( | 0.000131 | 1.12E-05 | -4.39281 | 0.272709 | -1.19796 | 51.52025 | NUDT11 |
| 222553 | ENSG0000( | 3.81E-08 | 1.55E-09 | -6.039 | 0.27508 | -1.66121 | 53.30879 | SLC35F1 |
| 124056 | ENSG0000( | 0.113399 | 0.033445 | 2.126706 | 0.857433 | 1.823508 | 6.86267 | NOXO1 |
| 440348 | ENSG0000( | 0.028444 | 0.00582 | -2.75773 | 1.302855 | -3.59292 | 3.951683 | NPIPB15 |
| 4311 | ENSG0000( | 0.002839 | 0.000376 | -3.55621 | 0.487738 | -1.7345 | 16.23515 | MME |
| 6925 | ENSG0000( | 7.14E-12 | 1.67E-13 | -7.37269 | 0.263291 | -1.94117 | 60.86975 | TCF4 |
| 1612 | ENSG0000( | 8.44E-09 | 3.04E-10 | -6.29692 | 0.225436 | -1.41955 | 81.41484 | DAPK1 |
| 7088 | ENSG0000( | 3.68E-12 | 8.25E-14 | -7.46633 | 0.193792 | -1.44691 | 99.66506 | TLE1 |
| 837 | ENSG0000( | 0.090019 | 0.024736 | 2.245493 | 0.949932 | 2.133066 | 5.058536 | CASP4 |
| 51703 | ENSG0000( | 4.93E-16 | 6.33E-18 | 8.626403 | 0.125909 | 1.086143 | 261.7755 | ACSL5 |
| 6799 | ENSG0000( | 0.26045 | 0.102118 | 1.63467 | 1.121881 | 1.833904 | 3.464075 | SULT1A2 |
| 64499 | ENSG0000( | 0.027507 | 0.005589 | 2.77097 | 0.681317 | 1.88791 | 9.720711 | TPSB2 |
| 1E+08 | ENSG0000( | 0.003852 | 0.000544 | -3.45817 | 0.679112 | -2.34848 | 9.769123 | RAMP2-AS: |
| 79027 | ENSG0000( | 2.73E-13 | 4.95E-15 | -7.8282 | 0.207222 | -1.62217 | 92.15252 | ZNF655 |
| 6693 | ENSG0000( | 0.001776 | 0.000216 | 3.699184 | 0.37642 | 1.392447 | 30.13613 | SPN |
| 1288 | ENSG0000( | 0.004523 | 0.000656 | -3.40744 | 0.418113 | -1.4247 | 21.62369 | COL4A6 |
| 5167 | ENSG0000( | 1.18E-05 | 7.80E-07 | -4.94042 | 0.213504 | -1.0548 | 78.93643 | ENPP1 |
| 645811 | ENSG0000( | 0.519937 | 0.284277 | 1.07076 | 1.111364 | 1.190005 | 3.132962 | CCDC154 |
| 388585 | ENSG0000( | 0.032495 | 0.006883 | -2.70247 | 1.03042 | -2.78468 | 4.563023 | HES5 |
| 54898 | ENSG0000( | 3.65E-10 | 1.10E-11 | -6.79229 | 0.315401 | -2.14229 | 45.58009 | ELOVL2 |

| | | | | | | | | |
|---|---|---|---|---|---|---|---|---|
| 79366 | ENSG0000( | 3.62E-09 | 1.24E-10 | -6.43398 | 0.226733 | -1.45879 | 74.59232 | HMGN5 |
| 777 | ENSG0000( | 4.26E-20 | 3.86E-22 | 9.674661 | 0.224183 | 2.168897 | 92.64884 | CACNA1E |
| 4914 | ENSG0000( | 0.2686 | 0.106826 | 1.612624 | 1.106902 | 1.785018 | 3.389355 | NTRK1 |
| 374900 | ENSG0000( | 0.008429 | 0.00137 | -3.20089 | 0.648486 | -2.07573 | 9.888267 | ZNF568 |
| 286319 | ENSG0000( | 3.50E-06 | 2.06E-07 | -5.19365 | 0.386 | -2.00475 | 31.95779 | TUSC1 |
| 9060 | ENSG0000( | 5.80E-06 | 3.57E-07 | -5.09036 | 0.224506 | -1.14282 | 72.27027 | PAPSS2 |
| 64093 | ENSG0000( | 0.007451 | 0.001184 | -3.24275 | 0.390957 | -1.26778 | 25.03632 | SMOC1 |
| 140606 | ENSG0000( | 0.003092 | 0.000419 | -3.5281 | 0.289154 | -1.02017 | 42.61632 | SELENOM |
| 57713 | ENSG0000( | 4.81E-09 | 1.67E-10 | -6.38879 | 0.271147 | -1.7323 | 58.80359 | SFMBT2 |
| 5455 | ENSG0000( | 0.005832 | 0.000885 | -3.3248 | 0.301306 | -1.00178 | 39.45405 | POU3F3 |
| 221002 | ENSG0000( | 0.002211 | 0.00028 | -3.63292 | 0.323491 | -1.17522 | 35.19934 | RASGEF1A |
| 57692 | ENSG0000( | 0.002797 | 0.000368 | -3.5619 | 0.56744 | -2.02117 | 13.74172 | MAGEE1 |
| 1756 | ENSG0000( | 3.95E-05 | 2.94E-06 | -4.67489 | 0.325485 | -1.52161 | 35.16351 | DMD |
| 84546 | ENSG0000( | 0.021155 | 0.004078 | -2.87204 | 1.632918 | -4.6898 | 2.07586 | SNORD35B |
| 594839 | ENSG0000( | 0.452783 | 0.229744 | 1.201018 | 0.992113 | 1.191546 | 3.753832 | SNORA33 |
| 389432 | ENSG0000( | 0.000273 | 2.57E-05 | -4.20883 | 0.400932 | -1.68745 | 24.14211 | SAMD5 |
| 286411 | ENSG0000( | 0.000218 | 1.98E-05 | -4.26691 | 0.46003 | -1.96291 | 20.43663 | LINC00632 |
| 26071 | ENSG0000( | 1.37E-12 | 2.80E-14 | -7.60735 | 0.233787 | -1.7785 | 73.39073 | RTL8A |
| 440590 | ENSG0000( | 8.90E-06 | 5.70E-07 | -5.00127 | 0.506167 | -2.53148 | 17.77125 | ZYG11A |
| 3111 | ENSG0000( | 0.001332 | 0.000155 | -3.78253 | 0.535235 | -2.02454 | 14.58812 | HLA-DOA |
| 1290 | ENSG0000( | 3.87E-08 | 1.58E-09 | -6.03626 | 0.304675 | -1.83909 | 47.1406 | COL5A2 |
| 643008 | ENSG0000( | 0.389826 | 0.184267 | 1.327731 | 0.983659 | 1.306034 | 3.966812 | SMIM5 |
| 143903 | ENSG0000( | 0.003417 | 0.000471 | -3.4968 | 0.495675 | -1.73328 | 16.33083 | LAYN |
| 80740 | ENSG0000( | 0.464793 | 0.238714 | 1.178208 | 1.375193 | 1.620263 | 2.280453 | LY6G6C |
| 11025 | ENSG0000( | 0.444314 | 0.223583 | 1.217056 | 1.028766 | 1.252067 | 3.885283 | LILRB3 |
| 1.03E+08 | ENSG0000( | 0.444314 | 0.223583 | 1.217056 | 1.028766 | 1.252067 | 3.885283 | LOC102725 |
| 1.08E+08 | ENSG0000( | 0.444314 | 0.223583 | 1.217056 | 1.028766 | 1.252067 | 3.885283 | LOC107987 |
| 1.08E+08 | ENSG0000( | 0.444314 | 0.223583 | 1.217056 | 1.028766 | 1.252067 | 3.885283 | LOC107987 |
| 441054 | ENSG0000( | 0.131091 | 0.040296 | 2.050702 | 0.546559 | 1.12083 | 13.28677 | C4orf47 |
| 388135 | ENSG0000( | 0.001285 | 0.000149 | -3.7935 | 0.510095 | -1.93505 | 15.5299 | INSYN1 |
| 3887 | ENSG0000( | 5.58E-23 | 3.78E-25 | -10.3596 | 0.107565 | -1.11433 | 319.2986 | KRT81 |
| 8110 | ENSG0000( | 0.028717 | 0.005887 | -2.754 | 0.824573 | -2.27087 | 6.599239 | DPF3 |
| 10215 | ENSG0000( | 0.008361 | 0.001357 | -3.20369 | 0.70794 | -2.26802 | 8.440081 | OLIG2 |
| 23617 | ENSG0000( | 0.419056 | 0.204412 | 1.269081 | 3.650364 | 4.632607 | 2.596284 | TSSK2 |
| 389136 | ENSG0000( | 0.001784 | 0.000217 | -3.69794 | 0.454841 | -1.68198 | 19.70698 | VGLL3 |
| 654319 | ENSG0000( | 0.483806 | 0.255115 | 1.138013 | 0.995872 | 1.133315 | 3.642633 | SNORA5A |
| 402317 | ENSG0000( | 0.519383 | 0.283778 | 1.071871 | 1.401746 | 1.50249 | 2.147487 | OR2A42 |
| 339803 | ENSG0000( | 0.001905 | 0.000234 | -3.67867 | 0.480346 | -1.76704 | 17.13031 | LOC339803 |
| 641311 | ENSG0000( | 0.51755 | 0.281897 | 1.076067 | 1.364171 | 1.467939 | 2.12572 | RPL31P11 |
| 1396 | ENSG0000( | 0.306556 | 0.12901 | 1.518017 | 0.682532 | 1.036096 | 50.68425 | CRIP1 |
| 4056 | ENSG0000( | 0.379147 | 0.176528 | 1.351523 | 1.280019 | 1.729976 | 2.45325 | LTC4S |
| 2946 | ENSG0000( | 2.87E-06 | 1.65E-07 | -5.23461 | 0.388929 | -2.03589 | 27.57861 | GSTM2 |
| 728643 | ENSG0000( | 0.359422 | 0.163133 | 1.394612 | 1.342952 | 1.872896 | 2.640334 | HNRNPA1P |
| 445329 | ENSG0000( | 0.063233 | 0.015749 | 2.414691 | 0.515835 | 1.245582 | 20.35708 | SULT1A4 |
| 344167 | ENSG0000( | 0.04531 | 0.010398 | -2.5623 | 0.663401 | -1.69983 | 8.679193 | FOXI3 |
| 57412 | ENSG0000( | 7.13E-06 | 4.47E-07 | -5.04781 | 0.367041 | -1.85275 | 31.12605 | AS3MT |
| 392862 | ENSG0000( | 0.049745 | 0.011704 | -2.52095 | 1.170342 | -2.95038 | 3.834929 | GRID2IP |
| 3166 | ENSG0000( | 7.54E-05 | 6.01E-06 | -4.52599 | 0.430662 | -1.94917 | 22.93148 | HMX1 |
| 6248 | ENSG0000( | 0.680057 | 0.44899 | 0.7571 | 1.529919 | 1.158302 | 72.75414 | RSC1A1 |
| 284498 | ENSG0000( | 0.001549 | 0.000185 | -3.7392 | 1.491406 | -5.57667 | 3.857681 | C1orf167 |

| | | | | | | | | |
|---|---|---|---|---|---|---|---|---|
| 677847 | ENSG00000( | 0.359478 | 0.163193 | 1.394412 | 0.944301 | 1.316744 | 4.686569 | SNORA81 |
| 1E+08 | ENSG00000( | 0.339127 | 0.149803 | 1.440227 | 3.736863 | 5.381931 | 4.356412 | MIR1184-1 |
| 257000 | ENSG00000( | 0.16045 | 0.052935 | 1.935451 | 0.60652 | 1.17389 | 10.52027 | TINCR |
| 1E+08 | ENSG00000( | 7.39E-14 | 1.22E-15 | -8.00236 | 0.245586 | -1.96527 | 73.3895 | LINC02593 |
| 399821 | ENSG00000( | 0.350054 | 0.156745 | 1.416106 | 0.968063 | 1.370879 | 4.063299 | EDRF1-DT |
| 721 | ENSG00000( | 0.163529 | 0.054331 | 1.924186 | 0.978892 | 1.883571 | 4.458197 | C4B |
| 26777 | ENSG00000( | 0.215401 | 0.078505 | 1.759426 | 1.231431 | 2.166612 | 3.11489 | SNORA71A |
| 1.02E+08 | ENSG00000( | 0.335087 | 0.147145 | 1.449689 | 1.580332 | 2.29099 | 2.201061 | LOC101928 |
| 1.08E+08 | ENSG00000( | 0.51995 | 0.284317 | 1.070672 | 1.24448 | 1.33243 | 2.670635 | LINC01772 |
| 65072 | ENSG00000( | 0.321848 | 0.138444 | 1.48161 | 1.258296 | 1.864304 | 2.653388 | CFLAR-AS1 |
| 388780 | ENSG00000( | 0.056434 | 0.013665 | 2.465961 | 0.706003 | 1.740976 | 8.224097 | LOC38878( |
| 1.02E+08 | ENSG00000( | 0.448989 | 0.227009 | 1.208099 | 0.932516 | 1.126572 | 4.24006 | LOC101929 |
| 57212 | ENSG00000( | 5.00E-14 | 7.86E-16 | -8.0563 | 0.233391 | -1.88026 | 75.98309 | TP73-AS1 |
| 4050 | ENSG00000( | 0.140532 | 0.044119 | 2.012963 | 0.964033 | 1.940563 | 4.594074 | LTB |
| 1.01E+08 | ENSG00000( | 0.005014 | 0.00074 | -3.37419 | 2.492048 | -8.40864 | 27.35595 | CERS6-AS1 |
| 1.01E+08 | ENSG00000( | 0.001146 | 0.000131 | -3.82531 | 0.642064 | -2.45609 | 11.4908 | SOX21-AS1 |
| 283008 | ENSG00000( | 0.637892 | 0.399786 | 0.842003 | 2.711236 | 2.28287 | 3.454154 | NUTM2E |
| 400946 | ENSG00000( | 0.677912 | 0.446524 | 0.761223 | 1.331461 | 1.013539 | 2.254183 | LINC00954 |
| 401093 | ENSG00000( | 0.026356 | 0.005309 | -2.78768 | 0.958096 | -2.67087 | 6.681015 | MBNL1-AS1 |
| 7503 | ENSG00000( | 2.23E-27 | 9.96E-30 | -11.3242 | 0.163765 | -1.85451 | 4961.538 | XIST |
| 441459 | ENSG00000( | 0.002062 | 0.000258 | -3.65447 | 0.398471 | -1.4562 | 30.51671 | ANKRD18B |
| 439931 | ENSG00000( | 0.006413 | 0.000992 | -3.29269 | 0.324225 | -1.06757 | 35.26276 | THAP7-AS1 |
| 1.02E+08 | ENSG00000( | 0.018702 | 0.003495 | 2.920454 | 0.373359 | 1.090378 | 27.81925 | SPRY4-AS1 |
| 3113 | ENSG00000( | 0.000218 | 1.98E-05 | -4.26674 | 0.494871 | -2.11149 | 17.10089 | HLA-DPA1 |
| 1.02E+08 | ENSG00000( | 0.505967 | 0.272532 | 1.09725 | 0.999571 | 1.09678 | 4.126973 | COA6-AS1 |
| 1.02E+08 | ENSG00000( | 0.102498 | 0.029381 | 2.178336 | 0.54622 | 1.189851 | 12.99641 | LOC101927 |
| 151484 | ENSG00000( | 0.611943 | 0.374205 | 0.888625 | 1.239799 | 1.101717 | 2.386852 | GCSIR |
| 1.01E+08 | ENSG00000( | 0.042016 | 0.009462 | -2.59488 | 0.58789 | -1.52551 | 11.17745 | ZNF687-AS |
| 1.01E+08 | ENSG00000( | 0.006315 | 0.000974 | -3.29795 | 0.530979 | -1.75114 | 14.05481 | LINC00665 |
| 79940 | ENSG00000( | 0.024666 | 0.004901 | -2.81344 | 0.558472 | -1.57123 | 12.15204 | LINC00472 |
| 117581 | ENSG00000( | 0.016678 | 0.003053 | -2.96236 | 0.355001 | -1.05164 | 28.66922 | TWIST2 |
| 440900 | ENSG00000( | 0.577929 | 0.339045 | 0.956054 | 1.07321 | 1.026046 | 3.457682 | LINC01191 |
| 1.01E+08 | ENSG00000( | 2.79E-13 | 5.10E-15 | -7.82441 | 0.248731 | -1.94617 | 65.98056 | MAGI2-AS3 |
| 344 | ENSG00000( | 0.028956 | 0.005947 | 2.750679 | 0.809357 | 2.226281 | 7.43608 | APOC2 |
| 1.01E+08 | ENSG00000( | 0.295316 | 0.122293 | 1.545222 | 0.653177 | 1.009303 | 9.040833 | TM4SF19-A |
| 1.02E+08 | ENSG00000( | 0.214679 | 0.078122 | 1.761688 | 0.758762 | 1.336702 | 6.645011 | ZPLD2P |
| 401491 | ENSG00000( | 0.421638 | 0.20607 | 1.264447 | 0.94773 | 1.198354 | 4.336564 | VLDLR-AS1 |
| 728262 | ENSG00000( | 0.007382 | 0.00117 | -3.24618 | 0.362361 | -1.17629 | 30.31116 | FAM157A |
| 1.02E+08 | ENSG00000( | 0.191731 | 0.067043 | 1.831385 | 0.753465 | 1.379885 | 6.818796 | LINC01844 |
| 643854 | ENSG00000( | 0.680413 | 0.449641 | 0.756014 | 2.567924 | 1.941386 | 2.709336 | CTAGE9 |
| 441094 | ENSG00000( | 2.06E-07 | 9.50E-09 | -5.73939 | 0.270821 | -1.55435 | 51.66957 | NR2F1-AS1 |
| 1.02E+08 | ENSG00000( | 0.425807 | 0.20993 | 1.253759 | 1.096032 | 1.37416 | 3.347703 | LINC01637 |
| 339804 | ENSG00000( | 4.56E-05 | 3.46E-06 | -4.64134 | 0.557475 | -2.58743 | 15.43806 | C2orf74 |
| 339751 | ENSG00000( | 0.050646 | 0.011986 | 2.512558 | 1.295595 | 3.255257 | 3.971586 | MAP3K20-A |
| 1.02E+08 | ENSG00000( | 0.559586 | 0.321132 | 0.992135 | 1.405673 | 1.394617 | 2.105331 | CD81-AS1 |
| 358 | ENSG00000( | 0.546308 | 0.308868 | 1.017601 | 1.009541 | 1.027309 | 4.009194 | AQP1 |
| 632 | ENSG00000( | 0.356842 | 0.161445 | 1.400227 | 1.544608 | 2.162802 | 2.122128 | BGLAP |
| 1.05E+08 | ENSG00000( | 0.437641 | 0.218765 | 1.229818 | 0.840566 | 1.033744 | 5.221972 | LINC02050 |
| 54657 | ENSG00000( | 0.0313 | 0.006557 | -2.71857 | 0.542557 | -1.47498 | 13.05439 | UGT1A4 |
| 91179 | ENSG00000( | 1.64E-11 | 4.04E-13 | -7.25433 | 0.233571 | -1.6944 | 71.7594 | SCARF2 |

| | | | | | | | | |
|---|---|---|---|---|---|---|---|---|
| 728339 | ENSG0000( | 0.307979 | 0.129862 | 1.514647 | 1.468486 | 2.224237 | 2.137931 | FRG1-DT |
| 553158 | ENSG0000( | 0.142211 | 0.044859 | 2.005971 | 1.649374 | 3.308595 | 2.239293 | PRR5-ARH( |
| 1E+08 | ENSG0000( | 0.037751 | 0.008245 | 2.641861 | 1.62429 | 4.291149 | 2.051205 | DDX11L9 |
| 1.02E+08 | ENSG0000( | 0.031204 | 0.006522 | -2.72029 | 0.861509 | -2.34356 | 6.151313 | CASC9 |
| 643401 | ENSG0000( | 4.46E-05 | 3.37E-06 | -4.64691 | 0.413656 | -1.92222 | 25.87868 | PURPL |
| 1.02E+08 | ENSG0000( | 0.03766 | 0.008221 | -2.64287 | 0.643375 | -1.70036 | 9.472571 | LOC10192 |
| 1.02E+08 | ENSG0000( | 0.437195 | 0.218484 | 1.23057 | 1.323577 | 1.628754 | 2.336459 | LINC01303 |
| 6414 | ENSG0000( | 4.09E-07 | 2.01E-08 | -5.61082 | 0.330945 | -1.85687 | 36.82503 | SELENOP |
| 400043 | ENSG0000( | 9.53E-07 | 4.98E-08 | -5.45219 | 0.292793 | -1.59636 | 45.00341 | LINC02381 |
| 162972 | ENSG0000( | 0.001733 | 0.00021 | -3.70624 | 0.465441 | -1.72504 | 18.49941 | ZNF550 |
| 1.01E+08 | ENSG0000( | 0.001966 | 0.000244 | -3.66844 | 0.672655 | -2.46759 | 12.02697 | LOC100507 |
| 1E+08 | ENSG0000( | 0.002347 | 0.000299 | -3.61578 | 0.455473 | -1.64689 | 20.05 | C8orf88 |
| 1.02E+08 | ENSG0000( | 0.360112 | 0.163877 | 1.392151 | 1.302305 | 1.813006 | 2.546368 | LINC01847 |
| 1.07E+08 | ENSG0000( | 0.000153 | 1.33E-05 | 4.354805 | 0.359379 | 1.565024 | 31.53329 | MIR3142H( |
| 1.05E+08 | ENSG0000( | 0.040078 | 0.008917 | 2.615201 | 0.504814 | 1.32019 | 16.72831 | LOC105375 |
| 56109 | ENSG0000( | 0.341482 | 0.15138 | 1.434673 | 0.7918 | 1.135974 | 6.694894 | PCDHGA6 |
| 56105 | ENSG0000( | 0.043444 | 0.009858 | -2.58078 | 0.660743 | -1.70524 | 9.951049 | PCDHGA11 |
| 56103 | ENSG0000( | 0.130294 | 0.039936 | 2.054411 | 0.622926 | 1.279746 | 9.710802 | PCDHGB2 |
| 441381 | ENSG0000( | 0.602455 | 0.363399 | 0.908908 | 1.835376 | 1.668188 | 7.740596 | LRRC24 |
| 132430 | ENSG0000( | 6.19E-06 | 3.85E-07 | -5.07642 | 0.323479 | -1.64211 | 38.74259 | PABPC4L |
| 1.02E+08 | ENSG0000( | 0.648589 | 0.411021 | 0.822099 | 1.285616 | 1.056903 | 2.310754 | LOC101928 |
| 730013 | ENSG0000( | 0.156934 | 0.051419 | 1.947963 | 0.519217 | 1.011415 | 13.78217 | ABCC6P2 |
| 1.01E+08 | ENSG0000( | 0.46505 | 0.238963 | 1.177584 | 0.90394 | 1.064465 | 5.347621 | DENND5B-/ |
| 1.02E+08 | ENSG0000( | 0.606987 | 0.368816 | 0.898694 | 1.147472 | 1.031226 | 2.940465 | LOC101928 |
| 90649 | ENSG0000( | 0.098854 | 0.027973 | 2.197665 | 0.556335 | 1.222637 | 13.73347 | ZNF486 |
| 27164 | ENSG0000( | 0.000401 | 3.97E-05 | -4.10918 | 0.488671 | -2.00804 | 16.80759 | SALL3 |
| 1.05E+08 | ENSG0000( | 0.279848 | 0.113091 | 1.584451 | 0.893599 | 1.415864 | 5.46619 | DDN-AS1 |
| 1.01E+08 | ENSG0000( | 0.208726 | 0.075324 | 1.778489 | 0.8102 | 1.440932 | 34.0199 | NDUFC2-K( |
| 643664 | ENSG0000( | 0.51238 | 0.27734 | 1.086314 | 1.398156 | 1.518836 | 2.1861 | SLC35G6 |
| 1E+08 | ENSG0000( | 0.08621 | 0.02337 | 2.267334 | 0.45916 | 1.041069 | 23.65538 | USP3-AS1 |
| 23732 | ENSG0000( | 0.030609 | 0.006373 | -2.72792 | 0.890906 | -2.43032 | 5.60871 | FRRS1L |
| 1.05E+08 | ENSG0000( | 0.436384 | 0.21789 | 1.232158 | 1.12191 | 1.382371 | 3.451745 | LOC105371 |
| 84210 | ENSG0000( | 0.00364 | 0.000508 | -3.47671 | 0.923996 | -3.21247 | 7.384811 | ANKRD20A |
| 1.05E+08 | ENSG0000( | 0.434991 | 0.216784 | 1.235125 | 1.079046 | 1.332757 | 3.36838 | LOC105371 |
| 1.01E+08 | ENSG0000( | 1.14E-09 | 3.70E-11 | -6.61547 | 0.313677 | -2.07512 | 45.68164 | TMEM178I |
| 83481 | ENSG0000( | 0.00018 | 1.60E-05 | -4.31392 | 0.30498 | -1.31566 | 41.84567 | EPPK1 |
| 145788 | ENSG0000( | 0.310015 | 0.131325 | 1.508896 | 1.50169 | 2.265894 | 2.205103 | C15orf65 |
| 10418 | ENSG0000( | 1.12E-06 | 5.90E-08 | -5.42172 | 0.416489 | -2.25809 | 25.28194 | SPON1 |
| 1.1E+08 | ENSG0000( | 0.470229 | 0.243342 | 1.166675 | 1.008802 | 1.176944 | 3.691016 | SNORA77B |
| 1.04E+08 | ENSG0000( | 0.582603 | 0.343762 | 0.946758 | 2.378757 | 2.252107 | 48.4364 | MIR3648-2 |
| 1.03E+08 | ENSG0000( | 0.037768 | 0.008251 | 2.641611 | 1.65649 | 4.375802 | 2.169671 | LOC102724 |
| 1.01E+08 | ENSG0000( | 0.250483 | 0.096823 | 1.660453 | 0.863623 | 1.434005 | 7.58369 | RNF157-AS |
| 1.02E+08 | ENSG0000( | 0.601796 | 0.362471 | 0.910667 | 1.178308 | 1.073046 | 2.968264 | LINC01775 |
| 200298 | ENSG0000( | 0.652443 | 0.416542 | 0.812435 | 1.266612 | 1.02904 | 2.297912 | LINC00528 |
| 1.04E+08 | ENSG0000( | 0.00129 | 0.000149 | -3.79206 | 0.712835 | -2.70311 | 9.79185 | LINC01224 |
| 8370 | ENSG0000( | 7.39E-06 | 4.64E-07 | -5.04049 | 0.228365 | -1.15107 | 80.98994 | H4C14 |
| 441294 | ENSG0000( | 0.385693 | 0.181468 | 1.336249 | 1.02565 | 1.370524 | 4.694647 | CTAGE15 |
| 340307 | ENSG0000( | 0.004081 | 0.000581 | -3.44042 | 1.543735 | -5.3111 | 3.189326 | CTAGE6 |
| 8447 | ENSG0000( | 0.094645 | 0.026395 | 2.220354 | 0.967386 | 2.14794 | 5.107857 | DOC2B |
| 1.01E+08 | ENSG0000( | 0.570801 | 0.331591 | 0.970915 | 1.250856 | 1.214474 | 2.487925 | CNNM3-DT |

| | | | | | | | | |
|---|---|---|---|---|---|---|---|---|
| 283683 | ENSG00000( | 0.035889 | 0.007757 | -2.66246 | 0.737177 | -1.96271 | 9.104686 | LOC283683 |
| 1.03E+08 | ENSG00000( | 2.81E-26 | 1.40E-28 | -11.0904 | 0.130247 | -1.44449 | 294.5795 | CBSL |
| 1602 | ENSG00000( | 7.20E-10 | 2.26E-11 | -6.6879 | 0.22538 | -1.50732 | 83.53023 | DACH1 |
| 85495 | ENSG00000( | 3.61E-07 | 1.76E-08 | -5.6341 | 0.453678 | -2.55607 | 29.17063 | RPPH1 |
| 4747 | ENSG00000( | 3.36E-08 | 1.35E-09 | -6.06148 | 0.273056 | -1.65512 | 51.82978 | NEFL |
| 401561 | ENSG00000( | 0.474506 | 0.247046 | 1.157554 | 1.207932 | 1.398246 | 2.730414 | LINC01451 |
| 64881 | ENSG00000( | 0.001869 | 0.000229 | -3.68462 | 0.466957 | -1.72056 | 19.0741 | PCDH20 |
| 1.02E+08 | ENSG00000( | 0.431085 | 0.213915 | 1.242872 | 0.871961 | 1.083735 | 4.720043 | PRNCR1 |
| 1E+08 | ENSG00000( | 0.011035 | 0.001873 | -3.10962 | 0.363179 | -1.12935 | 28.29635 | FAM95C |
| 1.03E+08 | ENSG00000( | 8.02E-06 | 5.07E-07 | -5.02369 | 1.311816 | -6.59016 | 7.753363 | TPTEP2-CSI |
| 728392 | ENSG00000( | 0.031482 | 0.006603 | -2.71625 | 0.463991 | -1.26032 | 16.94464 | LOC728392 |
| 1.05E+08 | ENSG00000( | 0.583356 | 0.344353 | 0.945599 | 1.303375 | 1.23247 | 2.506324 | LOC105375 |
| 1.02E+08 | ENSG00000( | 0.025359 | 0.005063 | -2.80298 | 0.526085 | -1.47461 | 14.10544 | LOC101929 |
