## Supplementary Tables 3,4,5 for "Genomic alterations drive brain metastases formation in colorectal cancer: The role of IRS2"

**Table 3:** List of constructs used for transfection

| Company | Catalog number | Construct name |
|---|---|---|
| GeneCopoeia (USA) | GC-EX-Y3534-Lv247 | pReceiver-Lv247-IRS2 |
| GeneCopoeia (USA) | GC-EX-NEG-Lv247 | pReceiver-Lv247-Empty control |
| Sigma (St. Louis, MO, USA) | TRCN0000061257 | shRNA IRS2 |
| Sigma (St. Louis, MO, USA) | TRCN0000364328 | shRNA nonspecific |
| Addgene (Massachusetts, USA) | 8454 | pCMV-VSV-G |
| Addgene (Massachusetts, USA) | 12260 | psPAX2 |
| Upstate Biotechnology (Lake Placid, NY, USA) | 17-285 | pTOPFLASH |
| Upstate Biotechnology (Lake Placid, NY, USA) | 17-285 | pFOPFLASH |

**Table 4:** qPCR Primers list

| | |
|---|---|
| β-actin (Human) | Forward: GCTCAGGAGGAGCAATGATCTT |
| | Reverse: TTGCCGACAGGATGCAGAA |
| IRS2 (Human) | Forward: CCACCATCGTGAAAGAGTGAAGA |
| | Reverse: GCCTTGTTGGTGCCTCATCT |
| DDR2 (Human) | Forward: CCACTATGCAGAGGCTGACA |
| | Reverse: CAGAGATGAACCTCCCCAAA |
| FLT4 (Human) | Forward: GCCATGTACAAGTGTGTGGTCTC |
| | Reverse: ACTTGTAGCTGTCGGCTTGG |
| C-KIT (Human) | Forward: CCGGTCGATTCTAAGTTCTAC |
| | Reverse: GATTGGTGCTCTCTGAAATCTG |
| FGF2 (Human) | Forward: CAAAAACGGGGGCTTCTTCCTG |
| | Reverse: CCATCTTCCTTCATAGCCAGGTAACG |
| CCND2 (Human) | Forward: TCCTGGCCTCCAAACTCAAA |
| | Reverse: AAGTCATGAGGAGTGACAGC |
| LIFR (Human) | Forward: TGTATGTGGTGACAAAGGAAAA |
| | Reverse: TGGATTTGGAATATCAGGGTAGA |
| FOXA1 (Human) | Forward: AGGAACTGTGAAGATGGAAGG |
| | Reverse: ATGTTGCCGCTCGTAGTC |
| NOTCH3 (Human) | Forward: CAAGGGTGAGAGCCTGATGG |
| | Reverse: GAGTCCACTGACGGCAATCC |
| NDUFC2 (Human) | Forward: GGCTTGTCTACATCGGCTTC |
| | Reverse: TGATGGTCCCTCACAGCATA |
| NDUFB8 (Human) | Forward: GCTCCCTGACCGCTCACAGC |
| | Reverse: TGCCAGTGCATCGGTTCACCC |
| NDUFA8 (Human) | Forward: TGTCGCAAACAGCAGGCAAA |
| | Reverse: CTGGGATCCGGTCTTGGTCT |
| COX15 (Human) | Forward: TGGTGTTCCTTACGGCCCTC |
| | Reverse: CCCAGAATCCGGTGATCAAACT |

| | |
|---|---|
| NDUFA12 (Human) | Forward: ACATTCTGGGATGTGGATGG |
| | Reverse: CTAGTGGTAGAATAAGGTAC |
| NDUFB10 (Human) | Forward: TAGAGCGGCAGCACGCAAAG |
| | Reverse: CTGACAGGCTTTGAGCCGATC |
| NDUFA11 (Human) | Forward: AAAGCCTACAGCACCACCAG |
| | Reverse: TGTCCAACCTTAGCCACTCC |
| COX11 (Human) | Forward: CTACGCTGCCGTACCCCTTT |
| | Reverse: TGACCTGCAACTGCTGATCCT |
| UQCRC2 (Human) | Forward: AAAGTTGCCCCGAAGGTTAAA |
| | Reverse: GAGCATAGTTTTCCAGAGAAGCA |

**Table 5:** List of antibodies used for Western Blot

| Company | Catalog number | Target name |
|---|---|---|
| Abcam (Cambridge, UK) | ab134101 | Anti-IRS2 |
| Upstate Biotechnology (Lake Placid, NY, USA) | 06-24 | Anti-IRS1 |
| Cell Signaling (Massachusetts, USA) | #9271 | Anti-Phospho-Akt (Ser473) |
| Cell Signaling (Massachusetts, USA) | #9272 | Anti-Akt |
| Upstate Biotechnology (Lake Placid, NY, USA) | 05-665 | Anti-active β-catenin |
| BD Transduction (Laboratories, Franklin Lakes, NJ, USA) | 610154 | anti-β-catenin |
| Sigma (St. Louis, MO, USA) | A5441 | Anti-β-actin |
